# Unraveling the hexaploid sweetpotato inheritance using ultra-dense multilocus mapping

**DOI:** 10.1101/689638

**Authors:** Marcelo Mollinari, Bode A. Olukolu, Guilherme da S. Pereira, Awais Khan, Dorcus Gemenet, Craig Yencho, Zhao-Bang Zeng

## Abstract

The hexaploid sweetpotato (*Ipomoea batatas* (L.) Lam., 2n = 6x = 90) is an important staple food crop worldwide and has a vital role in alleviating famine in developing countries. Due to its high ploidy level, genetic studies in sweetpotato lag behind major diploid crops significantly. We built an ultra-dense multilocus integrated genetic map and characterized the inheritance system in a sweetpotato full-sib family using our newly implemented software, MAPpoly. The resulting genetic map revealed 96.5% collinearity between *I. batatas* and its diploid relative *I. trifida.* We computed the genotypic probabilities across the whole genome for all individuals in the mapping population and inferred their complete hexaploid haplotypes. We provide evidence that most of the meiotic configurations (73.3%) were resolved in bivalents, although a small portion of multivalent signatures (15.7%), among other inconclusive configurations (11.0%) were also observed. Except for low levels of preferential pairing in linkage group 2, we observed a hexasomic inheritance mechanism in all linkage groups. We propose that the hexasomic-bivalent inheritance promotes stability to the allelic transmission in sweetpotato.

## Introduction

The cultivated hexaploid sweetpotato (*Ipomoea batatas* (L.) Lam., 2n = 6x = 90) is an important staple food crop worldwide with an annual production of ~112.84 tons^1^. It has a vital role in alleviating famine, especially in developing countries in Africa and Southeast Asia^2^. Despite its undeniable social and economic importance, genetic studies in sweetpotato lag behind major diploid crops significantly due to its complex polyploid genome. Polyploids are organisms with more than two chromosome sets. They are grouped into two categories, *allopolyploids* or *autopolyploids*, when these chromosome sets are originated from either different or same species, respectively^3^. While in diploid organisms the study of allelic transmission and genetic linkage are rather straightforward, in polyploids these studies are greatly hindered due to the wide range of meiotic configurations these species undergo^4–6^. Moreover, current linkage analysis methods for complex polyploids (i.e., ploidy level > 4) are mostly based on pairwise (or two-point) marker analyses^7–12^. These methods rely on the assumption that the information in isolated pairs of markers is sufficient to detect recombination events between them accurately. However, in cases of complex polyploids, this is rarely true due to the limited mapping population size and the incomplete information provided by biallelic markers. Here, we present a fully informative multilocus genetic map of a full-sib hexaploid sweetpotato population derived from a cross between the cultivars ‘Beauregard’ and ‘Tanzania’ (BT population) scored with more than 30,000 informative single nucleotide polymorphisms (SNPs) using our newly developed R package called MAPpoly. We also inferred the haplotypes of all individuals in the full-sib population, which provided novel insights into the multivalent formation and preferential pairing in the sweetpotato genome.

Our multilocus analysis considers multiple SNPs simultaneously and propagates their information through the linkage group (LG) to overcome the typical low informativeness of some two-loci combinations. This strategy is fundamentally important for complex polyploid genome analysis since pairs of biallelic markers carry very little information about the recombination process individually^13,14^. Moreover, the signal-to-noise (S/N) ratio in complex polyploid SNP data sets is considerably lower as compared to that in diploids and tetraploids^15^, thus making the genotype calling more prone to errors. The multilocus approach can more appropriately take into account these errors by using the probability distribution of genotypes provided by the genotype calling software^14^. Actually, multilocus methods are essential to use the information of multiple-dose markers to assess complex polyploid inheritance systems adequately.

Several studies attempted to elucidate the polyploidy nature in sweetpotato (allo vs. autopolyploid), including cytological and molecular marker analyses^9,11,16–22^, and more recently sequence-based studies^23–25^. Two polyploidization scenarios were proposed: the first suggests an allopolyploid origin involving the hybridization of two sweetpotato wild diploid relatives, *I. trifida* and *I. triloba*^20^; the second, well supported by the literature, suggests an autopolyploid origin with *I. triloba* having a dual role in the polyploidization process^21, 23–25^. Corroborating this scenario, the polysomic inheritance observed in several molecular marker studies^9,11,19,22^ rules out the strict allopolyploid sweetpotato origin. Nevertheless, none of these studies presented a comprehensive profile of chromosomal pairing for all homology groups across the whole genome nor the potential formation of multivalents at a population level. Solving these missing pieces of information is essential to unravel the precise mode of inheritance in sweetpotato, and consequently, allow an efficient application of molecular techniques in this complex polyploid breeding system. The BT population coupled with high-coverage sequence genotyping has two essential characteristics that enabled high-quality mapping: 1) high and uniform sequence read depth across the genome, which allows for good quality genotype calling that includes high-dose markers, and 2) sufficiently large sample size to allow the detection of recombination events in a hexaploid scenario. Additionally, we considered the uncertainty in the genotype calling in filtering problematic SNPs and also in modeling the error during the map construction using a hidden Markov model (HMM)^14^. Moreover, all methods can be readily used in tetraploid and octoploid full-sib populations.

## Results

### Genotype calling

Next-generation sequencing produced several millions of barcoded reads, resulting in approximately 41 million tags which were aligned against the genomes of two sweetpotato diploid relatives, *I. trifida* and *I. triloba*,^26^ resulting in 1,217,917 and 1,163,397 SNPs, respectively. We used the software SuperMASSA^27^ to call a total of 442,184 SNPs anchored to *I. trifída* genome and 438,808 anchored to *I. triloba* genome. After filtering out low-quality and redundant SNPs (Supplementary Fig. 1 A), we obtained 38,701 SNPs scored in 311 individuals. For all SNPs we obtained dosage-based calls and the associated probability distribution (exemplified in Fig. 1). From the total SNPs, 55.5% were classified as simplex (single-dose markers present in one parent) or double-simplex (single-dose markers present in both parents) and 44.5% were classified as multiplex (Supplementary Fig. 1 B).

**Fig. 1.**
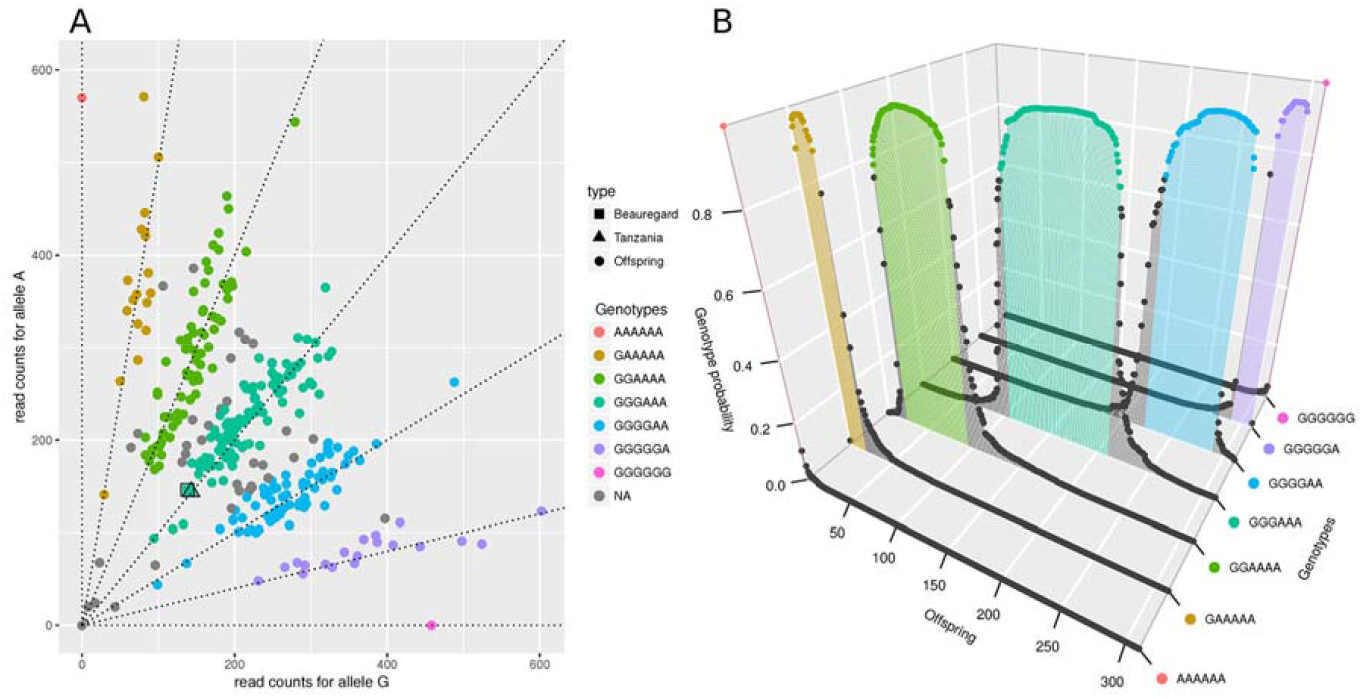
Example of genotype call of SNP *Tf_S1_30010438.* (A) Scatter plot of the read counts for the two allelic variants A and G. The axes represent the read counts of both allelic variants. Squared and triangle dots represent parents ‘Beauregard’ and ‘Tanzania’ respectively, and regular dots represent the offspring. Dashed lines indicate seven possible dosages in a hexaploid individual. The different colors indicate the dosages assigned to the individuals by SuperMASSA. The low number of individuals observed between genotypic classes (gray dots, with genotype probability smaller than 0.8), outlines a data set with low noise, producing a clear classification. The genotypes of both parents were estimated as three doses of the allelic variant A three doses of G. The genotype calling model also considered the expected Mendelian segregation ratio, which under random chromosome pairing is 1:18:99:164:99:18:1. (B) Inferred probability distribution of genotypes for each individual in the offspring. The colored dots correspond to individuals with the same genotypic classes in panel A. Loci where the highest posterior probability was smaller than 0.8 were assigned as missing data (gray dots).

### De novo map construction

To build the genetic map, we implemented the R package MAPpoly (https://github.com/mmollina/MAPpoly). The software comprises routines to perform all steps involved in the map construction of autopolyploid species using a combination of pairwise recombination fraction and HMM-based map estimation. Firstly, we obtained the recombination fractions and associated likelihoods for each possible linkage phase for all SNP pairs (~749 million pairs). Next, we selected the recombination fractions associated to the most likely linkage phase configuration for each SNP pair and applied the Unweighted Pair Group Method with Arithmetic Mean (UPGMA) hierarchical clustering. We formed 15 distinct clusters representing *I. batatas* homology groups (Supplementary Fig. 2). For the 15 groups, 93.4% of the SNPs were co-located on the same chromosomes in both references and LGs (Supplementary Table 1). These matched SNPs were selected to build the “de novo” map. Since each LG had the majority of their SNPs corresponding to a distinct chromosome in both references, LGs were numbered after the diploid references.

To order the SNPs in each LG, we used the Multidimensional Scaling (MDS) algorithm^28^. The reordered recombination fraction matrix is shown in Supplementary Fig. 3 A. With the proposed MDS order, the parental allelic variants were phased using the procedure presented by Mollinari and Garcia^14^. The algorithm is based on LOD scores of pair-wise markers as the first source of information to sequentially position the allelic variants in specific homologs. For situations where pairwise analysis had limited power, the algorithm used the likelihood of multiple markers in a Markov chain (see Material and Methods and Supplementary Note).

The “de novo” multilocus map is presented in Supplementary Fig. 3 B. The length of the LGs ranges from 723.7 centimorgans (cM) in LG 8 to 2,037.0 cM in LG 4, with a total map length of 20,201.8 cM and 32,200 SNPs (average inter-locus distance ~0.63 cM), with no considerable gaps between SNPs. Although MDS algorithm yielded adequate global marker orders for all LGs (Supplementary Fig. 3 C) the resulting map is considerably large. Two main reasons for this inflation are misplacement of closely linked SNPs and genotyping errors^14,29–31^, which will be systematically addressed in the next sections. The alignment of the “de novo” map against the reference genomes is shown in Supplementary Fig. 4. Despite several chromosomal rearrangements, we observed high levels of collinearity between both reference genomes and the estimated map. The collinearity extended in blocks with few megabase pairs (Mb), as in LGs 2 and 7, up to the whole chromosome in LGs 5, 9, 10, 11, 12, 14, and 15. In cases where the collinearity extended through the whole chromosome, we observed sites of suppressed recombination (plateaus in Supplementary Fig. 4), possibly indicating the location of centromeric regions.

### Genomic assisted map improvement and modeling of genotyping errors

To reduce the effects of the local marker misplacement in map inflation, we used *I. trifida* genome to propose alternative SNP orders within collinearity blocks and evaluated the likelihood of the resulting maps, keeping the one with the higher likelihood (see Material and Methods and Supplementary File 1). We used *I. trifida* as the primary reference genome because the quality of the assembly is superior and more closely related to *I. batatas* when compared to *I. triloba*^26^. After the order improvement, ~97.0% of the *I. trifida* SNPs present in the map were locally reordered (see Material and Methods and Supplementary File 1). From the remaining *I. trifida* SNPs, ~1.3% were kept in their original “de novo” order and ~1.7% were eliminated since their inclusion caused map inflation higher than 2.00 cM. We then positioned the SNPs private from *I. triloba* reference genome into the resulting map using the constraints imposed by both genomes (see Material and Methods and Supplementary Fig. 5, blue map). The genomic-assisted reordering resulted in a map with 30,723 SNPs spanning 12,937.3 cM with an average interlocus distance of ~0.42 cM, representing a reduction of 1.6-fold when compared to the de novo map. To address the effects of genotyping errors, we re-estimated the map using the probability distribution of the genotypes provided by SuperMASSA^27^ as prior information in the HMM emission function^14^, as implemented in MAPpoly (Supplementary Fig. 5, green map). In this case, the map length was 4,764.1 cM with an average inter-locus distance of ~0.16 cM, representing a map reduction of 2.7-fold when compared to the genomic-assisted map.

### Probability distribution of multiallelic genotypes and final map estimation

For all individuals in the BT offspring, we obtained the conditional probability distribution of the 400 possible hexaploid genotypes in the whole genome given the estimated genetic map. We used the Markovian process to propagate the information throughout each LG (see Material and Methods). The genotypic probability distribution at each genome position was assessed by using the information of all markers in the LG in all individuals of the full-sib population (Supplementary Table 2 and Supplementary Fig. 6). Next, we removed 13 individuals with inconsistent genotypic profiles (Supplementary Figs. 7 and 8) and, keeping the marker order, we re-estimated the final map considering 298 individuals. A comparison between the “de novo” and the final maps shows a length reduction of 7.5-fold due to the removal of spurious recombination events through the several steps of map improvement (Supplementary Fig. 5).

The final map contains 30,684 SNPs spanning 2,708.4 cM (average inter-locus distance of ~0.09 cM), with 60.7% simplex and double-simplex markers, and 39.3% multiplex (Table 1 and Fig. 2). All homologs showed allelic variations along the LGs indicating that their inheritance pattern can be assessed in the full-sib population. However, several LG segments showed identical composition for a subset of homologs, as shown by the *Genotypic Information Content* (GIC)^32^. In our results, 81.8% of all map positions in ‘Beauregard’ and 77.3 % in ‘Tanzania’ had a GIC > 80%, revealing that we can reliably trace back the inheritance of the most homologs from the offspring to the parents (Supplementary Fig. 9). A small number of homologs presented an identical allelic composition for certain segments, which is the case, for example, of homologs *i* and *j* for the most of LG 2 and *l* and *k* along the whole LG 11. The complete map can be interactively browsed at https://gt4sp-genetic-map.shinyapps.io/bt_map/. For a selected segment, the browser provides the name of markers, dosages in the parents and the linkage phase configuration of the allelic variants. Supplementary File 2 shows more map information, including the linkage phase configuration in both parents.

**Fig. 2.**
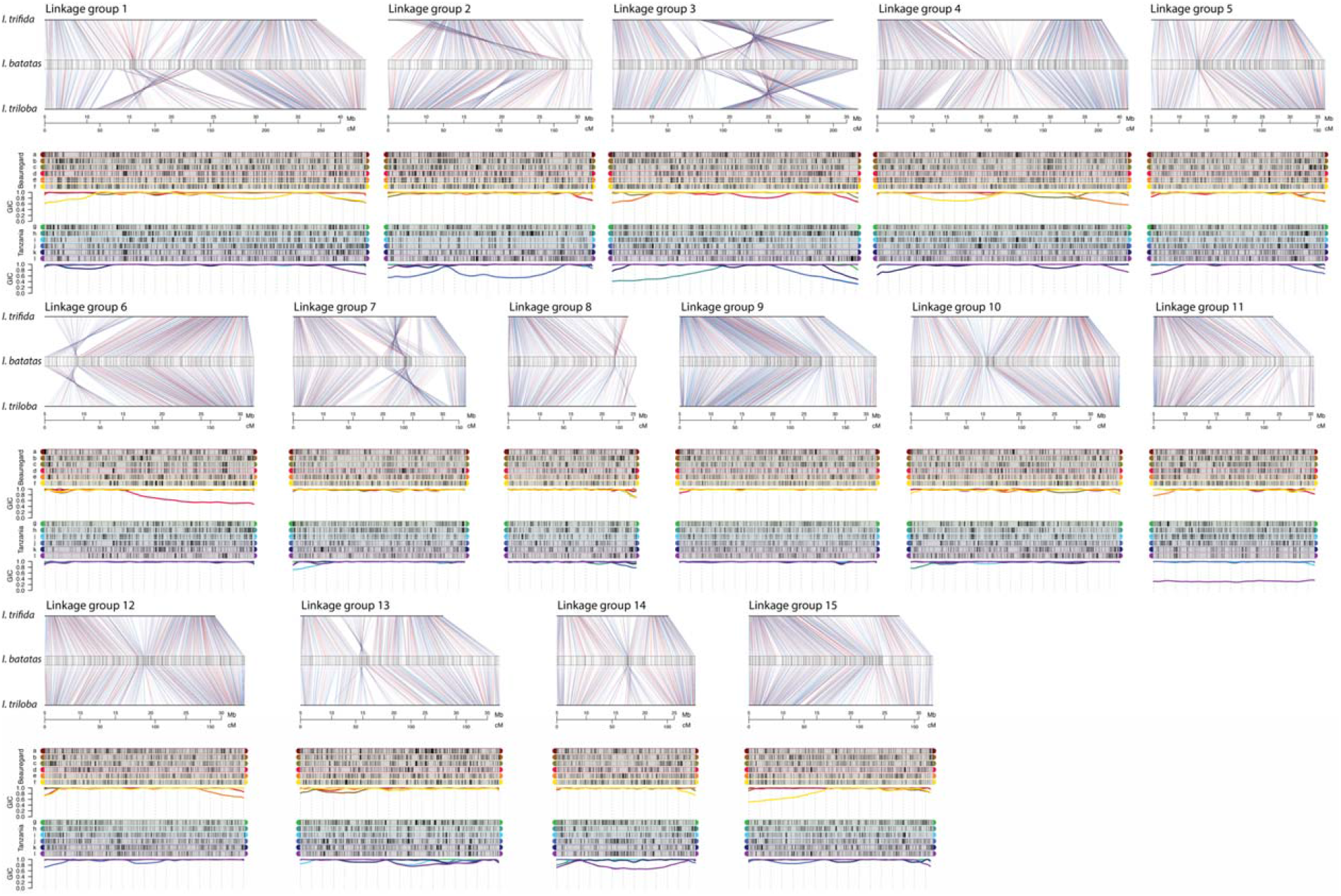
Sweetpotato genetic map. For each of the 15 LGs, we present the *I. batatas* genetic map with its SNPs anchored in both diploid reference genomes. Blue lines connecting the map and reference genomes indicate SNPs shared between *I. trifida* and *I. triloba* reference genomes and red lines indicate private SNPs. Above each map, we present a graphical representation of the parental linkage phase configuration of the homology groups for parents ‘Beauregard ‘and ‘Tanzania’. Black and gray rectangles indicate two allelic variants in each marker in all 12 parental homologs (6 × in ‘Beauregard’ and 6 × in ‘Tanzania’). The Genotypic Information Content (GIC), is presented below each homology group.

**Table 1.** Summary of sweetpotato genetic map

| LG | Length (cM) | Number of Markers |  |  | Total | SNPs/cM |
| --- | --- | --- | --- | --- | --- | --- |
|  |  | Simplex | Double-simplex | Multiple-dose |  |  |
| 1 | 290.9 | 1216 | 318 | 1211 | 2745 | 9.4 |
| 2 | 184.6 | 857 | 197 | 673 | 1727 | 9.4 |
| 3 | 222.1 | 1085 | 285 | 1052 | 2422 | 10.9 |
| 4 | 227.1 | 1372 | 377 | 1287 | 3036 | 13.4 |
| 5 | 157.1 | 892 | 193 | 816 | 1901 | 12.1 |
| 6 | 189.3 | 970 | 266 | 656 | 1892 | 10.0 |
| 7 | 156.3 | 1005 | 234 | 612 | 1851 | 11.8 |
| 8 | 115.5 | 712 | 140 | 312 | 1164 | 10.1 |
| 9 | 178.1 | 1403 | 261 | 715 | 2379 | 13.4 |
| 10 | 1188.7 | 1106 | 234 | 822 | 2162 | 11.5 |
| 11 | 145.6 | 724 | 177 | 729 | 1630 | 11.2 |
| 12 | 181.0 | 1367 | 246 | 1048 | 2661 | 14.7 |
| 13 | 180.1 | 761 | 174 | 743 | 1678 | 9.3 |
| 14 | 125.3 | 667 | 96 | 590 | 1353 | 10.8 |
| 15 | 166.6 | 1019 | 265 | 799 | 2083 | 12.5 |
| Total | 2708.3 | 15156 | 3463 | 12065 | 30684 | 11.3 |

Supplementary Tables 3 and 4 summarize the results of collinearity blocks containing the identical SNP sequences between *I. batatas* genetic map and *I. trifida* and *I. triloba* genomes, respectively. Thirty-nine blocks were aligned to 326.5 Mb of *I. trifida* genome, covering 96.5% of the *I. batatas* map (2,614.8 cM), with an average density of one SNP/14.2 kb; 107 blocks were aligned to 258.8 Mb of *I. triloba* genome, covering 83.1% of the map (2,251.8 cM), with an average density of one SNP/13.4 kb. The averaged genetic to physical map ratios for these regions were of ~124.8 kb per cM for *I. trifida* and ~114.9 kb per cM for *I. triloba.*

### Haplotype reconstruction and multivalent formation

To obtain the haplotype composition of all individuals in the full-sib population, we assessed the conditional probability distribution of the genotypes and appropriately combined them to build 12 profiles (one for each homolog) indicating the probability of inheritance of a particular homolog across the whole chromosomes for all individuals in the BT population (see Material and Methods). The results can be accessed at https://gt4sp-genetic-map.shinyapps.io/offspring_haplotype_BT_population/. By evaluating the recombination points and the homologs involved in the chromosomal exchange, we proposed a heuristic to obtain chains of homologs linked by recombination events (details in Material and Methods). These chains represent the inference of the meiotic process. Although they do not imply an exact pairing configuration, they can be classified according to the number of homologs involved in the chain. The number of parental homologs that are present in a single homolog of a particular offspring individual indicates the minimum valency of the meiotic configuration involved in its gamete formation (see example in Fig. 3). Thus, recombination chains with two homologs indicate the formation of at least a bivalent; three homologous, at least trivalent, and so on. For each LG, we calculated the percentage of the maximum number of homologs involved in the same recombination chain (Fig. 4). Most of the configurations involve recombination of two homologs (~73.8% in ‘Beauregard’ and 72.8 % in ‘Tanzania’) indicating that, homologs can undergo multivalent formation during the pachytene, though with no consequences of a multivalent formation in the majority of gametes formed. However, we also observed 12.8% of gametes in ‘Beauregard’ and 15.2% in ‘Tanzania’ with haplotype configurations involving three or four parental homologs in a single offspring homolog (indicating trivalent or tetravalent formation), and less than 2% of the meiotic configurations with five or six homologs (indicating pentavalent trivalent or hexavalent formation; details per LG in Supplementary Table 5). We also detected a significant positive linear correlation (*P* < 10^−3^) between the number of individuals with meiotic configurations originated from multivalent formations and the length of LGs (Supplementary Fig. 10).

**Fig. 3.**
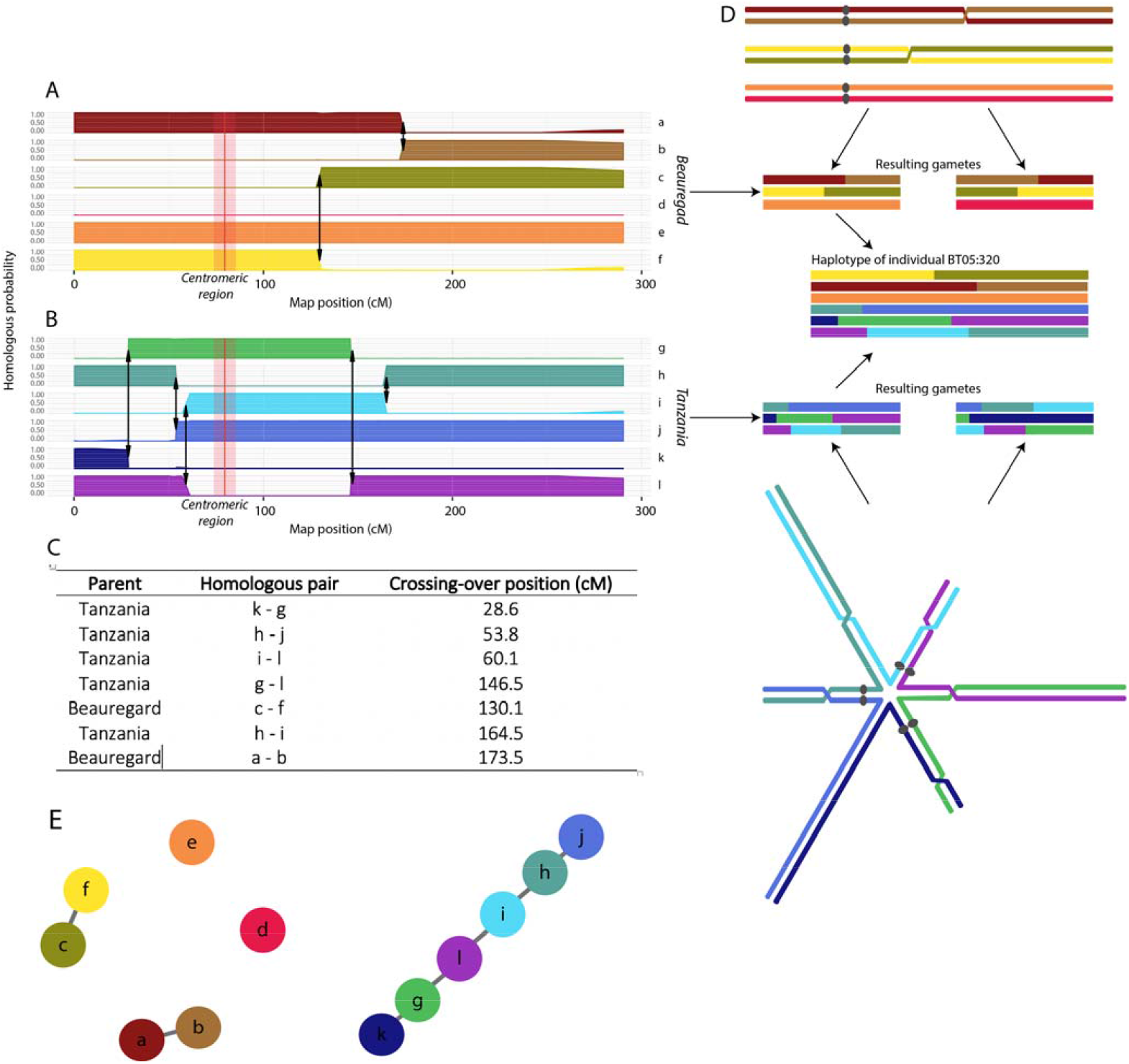
Example of haplotype reconstruction and distribution of meiotic configurations for individual BT05.320, linkage group 1. A) and B) Probability profiles for 12 homologs indicating the segments inherited from parents ‘Beauregard’ and ‘Tanzania’, respectively. The red line indicates the approximated centromeric region obtained using the *I. trifida* reference genome. The arrows indicate recombination points; C) Recombination signature table indicating the homolog pairs involved in each crossing-over and their position in the map; D) Possible meiotic configuration that originated gametes for individual BT05.320 in ‘Beauregard’ and ‘Tanzania’ and resulting gamete. Each chromosome is represented by one chromatid; E) Representation of the meiotic results as a graph: nodes represent the homologs and the edges represent recombination events between them.

**Fig. 4.**
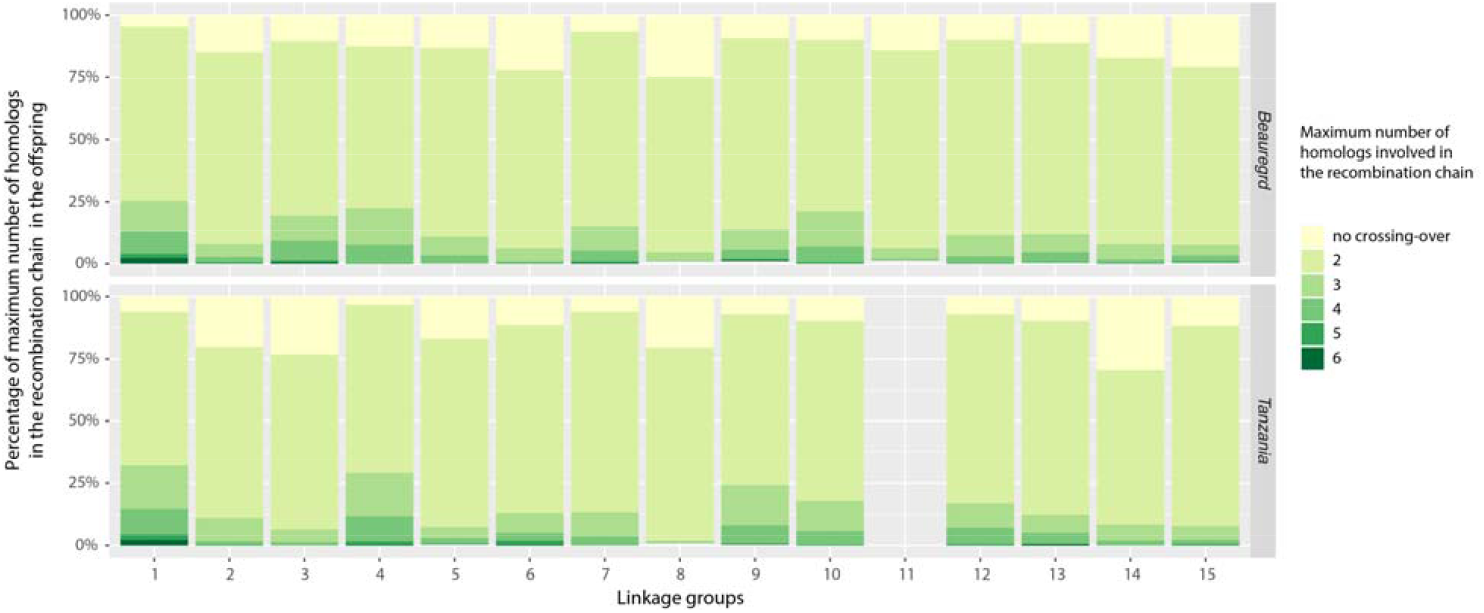
Percentage of maximum number of homologs connected in the same recombination chain during metaphase I in ‘Beauregard’ and ‘Tanzania’ for all 15 LGs. LG 11 for ‘Tanzania’ was mostly inconclusive and is not shown.

### Preferential pairing

In a hexaploid organism, there are 15 possible pairing configurations for a chromosome segment during the prophase I of meiosis. To assess the level of preferential pairing among homologs, we calculated the probability profile for each of the 15 possible pairing configurations across all LGs for parents (Fig. 5). We did not observe significant preferential pairing across the whole sweetpotato genome, except LG 2 (*p* < 10^−6^) which showed a low but significant preferential pairing. To further ascertain homolog preferential pairing, we evaluated the simplex marker information which confirmed our preferential pairing findings using the multilocus framework (Supplementary Fig. 11).

**Fig. 5.**
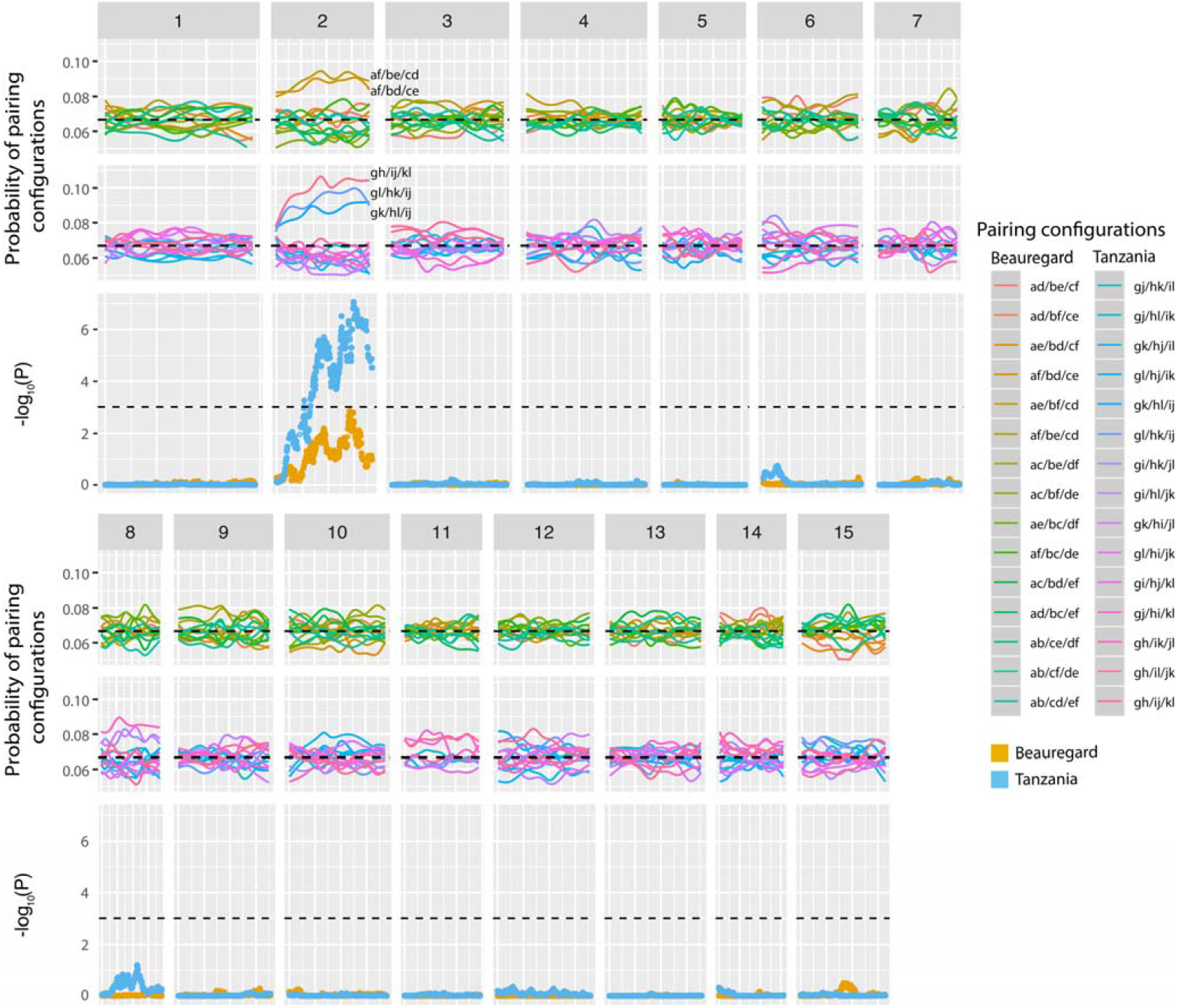
Preferential pairing profiles for 15 pairing configurations in parents ‘Beauregard’ and ‘Tanzania’ across 15 LGs. Notation *ab/cd/ef* indicates a configuration where homolog *a* paired with *b, c* with *d* and *e* with *f*. The dashed line in the probability profiles indicate the pairing probability expected under random pairing 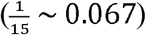. Orange and blue dots represent *log*_10_ *P* of a chi-square independence test for ‘Beauregard’ and ‘Tanzania’, respectively. Dashed line indicates 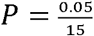. LG 2 presented a low, but significant preferential pairing involving three bivalent configurations (*gh/ij/kl, gl/hk/ij, and gk/hl/ij*). Homologs *i* and *j* appear in all three configurations, indicating a preferential association between these chromosomes.

## Discussion

We have built the first multilocus integrated genetic map of a hexaploid species, sweetpotato, using our newly implemented software MAPpoly. In the map, 90 homologs were densely represented in the 15 homology groups of cultivars ‘Beauregard and ‘Tanzania’ exhibiting high collinearity to two closely related diploid sweetpotato genomes, *I. trifida* and *I. triloba.* The high collinearity found by using our ultra-dense map corroborates with the levels of alignment (> 90%) between the diploid genomes and the parent ‘Tanzania’ reported by Wu et al.^26^, suggesting that the diploid genome assemblies could be used as robust references for the hexaploid sweetpotato. We also have constructed the hexaploid haplotypes of all individuals in the offspring, estimating the level of preferential pairing and multivalent formation during the meiotic process at a population level.

Haplotype inference is the ultimate attainment in linkage analysis since it contains the complete information about genome transmission across generations. The challenge of performing such inference, both in parents and offspring, would require new approaches to model the multiallelic transmission in a very complex meiotic scenario. Here we accomplished this by propagating the incomplete information of dosage-based SNPs throughout the LG using a Markov chain. As a result of the efficient combination of multiple SNPs, several LGs displayed fully informative parental haplotypes in most of their extension (Fig. 2 and Supplementary Fig. 9). Nevertheless, LG11 had two homologs (*k* and *l*) carrying the same allelic variations across its entire extension, which leads us to speculate that these two homologs were formed by nondisjunction of sister chromatids in meiosis II in one of Tanzania’s parent resulting in an unreduced gamete transmitted to the next generation^33^. Even though in some cases where not all homologs could be distinguished, we estimated their probability distribution, which can be readily used in further genetic studies, such as quantitative trait loci mapping performed for the BT population^34^. We also investigated how the pre-assembled parental homologs were transmitted to their offspring by assessing the probability distribution of the multiallelic genotypes across the whole genome for all individuals in the mapping population. Based on the inferred probability distributions, we presented a comprehensive probabilistic reconstruction of the haplotypes of all individuals in a full-sib hexaploid population. We found that ~15% of the offspring showed the evidence of multivalent formation, i.e., offspring homologs containing more than two parental homologs. This leads to intra-homolog variation, which could not be due to exclusive bivalent pairing.

Multivalent configurations often cause faulty chromosomal segregation leading to aneuploidy^35,36^. Such a phenomenon causes unbalanced gametes, and consequently the production of pollen and seeds with low viability, posing a significant hindrance to a stable genomic transmission throughout generations in polyploids^37^. Multivalents are usually observed in high numbers in recently formed polyploids, as in the case of the synthetic autopolyploid *Arabidopsis thaliana*^38^. Most of the established autopolyploids, however, show considerably fewer multivalents. In a survey involving 93 autopolyploid species, Ramsey and Schemske^39^ showed that the average frequency of bivalents was 63.7% whereas the average frequency of quadrivalents was 26.8%, which are significantly different from the theoretically expected (1 × two bivalents (II + II) to 2 × one quadrivalent VI)^4,40^. For hexaploids, the theoretical proportion of bivalent to multivalent configurations is 1 × three bivalents (II + II + II) to 6 × one tetravalent plus one bivalent (IV + II) to 8 × one hexavalent (VI)^40^. However, in our work, the number of multivalent signatures observed was notably low, whereas the number of bivalents was relatively high (Fig. 4). These results corroborate the previous cytological study by Magoon and coauthors^17^, who found similar levels of multivalent configurations in sweetpotato pachytene cells. Nevertheless, our results provide population-level evidence to the prevalence of bivalent configurations in sweetpotato meiosis.

In a scenario of scarce multivalent formation, the double reduction (DR) phenomenon becomes a somewhat rare event. The DR of a given locus is a consequence of a series of events: the occurrence of a crossing-over event between a locus and its centromere in a multivalent and subsequent migration of sister chromatids carrying a duplicated region to the same pole of the cell at anaphase I^41,42^. Such events could generate genotypes which are not observable under random chromatid segregation, potentially producing new genotypes. Multivalent formation is a necessary, but not sufficient condition for the occurrence of DR, which is expected to occur in low frequency^43^. Consequently, the observed low frequency of multivalent formation would indicate that the occurrence of DR events is much less likely. Although we did not take into account DR events during the construction of genetic map, it would have little impact to our map as the algorithm used here was found to be robust under low levels of multivalent formation^14^. Nevertheless, even a rare event, it could generate transgressive phenotypes that can be inherited through the next generations.

All sweetpotato genetic maps publish to date^9,18,19,44–47^ have acknowledged the hexasomic segregation in sweetpotato. However, none of them systematically characterized this phenomenon using the information of multiple markers assembled in complete hexaploid homology groups. Here we used the multilocus map to assess this information generating preferential pairing profiles (Fig. 5). We showed that even though sweetpotato origin could be traced to an interspecific hybridization as suggested by some studies^17,24,48,49^, its inheritance pattern is vastly autopolyploid-like and random chromosome pairing would enable recombination between sub-genomes across generations.

A variety of intrachromosomal rearrangements were observed between *I. batatas* map and *I. trifida* and *I. triloba* genomes. Rearrangements mapped to both diploid references, such as the chromosome inversion at the beginning of LG 6 (Fig. 2), represent structural changes exclusive to *I. batatas.* While the occurrence of such rearrangements could cause instability to meiotic process at some point of the evolutionary history of a polyploid species^50^, given the high level of bivalent signatures and the stable hexasomic segregation observed in our analysis, we concluded that these structural changes became fixed and do not cause major disturbances to the meiotic process in sweetpotato.

More than a linear order of genetic markers positioned in LGs, a genetic map is a statement about the inheritance pattern involved in the transmission of genome from parents and their offspring. A full characterization of this process can be achieved if the mapping method allows the estimation of haplotypes in both generations. In diploid organisms, a hidden Markov model was proposed by Lander and Green for linkage analysis of multiple markers^51^. Later on, several studies paved the way for a linkage map construction and haplotype inference in autotetraploid species^52–55^. However, for complex polyploids the map construction was restricted mostly to two-point marker analysis. We present the first integrated multilocus genetic map with fully phased haplotypes for both parents and offspring in a complex polyploid and, accompanied with it, the fully developed statistical methods and computational tool MAPpoly. This opens the door for detailed genetic analysis in complex polyploid species in general.

## Material and Methods

### Plant material

The mapping population consists of 315 full-sib individuals originated from a cross between the oranged-flesh cultivar ‘Beauregard’ (CIP440132 – male) and the African landrace ‘Tanzania’ (CIP440166 – female). These two cultivars were selected due to their agronomic importance and contrasting traits, such as, beta-carotene, dry matter, drought tolerance and resistance for viruses and nematodes^11,56^, for further QTL studies.

### Optimized genotyping-by-sequencing protocol – GBSpoly

Next-Generation Sequencing (NGS) library preparation protocol was optimized for polyploids and highly heterozygous genomes to produce uniform coverage across samples and loci, GBSpoly^57^ (details in Supplementary Note). The optimizations were based on re-engineered barcoded adapters that ensure accurate demultiplexing and base calls. The 6-9 bp variable length barcodes with designed to account for both substitution and indel errors (based on edit/levenshtein distance), minimizes phasing error and maintains nucleotide diversity at every position along the reads. A new feature, buffer sequences, upstream of the barcodes ensures that the barcodes lie in high-quality base regions by avoiding the elevated error rates at the ends of the reads. The adapters were ligated to fragments generated by double digests, *Tse*I and *Cvi*AII, and then size selected to minimize PCR bias. By designing barcodes that did not reconstitute the restriction sites, ligated fragments were subjected to a secondary digest to eliminate chimeric fragments. Sequencing was performed on the Illumina HiSeq 2500.

### Genotype calling

We used the software SuperMASSA^27^ to perform the genotype calling of parents and offspring of the full-sib population. For quality control purposes, we eliminated SNPs with read depth < 20 and estimated ploidy level different from six. We also filtered out SNPs with more than 25% of missing data and with segregation distortion (*P* < 5 × 10^−4^). Additionally, we removed four individuals with less than 100 reads on average for the selected SNPs (see Supplementary Note). We obtained the physical positions of the selected markers in two diploid reference genomes of *I. trifida* and *I. triloba*^26^ and classified them into shared between both genomes or private to a specific genome based on the full-sib population genotype calls.

### De novo map construction

#### Grouping and SNP ordering

We computed recombination fractions for all marker pairs considering all possible linkage phase configurations. For each marker pair, we selected the recombination fraction associated to the most likely linkage phase and assembled a recombination fraction matrix for all marker pairs. Using UPGMA hierarchical clustering we generated a dendrogram representing 15 LGs corresponding to the 15 sweetpotato homology groups. To order the SNPs in each LG, we converted the recombination fractions to genetic distances using Haldane’s map function and applied the unconstrained MDS algorithm with the squared linkage LOD Scores to construct the stress criterion ^28^.

#### Phasing and multilocus map estimation

The parental linkage phase configuration was obtained by serially adding markers to the map sequence and evaluating two-point likelihoods associated to possible configurations between the inserted markers and the ones already positioned. If the LOD Score between the two most likely configurations was less than ten for a subset of configurations, we compared the multipoint likelihoods to proceed to the next marker insertion. When the last marker was inserted, we re-estimated the multipoint recombination fractions between all adjacent markers^14^. For more details see Supplementary Note.

### Genome-assisted map improvement

Using the *I. trifida* reference, we detected collinearity blocks within each LG by visually inspecting abrupt breakages in the scatter plots continuity (Supplementary Fig. 4). For each collinearity block, we evaluated the multilocus likelihood associated with the “de novo” order and the order provided by *I. trifida* reference. We selected the maximum likelihood order for each block, tested several orientations among them (Supplementary File 1) and chose the configuration that yielded the highest multilocus likelihood for the complete map. Next, we inserted the remaining private SNPs from *I. triloba* using the genomic position constraints imposed by SNPs shared by both genomes. We also eliminated SNPs that caused substantial map expansions (see Supplementary Note). Finally, we re-estimated the map by considering the probability distribution of the genotypes provided by SuperMASSA^27^. We also computed the GIC^32^ for each homolog across the entire genome.

### Probability distribution of the offspring genotypes

The probability distribution for all possible 400 hexaploid genotypes was calculated using the HMM framework detailed in Supplementary Note. Briefly, if 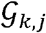 denote the *j*^th^ genotype, *j* ∈ {1,…,400} of an individual in a hexaploid full-sib population at locus *k*, the conditional probability distribution of 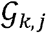 is defined as

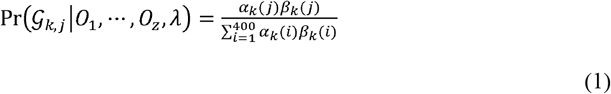

where *O*_1_,…,*O_Z_* is a sequence of observations of *z* markers, *λ* denotes the map parameters, *α_k_*(*j*) denotes the joint probability of the partial observation sequence to the left of marker *k* (including *k*) and genotype 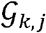, given the map parameters *λ*; similarly, *β_k_*(*j*) denotes the probability of the partial observation sequence to the right of the position *k* given the genotype 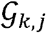 and the map parameters *λ*. The quantities *α_k_*(*j*) and *β_k_*(*j*) can be obtained using the classical *forward-backward* algorithm^58,59^ and their derivation is presented in Supplementary Note.

### Offspring haplotype reconstruction

The probability that an offspring individual carries a specific parental homolog 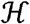 at position *k* can be obtained using

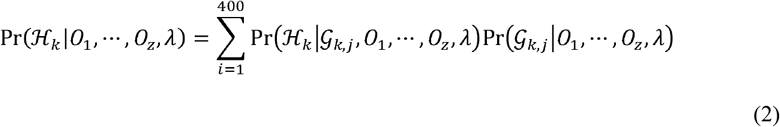

where, 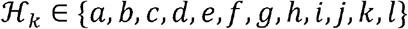 is the inherited homolog at locus *k*, 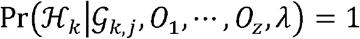 if 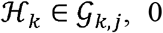 otherwise (see Supplementary Table 2). We obtained the haplotype probability profile for all 15 homology groups (one curve for each homolog, from *a* through *1*) for all individual in the bi-parental cross population by computing 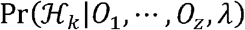 at every marker across the genome.

### Heuristic algorithm to detect crossing-over events

Given the probabilistic nature of the haplotype profiles, we proposed the following heuristic algorithm to detect crossing-over events:

1. Regions with haplotype probabilities greater than 0.8 are assumed to be 1.0, otherwise 0.0, forming a binary profile;
2. SNPs within a continuous segment of homolog or gaps flanked by crossing-overs smaller than 10 cM are removed.
3. If the remaining SNPs represent 20% or more of all SNPs in the analyzed LG, use Eq. (1) to re-estimate the 400 genotypes across the whole LG and compute a new homolog probability profile using Eq. (2). Otherwise, consider the probability profile inconclusive.
4. The crossing-over points are assessed by checking the points of probability transition across the LG. Homologs involved in the chromosomal exchange can be trivially assessed.
5. Exchange points closer than 0.5 cM are considered inconclusive since the haplotypes involved in the exchange could be erroneously assigned due to the lack of resolution in the mapping population.

We applied this procedure to the 15 LGs of all individuals in the population. We also present an interactive version of the heuristic algorithm at https://gt4sp-genetic-map.shinyapps.io/offspring_haplotype_BT_population/

### Preferential pairing profiles

Considering that all homologs pair during a hexaploid meiosis, there are 15 possible pairing configurations for a chromosomal segment. Let Ψ = {*ψ_i_*}, *i* = 1, …, 15 denote a set containing all 15 possible configurations^14^ (see Supplementary Note). The posterior probability distribution of the pairing configurations at any position in the genome can be computed using

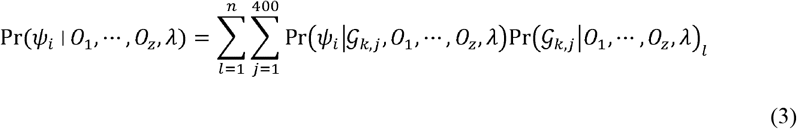

where *n* is the number of individuals in the population, 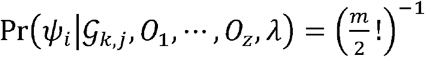 (*m* = 6 for hexaploids) if 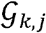 is consistent with *ψ_i_* i.e., if genotype 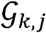 can be originated from the pairing configuration *ψ_i_*, 0 otherwise^14^. To test whether the observed frequencies of the 15 bivalent configurations differ from the expected under random pairing 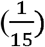, we used the *χ*^2^ test with *P* < 0.001 to declare significance. We also used the likelihood associated to recombination fractions of single-dose markers to assess preferential pairing, as suggested by Wu and co-authors^60^.

Further details of methods are given in the Supplementary Note.

