## Supplementary Note for "Unraveling the hexaploid sweetpotato inheritance using ultra-dense multilocus mapping"

**2 Brief description of the hidden Markov model proposed by Mollinari and Garcia (2019)** **12**

**3 Supplementary Figures** **16**

### 1 Extended Material and Methods

#### 1.1 GBSpoly analysis and SNP mining

##### 1.1.1 Selection of restriction enzyme combination

Using the SimRAD package in the R software[1], *in silico* double digests were performed on assembled chloroplast[2] and nuclear genomes of the putative diploid ancestral[3] progenitors of hexaploid sweetpotato (i.e. *I. trifida* and *I. triloba*). Various restriction enzyme combinations were evaluated to determine a combination that produces the most abundant nuclear DNA fragments between 100 and 500 bp long, but with limited chloroplast (several thousand copies per cell) DNA fragment contamination (within the same size range). Similar to typical double digest GBS protocols, one of the enzymes is methylation sensitive in order to avoid producing fragments derived from repetitive sequences such as transposons, which are difficult to use for SNP-calling and genetic analyses. The *Tse*I/*Cvi*AII restriction enzyme combination performed the best for various parameters evaluated (Fig. 1). The number of fragments generated is only a relative estimate between enzyme combinations, and is not an effort to replicate *in vitro* conditions, especially since methylated DNA will not be digested and represented in the library. The *Tse*I/*Cvi*AII combination provides the flexibility of generating low, medium or high-density markers by varying the window size selection. This is useful for applications like genetic linkage mapping that only require a few thousand markers, or genome-wide association analysis that require tens/hundreds of thousand of markers.

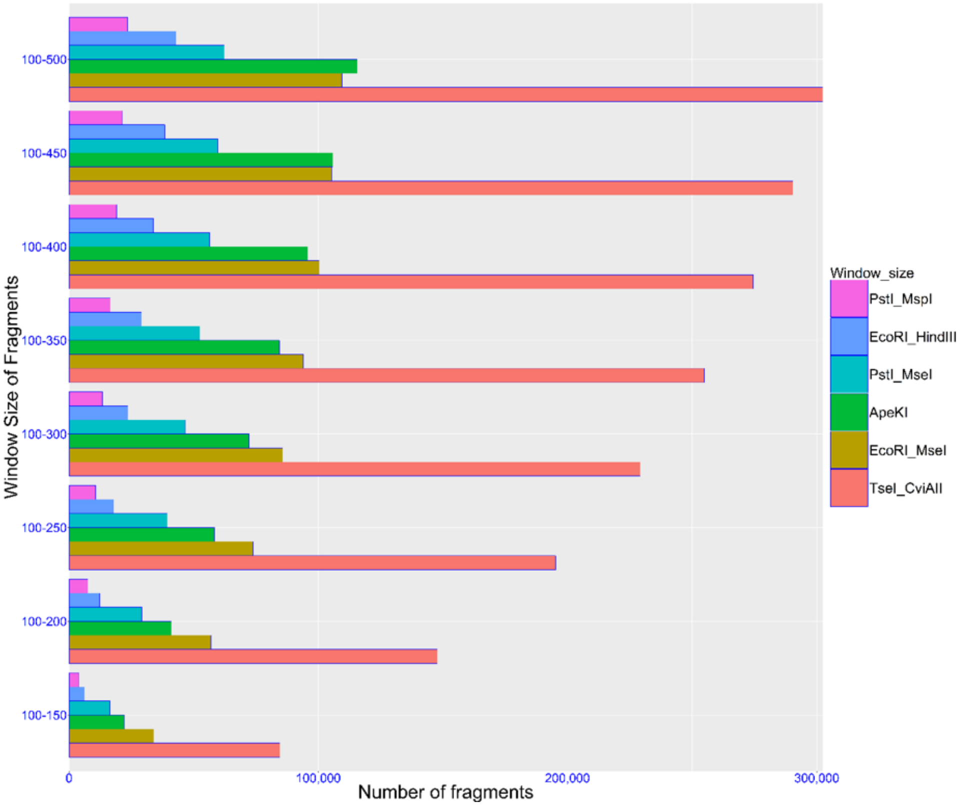

**Figure 1.** in silico double digest of *I. trifida* and *I. triloba* draft nuclear and complete chloroplast genome assemblies.

##### 1.1.2 Buffer and barcode sequence design

Even though the Illumina sequencing platform has a high overall base calling accuracy rate, the ends of the reads tend to have higher error rates. This is particularly important for de-multiplexing barcoded pooled samples since the barcode sequences themselves lie within this high error rate region. To resolve this problem, we have introduced a set of ninety-six 8 bp buffer sequences upstream of the barcodes (Fig. 2). This shifts the barcode sequence into a region that has high base calling accuracy rates. The high nucleotide diversity (having equal proportions of A, C, G and T nucleotides at each base position), which is required for optimal performance on the Illumina platform, was maintained among the buffer sequences. Due to the introduction of the buffer sequence, we were able to assign near 100% of all reads to individual samples after de-multiplexing. Using the buffer sequence (elevated error rate in first 6 bases) for de-multiplexing, we empirically determined that less than 50% of the reads could be recovered accurately. Recovery of more reads can be achieved by allowing for one or more base mismatches in these barcode sequences, but this can lead to high error rates. The barcode sequences are central to obtaining high quality SNP/variants during GBS. The two most important elements of the barcode design are their variable lengths (6-9 bp in this protocol) and the edit/Levenshtein distance, which accounts for both nucleotide substitution and indel errors[4]. Substitution or indel errors can cause a barcode to take on the sequence of another barcode, hence, assigning the reads to the wrong sample within the pool. Also, such errors can result in the inability to assign reads to any barcode altogether leading to lower recovery of reads. High edit distances between barcode sequences are ensured and this provides accuracy for de-multiplexing, compared to the hamming distance that only accounts for substitution errors. The variable length nature of the barcodes is important for phasing errors associated with the non-variable sequence restriction site (*Tse*I) common across all pooled samples. The diversity in the buffer sequence and barcodes also ensures reduced phasing errors. The variable length leads to the restriction site sequence to be staggered among samples, hence introducing variability at each base position in the library. This is important for nucleotide diversity at each base position. Additional modifications to the protocol include design of adapters to eliminate restriction enzyme recognition site upon ligation. This is important because our modified protocol introduces a step that eliminates chimeric sequences by performing a post-ligation digest with the same *Tse*I/*Cvi*AII enzyme pair. Typically, these chimeric sequences are useless, as they do not align to reference genome, or at best they could be erroneously mapped. Single target DNA fragments incorporated into adapters will remain intact during the secondary digest since the restriction site is destroyed upon ligation with adapters.

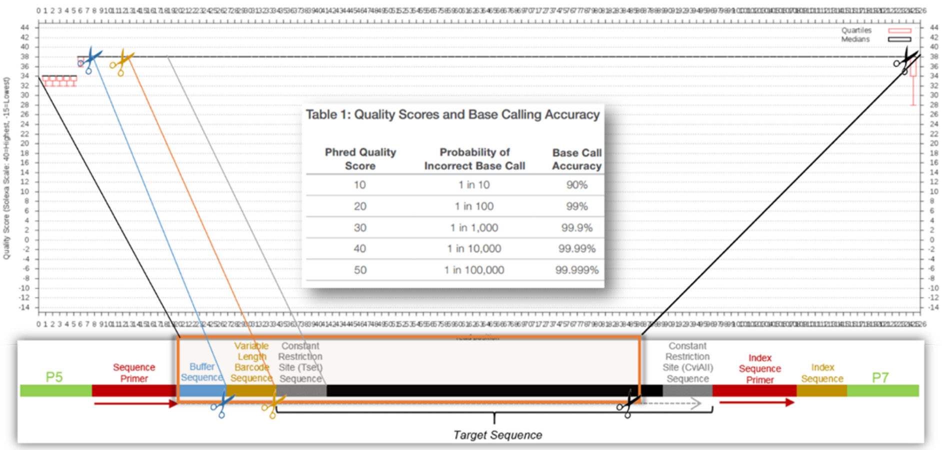

**Figure 2.** Structure of adapters (including the buffer and barcode sequences) and incorporated target DNA fragments used for the GBSpoly protocol. Plot of quality scores at each base calling position are shown before trimming.

##### 1.1.3 Modifications to GBS protocol during library construction

The main modification in our current protocol[5] ensures that all loci and pooled samples are sequenced with uniform read depth. An imbalance in read depth translates to a high proportion of missing data, as well as the inability to resolve heterozygous genotypes and dosage in loci lacking sufficient read depth. This is predominantly due to a PCR bias, where smaller fragments are amplified more efficiently than larger fragments. To reduce this bias, we adopted a size selection step, so that variation in the length of fragments is significantly reduced. Additionally, to eliminate chimeric sequences that can lead to unusable reads, or erroneous mapping them, we have introduced a second round of double digests of the pooled samples. The steps in the protocol include:

**Step 1**: Extract DNA using the CTAB method[6] in a 96 well plate format, quantify samples using an inexpensive absorbance-based method, normalize concentration to 50-200 ng/ul and run DNA samples on a 1% agarose gel to ensure there is no smearing and no significant RNA contamination. An estimate of DNA quantity is sufficient at this step, but a more sensitive florescent-based quantification method (optional in updated protocol) will be performed downstream, as concentrations will change during post-digest cleanup. Digestion is performed as shown in Table 1:

**Table 1.** Sequential double digest of DNA samples.

| Two-step double digest | Vol. (ul) |
|---|---|
| Step I: | |
| 50-200 ng/ul DNA | 8 |
| Molecular grade H2O | 18.5 |
| 10x CutSmart Buffer | 3 |
| *CviA*II (10 units/ul) | 0.5 |
| Total Volume | 30 |
| Thermocycler: 25°C for 3 hrs, 65°C for 20 mins, Hold 4°C ∞ | |
| Step II: | |
| Step 1 reaction | 30 |
| Molecular grade H2O | 4 |
| 10x CutSmart Buffer | 0.5 |
| *Tse*I (10 units/ul) | 0.5 |
| Total Volume | 35 |
| Thermocycler: 65°C for 3 hrs (or overnight), then hold 4°C ∞ | |

**Step 2**: MagBead Purification step using AMPure XP beads (not required in updated protocol):

- Add 52.5 ul (1.5x volume) of AMPure XP beads to each digested sample (from step 1).
- Pipette up and down to mix properly and incubate for 5 mins at room temperature.
- Place on a magnetic stand to collect beads, wait until the solution is clear.
- Aspirate the solution with a pipette set to 75 ul and discard.
- Wash once with 200 ul of freshly made 70% ethanol. To avoid beads floating too high in well, leave plate on the magnetic stand.
- Wash once with 200 ul of freshly made 70% ethanol. To avoid beads floating too high in well, leave plate on the magnetic stand.
- Aspirate the ethanol completely and leave to air dry for 5-7 mins at room temperature.
- Add 35 ul of water, remove plate from magnetic stand and mix by pipetting up and down.
- Return plate to magnetic stand and recover only 25 ul of solution containing DNA.
- Quantify cleaned-up post-digested DNA samples on a plate reader using the picogreen assay and normalize DNA concentration to a concentration between 5 to 10 ng/ul (or a lesser concentration within limits of detection).

**Step 3:** Pool product of ligation reaction (Table 2) between barcoded adapters and genomic fragments (normalized concentrations of digested genomic DNA fragments were used as limiting reactant in reaction. In updated protocol, the adapters are the limiting reactant. This ensures limited labor and lower cost of library prep):

- Make a master mix of buffer, ATP, water and add ligase last.
- To 25 ul of digested DNA, add 5 uL of adapter mix to each well
- Then add 15 ul of master mix made in step “a” to each well (Thermocycler: 22° C for 2 hrs, 65°C for 20 mins, Hold 12°C inf. Ligation can be stored at -20°C)
- Pool samples by pipetting 5 ul of each ligated DNA sample (96 samples per pool for diploid genomes and 64 samples per pool for hexaploid genomes).

**Table 2.** Ligation reaction of digested DNA samples.

| Ligation reaction | Volume (ul) |
|---|---|
| 125-250 ng of digested DNA (from step 2) | 25 |
| Adapter mix (combined P2 [0.8 uM] & P1 [3.4 uM]) | 5 |
| 10x CutSmart Buffer | 4.5 |
| ATP (final concentration of 1 mM) | 0.5 |
| Molecular grade H2O | 9.5 |
| T4 DNA ligase | 0.5 |
| Total volume | 45 |

**Table 3.** Secondary double digest of ligated and 64 pooled samples.

| Remove chimeric sequences | Vol (ul) | Remove chimeric sequences | Vol (ul) |
|---|---|---|---|
| Step 1: | | Step 2: | |
| Pooled DNA | 320 | Step 1 reaction | 321 |
| *CviA*II(or NlaIII) (10 units/ul) | 1 | *Tse*I (10 units/ul) | 1 |
| Thermocycler: 25°C (37°C for NlaIII) for 3 hrs, 65°C for 20 mins, Hold 4°C $infty$; | | Thermocycler: 65°C for 3 hrs, 65°C for 20 mins, Hold 4°C $infty$; | |

**Step 4:** Secondary digest of ligated and pooled samples to eliminate chimeric sequences (Table 3):
**Step 5:** MagBead Purification of Pooled Libraries:

- Add 483 ul (1.5x volume) AMPure beads to each pool of samples from step 4, mix by pipetting up and down, then incubate for 5 mins at room temperature.
- Place tube on a magnetic stand to collect beads. Allow solution to clear, then aspirate solution with a pipette set to 700 ul and discard.
- Wash once with 200 ul of freshly made 70% ethanol. To avoid beads floating too high in well, leave plate on the magnetic stand.
- Aspirate the ethanol completely and leave to air dry for 5-7 mins at room temperature.
- Add 40 ul of water, remove plate from magnetic stand and mix by pipetting up and down.
- Return plate to magnetic stand and recover only 30 ul of solution containing purified DNA.

**Step 6:** Size selection with PippinPrep and Magbeads:

- Quantify pooled and purified library with Picogreen assay. Note a maximum quantity 10 ug of DNA per lane can run on PippinPrep.
- Run on the PippinPrep to the specific fragment size you want to select. The libraries for the diploid and hexaploid mapping populations were created by size selection within a range of 250-450 bp and 300-400 bp, respectively.
- Elute each aliquot to 40 ul two times (1st and 2nd elution for each pool) and run the eluted samples on a Tapestation or Bioanalyzer instrument (Figure 3) to quantify and check size selection.
- Pool the two 40 ul elutions (80 uL per pool) and calculate the average concentration per pool (Note: Do not pool the samples if the concentration of the second elution is too low).

**Step 7:** Amplification of the size selected library (Table 4):

- Clean-up PCR product using PippinPrep (remove fragments outside size selection window that might have been carried over and enriched by PCR amplification).
- Perform MagBead purification (like in step 5). This ensures exclusion of small fragments outside size selection window, which is not completely eliminated with PippinPrep.

**Table 4.** PCR reaction setup for library of pooled samples.

| PCR reaction mix | |
|---|---|
| Library (@ approximately 20 pg/ul) | 10 ul |
| Phusion HF PCR masterMix (2x) | 12.5 ul |
| 10uM Reverse Primer (P1) | 1.25 ul |
| 10uM Forward Primer (P2) | 1.25 ul |
| Total reaction volume | 25 ul |
| PCR: 94°C (5 mins); 94°C (15 sec), 65°C (30 sec), 72°C (60 sec) for 18 cycles; 72°C (5min); 4°C (forever) | |

- Run samples on a Tapestation or Bioanalyzer instrument to ensure proper size selection (Fig. 3).

**Step 9:** Sequencing set-up for the HiSeq 2500 Illumina platform at UNC or NCSU:

- Submit a 1: 25 dilution of the 60 uL amplified library and quantify on Tapestation.
- From this use the pmol/l (molarity) to calculate the nmol/l of the amplified library
- Dilute the stock amplified library to 10 nmol/l into 20 ul for sequencing on Illumina HiSeq 2500.

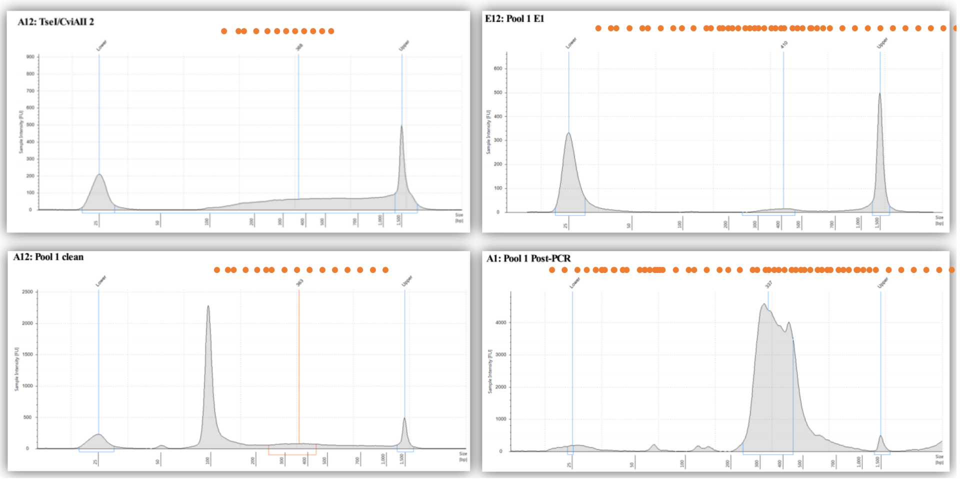

**Figure 3.** Electropherograms showing concentrations and profile of fragment sizes from digested DNA to PCR products of size selected libraries. Small peaks outside size selection range (bottom right) reduce sequence yield and is eliminated by additional Magbead purification.

#### 1.2 Genotype calling and Quality control

We used the software SuperMASSA[7] to perform the genotype calling of all individuals in the BT population. For all loci, parents and offspring were assigned to genotypic classes based on the maximum *a posteriori* probability (MAP) of a Bayesian network which took into account the ratio of read-counts for both allelic variants as well as the expected Mendelian segregation under random chromosome pairing in the full-sib population. For quality control purposes, we eliminated SNPs with less than 20 reads on average and also removed data points with read-counts smaller than 10% of the maximum read-count in the offspring[8]. Also as part of quality control, we tested ploidy levels 2, 4 and 6 to find the MAP of the model and only considered solutions with estimated ploidy level 6[9]. With a modification in the SuperMASSA software, we obtained the posterior probability distribution of the genotypic classes for each locus for all individuals in the offspring. We considered as missing data those loci where the highest posterior probability was smaller than 0.8. Loci with more than 25% missing data were not used in the analysis. We also tested the segregation of the SNPs using a classic chi-square test. We reject $H_0$ (Mendelian polysomic segregation under random chromosome pairing) if $P < 5 \times 10^{-4}$. Additionally, we removed four individuals (*BT13.315*, *BT05.342*, *BT13.286*, and *BT13.342*) with less than 100 reads on average for the selected SNPs. To avoid sampling of SNPs from duplicated sites across the genome, we also removed 101 SNPs with average read counts higher than 1.5× the interquartile range of the read count distribution of all SNPs. Using the tags aligned against the genomes of *I. trifida* and *I. triloba*[3], we obtained the physical position of the SNPs in those genomes. The resulting datasets were merged, and SNPs presenting at least one different genotype in the offspring were considered private either to *I. trifida* or *I.triloba* genomes and were named $Tf\_Sx\_y$ and $Tl\_Sz\_w$, respectively, where $x$ and $z$ are the relative chromosome in the reference genomes, and $y$ is the position (in base-pairs) in *I. trifida* genome, and $w$ is the position in *I. triloba* genome. We used a similar nomenclature ($Tf\_Tl\_Sx\_ySz\_w$) for SNPs were all individuals in the offspring presented the exact same genotypes.

#### 1.3 *De novo* map construction

##### 1.3.1 Grouping and SNP ordering

For each SNP pair, we obtained the recombination fractions and the associated likelihood for each possible linkage phase[10]. We used the most likely linkage phase as the baseline to compute LOD Scores associated with all linkage phases, or $LOD_{ph}$, (i.e., the most likely configuration has a $LOD$ equal to 0.00). We also obtained the classical LOD Score of linkage analysis ($LOD_{rf}$) using the estimated recombination fraction and the likelihood for unlinked SNPs ($r = 0.5$). In this case, high values of LOD Score indicate evidence of linkage. We applied the UPGMA hierarchical clustering to all pairwise recombination fractions associated with the most likely linkage phase configuration and generated a dendrogram representing 15 LGs corresponding to the 15 sweetpotato homology groups. To build a framework map, we considered only SNPs assigned to the same LG using both physical and linkage information. For each LG, we selected recombination fractions such as $\hat{r} < 0.15$, $LOD_{rf} > 5.0$ and the second most likely linkage phase configuration with $LOD_{ph} > 5.0$. SNPs that do not meet these criteria for at least 1% of the pairwise combinations with other SNPs on the same LG were not used in subsequent analysis. We applied the MDS algorithm[11] to perform a *de novo* ordering of SNPs within each LG. We obtained the recombination fraction matrices using the most likely linkage phase configuration ($LOD_{ph} = 0.00$). We used the unconstrained MDS and the squared linkage LOD Scores ($LOD^2_{rf}$) to construct the stress criterion[11]. To convert the pairwise recombination fractions in genetic distances we used the Haldane map function, which was also used throughout this work. Since we had a filtering procedure after the recombination fraction computation and the subsequent phasing procedure removes markers that may cause disturbances in the map, we opted to run the MDS algorithm just once. We also inspect the monotonicity of the resulting re-ordered recombination fraction matrices to find possible ordering problems, with no SNPs removed at this point.

##### 1.3.2 Phasing and multilocus map estimation

Parental allelic variants were phased using the procedure presented in Mollinari and Garcia (2019)[10]. In brief, assume a sequence of $n$ ordered markers $M_1, M_2, \cdots, M_n$. For the first pair of markers $\{M_1, M_2\}$ we enumerate all possible linkage phase configurations and eliminated those with $LOD_{ph} > 10$ ($\eta$[10]). We then evaluate $LOD_{ph}$ between the next marker ($M_3$) and all previously positioned ones ($\{M_1, M_2\}$) eliminating configurations with $LOD_{ph} > 10$. We then computed the multilocus sub-map including markers $\{M_1, M_2, M_3\}$ considering the selected configuration phases. Hidden Markov model based (HMM) likelihoods of the resulting maps were assessed in terms of LOD Scores ($LOD_{ph_m}$) and, similarly to $LOD_{ph}$, the most likely configuration was used as the baseline[10]. Configurations with $LOD_{ph_m} > 10$ were eliminated for the next round of marker insertion. This procedure was repeated until all $n$ markers were positioned. If a complete map presents more than one linkage phase configuration, we choose the one with the highest multilocus likelihood. For saturated maps, as the ones presented in this study, we used three approaches to speed up the phasing process: First, instead of using the two-point information of all positioned markers, we limited our search to a sub-map of the last $k$ inserted markers where $k$ is sufficiently large to differentiate all 12 homologs, but never smaller than 50. Second, if the number of linkage phase configurations yielded by the two-point analysis was bigger than 20, we did not position the marker. Finally, we did not insert markers that caused a sub-map expansion higher than 2 centimorgans (cM), since such expansion is not expected in highly saturated maps. The final map was reconstructed using the HMM-based multipoint algorithm proposed in Mollinari and Garcia (2019) [10]. All analysis described in this section were performed using the software MAPpoly, available at https://github.com/mmollina/MAPpoly.

#### 1.4 Genome-assisted map improvement

SNPs positioned in the *de novo* map were aligned against the diploid reference genomes (*I. trifida* and *I. triloba*) [3] generating 15 scatter plots (Supplementary Figure 4). In this section, we refer to LG 7 (Figure 4) to describe the improvement procedure. The remaining LGs are presented in Supplementary File 1. Using the *I. trifida* reference, we detected collinearity blocks by visually inspecting abrupt breakages in the scatter plots continuity (Figure 4 A). For each collinearity block, we reordered the SNPs according to the *I. trifida* reference genome and computed the HMM-based log-likelihood of the *de novo* and the genome-based orders (Figure 4 B). We selected the orders with the highest log-likelihood; if the *de novo* order yielded the highest log-likelihood, we used both orders of that block in the next step. In a second visual inspection, we proposed a set of plausible orders and orientations of the collinearity blocks. Using the linkage phase configuration from the *de novo* map, we computed the log-likelihood of the resulting map obtained by concatenating the blocks (Figure 4 C). The result with the highest log-likelihood was selected as the genome-assisted map for that LG. For all LGs, we compared the log-likelihoods of the original *de novo* map and the genome-assisted map, to confirm the superiority of the genome-assisted map. In all cases the log-likelihood of the genome-assisted maps was substantially higher when compared to the *de novo* map (Figure 4 D). During the segment inspection, 411 SNPs that caused map inconsistencies such as map expansion and gaps between adjacent markers (1.7% of the *I. trifida* SNPs) were removed.

We then inserted the remaining private SNPs from *I. triloba* using the genomic position constrains imposed by SNPs shared by both genomes (Figure 5). If there were private *I. trifida* SNPs between the shared SNPs, the private *I. triloba* SNP was inserted in all possible inter-marker intervals and was positioned in the interval that yielded the highest multilocus likelihood for a sub-map containing 15 SNPs to both sides of the shared SNPs. SNPs that needed to be tested in more than 100 intervals in the sub-map were eliminated. We also eliminated SNPs that caused an expansion in the sub-map greater than the maximum inter-marker distance presented in the original sub-map. Using the same strategy, we included SNPs that were eliminated during the visual inspection. Finally, using the resulting map containing SNPs anchored in both genomes, we re-estimated the final multilocus map considering the probability distribution of the marker dosage genotypes provided by SuperMASSA [7]. As a means to detect the information content along the genome given the multi-point map, we also computed the *Genotypic Information Content* (GIC) using the method proposed by Bourke and co-authors [12].

#### 1.5 Genotype probabilities and offspring haplotype reconstruction

A cross between two hexaploid individuals potentially yields 400 distinct genotypes for a specific locus in the genome. The probability distribution underling these genotypes was calculated using the HMM framework briefly explained in Section 2 and presented in details in Mollinari and Garcia [10]. Let $\mathcal{G}_{k,j}$ denote the $j$-th genotype, $j \in \{1, \cdots, 400\}$, of an individual in a hexaploid full-sib population at locus $k$. Supplementary Table 2 shows all 400 possible genotypes in an full-sib hexaploid population. The

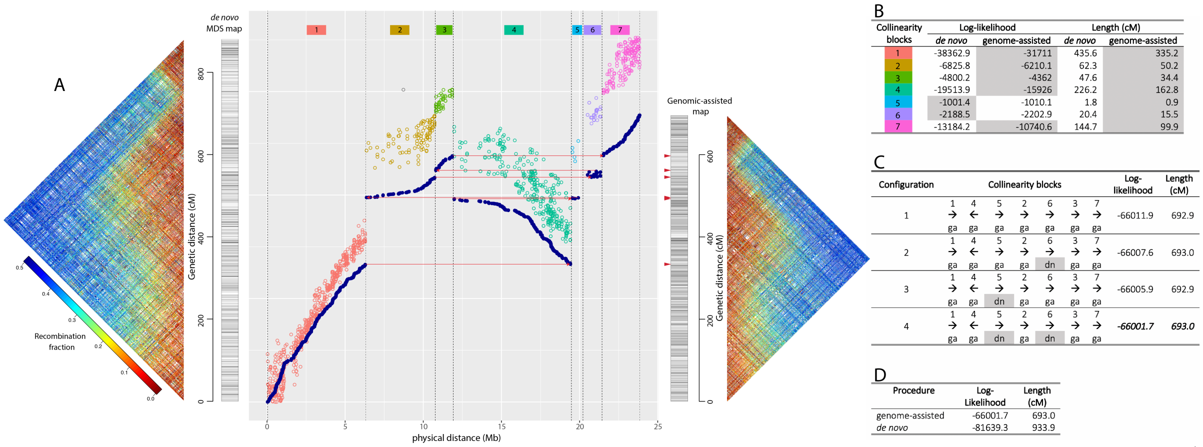

B

| Collinearity blocks | Log-likelihood | | Length (cM) | |
|---|---|---|---|---|
| | *de novo* | genome-assisted | *de novo* | genome-assisted |
| 1 | -38362.9 | -31711 | 435.6 | 335.2 |
| 2 | -6825.8 | -6210.1 | 62.3 | 50.2 |
| 3 | -4800.2 | -4362 | 47.6 | 34.4 |
| 4 | -19513.9 | -15926 | 226.2 | 162.8 |
| 5 | -1001.4 | -1010.1 | 1.8 | 0.9 |
| 6 | -2188.5 | -2202.9 | 20.4 | 15.5 |
| 7 | -13184.2 | -10740.6 | 144.7 | 99.9 |

C

| Configuration | Collinearity blocks | | | | | | | Log-likelihood | Length (cM) |
|---|---|---|---|---|---|---|---|---|---|
| 1 | 1 → ga | 4 ← ga | 5 → ga | 2 → ga | 6 → ga | 3 → ga | 7 → ga | -66011.9 | 692.9 |
| 2 | 1 → ga | 4 ← ga | 5 → ga | 2 → ga | 6 → dn | 3 → ga | 7 → ga | -66007.6 | 693.0 |
| 3 | 1 → ga | 4 ← ga | 5 → dn | 2 → ga | 6 → ga | 3 → ga | 7 → ga | -66005.9 | 692.9 |
| 4 | 1 → ga | 4 ← ga | 5 → dn | 2 → ga | 6 → dn | 3 → ga | 7 → ga | ***-66001.7*** | ***693.0*** |

D

| Procedure | Log-Likelihood | Length (cM) |
|---|---|---|
| genome-assisted | -66001.7 | 693.0 |
| *de novo* | -81639.3 | 933.9 |

**Figure 4.** Schematic overview of the map order improvement using *I. trifida* reference genome in LG 7. A) Scatter plot of the physical distance in *I. trifida* reference genome (horizontal axis) versus the genetic distance in the *de novo* genetic map (colored dots) and the genome-assisted genetic map (dark blue dots). The *de novo* and the genome-assisted maps and their recombination fraction matrices are presented to the left and to the right hand side of the scatter plot, respectively. Using visual inspection, we detected seven distinct collinearity blocks (separated by vertical dashed lines) in the *de novo* scatter plot. We reconstructed sub-maps for each collinearity block using both *de novo* and reference genome orders. The multilocus log-likelihood and the final length of each collinearity block is presented in panel B. We noticed only two small blocks where the *de novo* order (5 and 6) yield a superior log-likelihood when compared with the genome-assisted orders. We then tested four complete map models concatenating all seven collinearity blocks (panel C). For all models, based in visual inspection, we proposed the same order of the blocks (1,4,5,2,6,3 and 7, also indicated in panel A by the red arrows connecting segments). The orientation of the segments, presented in panel C by the black arrows, were also obtained by visual inspection. Right arrows (→) indicate collinearity blocks with the same orientation of the *I. trifida* genome (forward); left arrows (←) indicate collinearity blocks with the opposite orientation of the *I. trifida* genome (backward); Except for blocks 5 and 6, where the *de novo* (dn) order yielded higher log-likelihoods, we used the genomic-assisted order (ga) within the other blocks. We tested both *de novo* and genomic orders for blocks 5 and 6, resulting in four different map models. Their final log-likelihood and length indicate that model 4 was the best between the models. Panel D shows a final comparison of the map obtained by using the *de novo* and the genome-assisted map, indicating a clear superiority of the latter

conditional probability distribution of $\mathcal{G}_{k,j}$ is defined as

$$\begin{aligned} \gamma_k(j) &= \Pr(\mathcal{G}_{k,j} \mid O_1, \cdots, O_z, \mathbf{r}, \Phi_{P_1}, \Phi_{P_2}) \\ &= \Pr(\mathcal{G}_{k,j} \mid O_1, \cdots, O_z, \lambda) \\ &= \frac{\alpha_k(j)\beta_k(j)}{\sum\limits_{i=1}^{400} \alpha_k(i)\beta_k(i)} \end{aligned} \tag{1}$$

where $\gamma_k(j)$ denotes the probability distribution of the genotype $\mathcal{G}_{k,j}$, given the sequence of observation of $z$ markers, denoted by $O_1, \cdots, O_z$ and the model parameters $\lambda = \{\mathbf{r}, \Phi_{P_1}, \Phi_{P_2}\}$, where $\mathbf{r}$ is a vector containing the sequence of recombination fractions between all markers and $\Phi_{P_1}$ and $\Phi_{P_2}$ are the linkage phase configurations in parents $P_1$ and $P_2$, respectively ('Beauregard' and 'Tanzania'). $\alpha_k(j) = \Pr(O_1, \cdots, O_k; \mathcal{G}_{k,j} \mid \mathbf{r}, \Phi_{P_1}, \Phi_{P_2})$ denotes the joint probability of the partial observation sequence to the left of marker $k$ (including $k$) and genotype $\mathcal{G}_{k,j}$, given the model parameters $\lambda$. Similarly, $\beta_k(j)$ denotes the probability of the partial observation sequence to the right of the position $k$ $(O_{k+1}, \cdots, O_z)$, given the genotype $\mathcal{G}_{k,j}$ and the model parameters $\lambda$. The quantities $\alpha_k(j)$ and $\beta_k(j)$ can be obtained using the classical *forward-backward* algorithm[13] (details in Section 2).

The probability that an individual carries a specific homolog at position $k$ can be obtained using

$$\Pr(\mathcal{H}_k \mid O_1, \cdots, O_z, \lambda) = \sum_{j=1}^{400} \Pr(\mathcal{H}_k \mid \mathcal{G}_{k,j}, O_1, \cdots, O_z, \lambda) \Pr(\mathcal{G}_{k,j} \mid O_1, \cdots, O_z, \lambda) \tag{2}$$

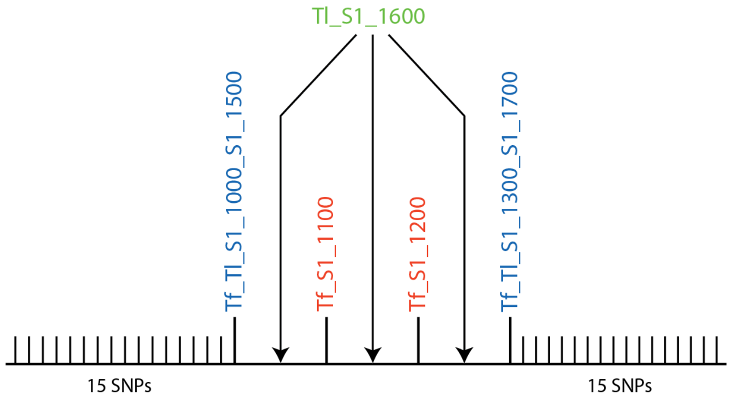

**Figure 5.** Insertion of a hypothetical private *I. triloba* SNP, *Tl_S1_1600*, using the genomic position constrains imposed by SNPs shared by both genomes. SNP *Tl_S1_1600* is located at 1600 basepairs in chromosome 1 of *I. triloba* genome and it will be positioned between SNPs *Tf_Tl_S1_1000_S1_1500* and *Tf_Tl_S1_1300_S1_1700*, already positioned in the framework map. These SNPs are located at 1500 and 1700 basepairs in the *I. triloba* genome. SNPs *Tf_S1_1100* and *Tf_S1_1200* were already positioned in the framework map, thus there are three possible intervals to position SNP *Tl_S1_1600*, indicated by the arrows. The SNP is positioned in the interval that yields the highest log-likelihood of a sub-map containing 15 SNPs to both sides of the shared SNPs. Also, SNPs that cause a map expansion greater than the maximum inter-marker distance in the original sub-map or needed to be tested in more than 100 intervals are not positioned into the map

where, $\mathcal{H}_k \in \{a, b, c, d, e, f, g, h, i, j, k, l\}$ is the inherited homolog at locus $k$, $\Pr(\mathcal{H}_k \mid \mathcal{G}_{k,j}, O_1, \cdots, O_z, \lambda) = 1$ if $\mathcal{H}_k \in \mathcal{G}_{k,j}$, 0 otherwise. We obtained the haplotype probability profile for all 15 homology groups (one curve for each homologs, from $a$ to $l$) for all individual in the bi-parental cross by computing $\Pr(\mathcal{H}_k \mid O_1, \cdots, O_z, \lambda)$ at every marker across the genome.

#### 1.6 Heuristic algorithm to re-estimate haplotype probabilities and detect crossing-over events

Given the final map, a small proportion of the individuals displayed abnormal fluctuations in the haplotype probability profiles for specific genomic regions. These profiles were most likely caused by genotyping errors that were not captured by the filtering processes and can impair the detection of crossing-over points. To recognize and re-estimate these segments, we proposed the following heuristic algorithm:

1. Regions with haplotype probabilities greater than 0.8 are assumed to be 1.0, otherwise 0.0, forming a binary profile;
2. SNPs within regions smaller than 10 cM are removed. These regions are defined as a continuous segment of homolog or gaps flanked by crossing-overs. In the binary profile these regions are represented by a sequence of ones (segments) or zeros (gaps).
3. If the remaining SNPs represent 20% or more of all SNPs in the LG, use Eq. 1 to re-estimate the 400 genotypes across the whole LG and compute a new homolog probability profile. Otherwise, consider the probability profile *inconclusive*. At this point, the erratic probabilities should be resolved.
4. The crossing-over points are assessed by checking the points of probability transition across the LG. Also, the homologs involved in the chromosomal exchange can be trivially assessed.
5. Exchange points closer than 0.5 cM are considered *inconclusive*, since the haplotypes involved in the exchange could be erroneously assigned due to the lack of resolution in the mapping population.

We applied this procedure to the 15 LGs of all individuals in the population. Threshold values were tested through a grid search. The chosen values appeared to be effective in capturing recombination phenomenon adequately. Notice that this heuristic algorithm will not disturb well estimated genotype profiles, such as the one presented in Supplementary Figure 6 A. We also present an interactive version of the heuristic algorithm (https://gt4sp-genetic-map.shinyapps.io/offspring_haplotype_BT_population/) where the reader can choose new values for all thresholds aforementioned.

#### 1.7 Preferential pairing

In polyploids, homologs can be arranged in several configurations during synapsis in prophase I[14]. Regardless their complete configuration (involving bivalents or multivalents), chromosomal segments always are arranged in pairs due to the synaptonemal complex properties[15]. In hexaploids, as presented by[10], there are always 15 possible pairing configurations. Let $\Psi = \{\psi_i\}$, $i = 1, \cdots, 15$ denote a set containing all 15 possible pairing configurations (Table 5).

**Table 5.** Homolog pairing configurations during prophase I for parents 'Beauregard' and 'Tanzania'

| $\Psi$ | 'Beauregard' | 'Tanzania' |
|---|---|---|
| $\psi_1$ | ab/cd/ef | gh/ij/kl |
| $\psi_2$ | ab/ce/df | gh/ik/jl |
| $\psi_3$ | ab/cf/de | gh/il/jk |
| $\psi_4$ | ac/bd/ef | gi/hj/kl |
| $\psi_5$ | ac/be/df | gi/hk/jl |
| $\psi_6$ | ac/bf/de | gi/hl/jk |
| $\psi_7$ | ad/bc/ef | gj/hi/kl |
| $\psi_8$ | ad/be/cf | gj/hk/il |
| $\psi_9$ | ad/bf/ce | gj/hl/ik |
| $\psi_{10}$ | ae/bc/df | gk/hi/jl |
| $\psi_{11}$ | ae/bd/cf | gk/hj/il |
| $\psi_{12}$ | ae/bf/cd | gk/hl/ij |
| $\psi_{13}$ | af/bc/de | gl/hi/jk |
| $\psi_{14}$ | af/bd/ce | gl/hj/ik |
| $\psi_{15}$ | af/be/cd | gl/hk/ij |

Under random pairing model, these configurations are equally likely. Although we use this assumption to construct the genetic map, it is possible to compute the posterior probability distribution of the pairing configurations at any position $k$ in the genome using

$$\Pr(\psi_i \mid O_1, \cdots, O_z, \lambda)_k = \frac{1}{n} \sum_{l=1}^{n} \sum_{j=1}^{400} \Pr(\psi_i \mid \mathcal{G}_{k,j}, O_1, \cdots, O_z, \lambda) \Pr(\mathcal{G}_{k,j} \mid O_1, \cdots, O_z, \lambda)_l \tag{3}$$

where $O_1, \cdots, O_z$ denotes the complete sequence of observations and $\lambda$ denotes parameter set of the multilocus map, $n$ is the number of individuals in the population, $\Pr(\psi_i \mid \mathcal{G}_{k,j}, O_1, \cdots, O_z, \lambda) = \left(\frac{m}{2}!\right)^{-1}$ ($m = 6$ for hexaploids) if $\mathcal{G}_{k,j}$ is consistent with $\psi_i$, i.e., if genotype $\mathcal{G}_{k,j}$ can be originated from the pairing configuration $\psi_i$, 0 otherwise (see[10] for details) and $\Pr(\mathcal{G}_{k,j} \mid O_1, \cdots, O_z, \lambda)_l$ is obtained for individual $l$ using Eq. 1. To test whether the observed frequencies of the 15 bivalent configurations differ from the expected under random pairing $\frac{1}{15}$, we used the $\chi^2$ test with $P < 0.001$ to declare significance.

**Using single-dose markers to assess preferential pairing** Single-dose markers will segregate in a 1:1 fashion regardless the ploidy type or level of the analyzed species[16]. In this case, markers are either in coupling, if they are in the same homolog, or repulsion, if they are in different homologs. It is well established that, for autopolyploids, closely linked markers in repulsion carry substantially less information about the recombination fraction when compared to coupling markers at the same distance[17,18,10], while in allopolyploids markers linked in repulsion phase will carry higher amount of information when compared to autopolyploids. Thus, we performed an additional preferential pairing assessment based only on simplex markers which are less prone to genotyping errors and segregation distortion filtering. First we ordered all simplex markers from homologs $a$ to $f$ in parent 'Beauregard' and from homologs $g$ to $l$ in 'Tanzania' for each LG. Then, for each LG, we arranged the $LOD_{ph}$ for all simplex SNP pairwise combinations in a matrix from which we generated a heatmap. For an autopolyploid genomes, we expect that high $LOD_{ph}$, will be concentrated within each homolog, generating a block-diagonal pattern matrix, while for allopolyploids, besides high $LOD_{ph}$ within homolog, we expect to observe high $LOD_{ph}$ between homologs, yielding off-diagonal patterns.

#### 2 Brief description of the hidden Markov model proposed by Mollinari and Garcia (2019)

In this section we briefly describe the hidden Markov model proposed by Mollinari and Garcia [10]. We derived the formulas for the autohexaploid case and, for simplicity, we omitted the inclusion of the probability distribution of the genotypes, but the complete derivation can be found in the original work.

**Definition of HMM elements:** According to Rabiner [13], three elements are needed to specify an HMM: the *transition probability distribution*, the *initial state distribution*, and the *observation symbol probability distribution*, also known as *emission probability distribution*. These elements are used recursively in the classical *forward-backward* algorithm in order to obtain the likelihood of the model and estimate the its parameters [19]. For an autohexaploid full-sib population, the transition probability is defined as

$$t_k(j, j') = \Pr(\mathcal{G}_{k+1,j'}|\mathcal{G}_{k,j}) = \frac{(1-r_k)^{6-l_{P_1}-l_{P_2}}(r_k)^{l_{P_1}+l_{P_2}}}{\binom{3}{l_{P_1}}\binom{3}{l_{P_2}}} \tag{4}$$

where $\mathcal{G}_{k,j}$ denotes the $j$-th genotype, $j \in \{1, \cdots, 400\}$, at position $k$ of a particular individual. $l_{P_1}$ and $l_{P_2}$ denote the number of recombination events between loci $k$ and $k+1$ in parents $P_1$ and $P_2$, ('Beauregard' and 'Tanzania' respectively) and $r_k$ is the recombination fraction between markers at position $k$ and $k+1$. The initial state distribution under the assumption of no preferential pairing is

$$\zeta_j = \Pr(\mathcal{G}_{1,j}) = \frac{1}{400}, \; j \in \{1, \cdots, 400\} \tag{5}$$

To derive the emission probability distribution, let $\varphi^k_{P_1}$ and $\varphi^k_{P_2}$ denote a subset of size 3 sampled from the 6 possible alleles in parents $P_1$ and $P_2$, respectively, at locus $k$. For two particular subsets $\varphi^k_{P_1}$ and $\varphi^k_{P_2}$ the 400 genotypic states in the full transition space can be associated to a dosage. The dosage associated to the $j$-th state is obtained by counting the number of alleles present in the intersection between the parental allelic set $(\varphi^k_{P_1} \cup \varphi^k_{P_2})$ and $\mathcal{G}_{k,j}$. For a biallelic marker, the emission function is defined in terms of the observed marker dosage $O$ as

$$b_j(O) = \Pr(O|\mathcal{G}_{k,j}, \varphi^k_{P_1}, \varphi^k_{P_2}) = \begin{cases} 1-\epsilon & \text{if} \quad O = \delta(k,j) \\ \frac{\epsilon}{m} & \text{otherwise} \end{cases} \tag{6}$$

where $\delta(k,j) = |(\varphi^k_{P_1} \cup \varphi^k_{P_2}) \cap \mathcal{G}_{k,j}|$, $|.|$ defines the cardinality of a set, and $\epsilon$ denotes the global genotype error rate.

**Computation of $\alpha_k(j)$, $\beta_k(j)$, and the HMM-based multilocus likelihood** Suppose there are $z$ markers in a hexaploid linkage group in a known order represented by $M_1, \cdots, M_k, \cdots, M_z$. Let $\mathbf{r} = (r_1, \cdots, r_k, \cdots, r_{z-1})$ denote the recombination fraction vector between all marker intervals in this sequence. Also, assume linkage phase configurations in parents $P_1$ and $P_2$ denoted respectively by $\Phi_{P_1} = (\varphi^1_{P_1}, \cdots, \varphi^k_{P_1}, \cdots, \varphi^z_{P_1})$ and $\Phi_{P_2} = (\varphi^1_{P_2}, \cdots, \varphi^k_{P_2}, \cdots, \varphi^z_{P_2})$. The sequence of observations for the

$z$ markers is denoted by $(O_1, \cdots, O_k, \cdots, O_z)$. The likelihood of $M_1, \cdots, M_k, \cdots, M_z$ can be obtained using Eqs (4), (5) and (6) following the classical *forward procedure* [13]. Let $\alpha_k(j) = \Pr(O_1, \cdots, O_k; \mathcal{G}_{k,j} \mid \mathbf{r}, \Phi_{P_1}, \Phi_{P_2})$ denote the probability of the partial observation sequence $(O_1, \cdots, O_k)$ and genotype $\mathcal{G}_{k,j}$ given the sequence of recombination fractions $\mathbf{r}$, and the linkage phase configurations $\Phi_{P_1}$ and $\Phi_{P_2}$. The forward procedure follows the steps below:

1. Initialization:
$$\alpha_1(j) = \zeta_j b_j(O_1),\ j = 1, \cdots, 400 \tag{7}$$

2. Induction:
$$\alpha_{k+1}(j') = \left[\sum_{j}^{400} \alpha_k(j)\, t_k(j, j')\right] b_{j'}(O_{k+1}) \tag{8}$$
where $\quad k = 1, \cdots, z-1$ and $j' = 1, \cdots, 400$

3. Termination:
$$\Pr(O_1, \cdots, O_z | \mathbf{r}, \Phi_{P_1}, \Phi_{P_2}) = \sum_{j=1}^{400} \alpha_z(j) \tag{9}$$

Then, the likelihood of the model is defined as

$$\prod_{i=1}^{n} \Pr(O_{1,i}, \cdots, O_{z,i} | \mathbf{r}, \Phi_{P_1}, \Phi_{P_2}) \tag{10}$$

where $n$ is the number of individuals in the full-sib population, $O_{1,i}, \cdots, O_{z,i}$ is the sequence of marker observations for individual $i$. For the backward procedure, consider the variable $\beta_k(j) = \Pr(O_{k+1}, \cdots, O_z \mid \mathcal{G}_{k,j}, \mathbf{r}, \Phi_{P_1}, \Phi_{P_2})$ as the probability of the partial observation sequence from $k+1$ to $z$, given the genotype $\mathcal{G}_{k,j}$, the recombination fraction vector $\mathbf{r}$, and the linkage phase configurations $\Phi_{P_1}$ and $\Phi_{P_2}$. The solution to $\beta_k(j)$ was also described by[13] as follows:

1. Initialization:
$$\beta_z(j) = 1,\ j = 1, \cdots, 400 \tag{11}$$

2. Induction:
$$\beta_k(j) = \sum_{j'}^{400} t_k(j, j') b_{j'}(O_{k+1}) \beta_{k+1}(j') \tag{12}$$
where $\quad k = z-1, z-2, \cdots, 1$ and $j = 1, \cdots, 400$

With $\alpha_k(j)$ and $\beta_k(j)$ in hands for any $k$ position, we can compute the probability distribution underling the 400 possible genotypes in the full-sib population using Eq. 1 described in Section Genotype probabilities and offspring haplotype reconstruction.

3. Shan Wu, Kin H. Lau, Qinghe Cao, John P. Hamilton, Honghe Sun, Chenxi Zhou, Lauren Eserman, Dorcus C. Gemenet, Bode A. Olukolu, Haiyan Wang, Emily Crisovan, Grant T. Godden, Chen Jiao, Xin Wang, Mercy Kitavi, Norma Manrique-Carpintero, Brieanne Vaillancourt, Krystle Wiegert-Rininger, Xinsun Yang, Kan Bao, Jennifer Schaff, Jan Kreuze, Wolfgang Gruneberg, Awais Khan, Marc Ghislain, Daifu Ma, Jiming Jiang, Robert O. M. Mwanga, Jim Leebens-Mack, Lachlan J. M. Coin, G. Craig Yencho, C. Robin Buell, and Zhangjun Fei. Genome sequences of two diploid wild relatives of cultivated sweetpotato reveal targets for genetic improvement. *Nat Commun*, 9(1), 2018. doi: 10.1038/s41467-018-06983-8.

4. Brant C Faircloth and Travis C Glenn. Not all sequence tags are created equal: designing and validating sequence identification tags robust to indels. *PloS one*, 7(8):e42543, 2012. ISSN 1932-6203.

5. Phillip A. Wadl, Bode A. Olukolu, Sandra E. Branham, Robert L. Jarret, G. Craig Yencho, and D. Michael Jackson. Genetic Diversity and Population Structure of the USDA Sweetpotato (Ipomoea batatas) Germplasm Collections Using GBSpoly. *Frontiers in Plant Science*, 9(August):1–13, 2018.

6. Jj Doyle and Jl Doyle. A rapid dna isolation procedure from small quantities of fresh leaf tissues. *Phytochem. Bull.*, 19, 11 1987.

7. Oliver Serang, Marcelo Mollinari, and Antonio AF Garcia. Efficient exact maximum a posteriori computation for bayesian snp genotyping in polyploids. *PLoS One*, 7(2):e30906, 2012.

8. Antonio A. F. Garcia, Marcelo Mollinari, Thiago G. Marconi, Oliver R. Serang, Renato R. Silva, Maria L. C. Vieira, Renato Vicentini, Estela A. Costa, Melina C. Mancini, Melissa O. S. Garcia, Maria M. Pastina, Rodrigo Gazaffi, Eliana R. F. Martins, Nair Dahmer, Danilo A. Sforça, Claudio B. C. Silva, Peter Bundock, Robert J. Henry, Glaucia M. Souza, Marie-Anne van Sluys, Marcos G. A. Landell, Monalisa S. Carneiro, Michel A. G. Vincentz, Luciana R. Pinto, Roland Vencovsky, and Anete P. Souza. SNP genotyping allows an in-depth characterisation of the genome of sugarcane and other complex autopolyploids. *Sci Rep*, 3(1), 2013. doi: 10.1038/srep03399.

9. Marcelo Mollinari and Oliver Serang. *Plant genotyping: methods and protocols*, chapter Quantitative SNP Genotyping of Polyploids with MassARRAY and Other Platforms, pages 215–241. Springer, 2015. doi: 10.1007/978-1-4939-1966-6_17.

10. Marcelo Mollinari and Antonio AF Garcia. Linkage analysis and haplotype phasing in experimental autopolyploid populations with high ploidy level using hidden markov models. *G3*, -(Under review), 2019.

### 3 Supplementary Figures

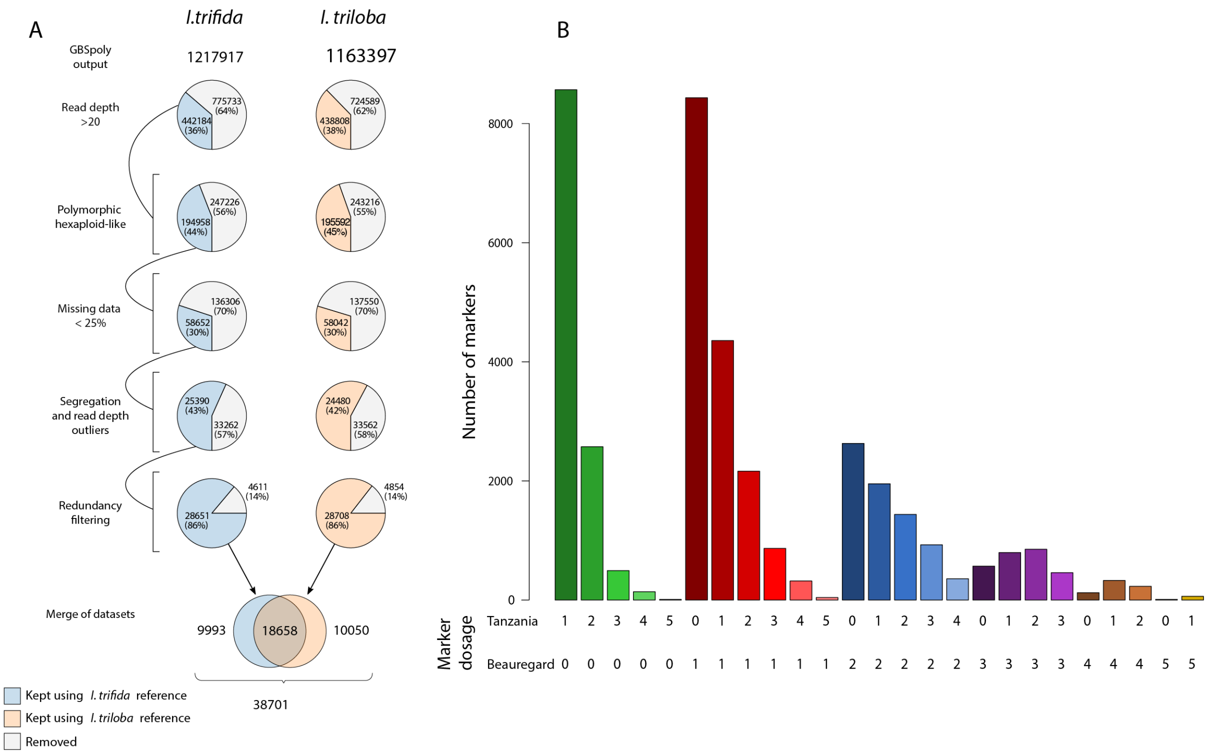

**Supplementary Figure 1.** SNP filtering process and resulting dosage distribution for 'Beauregard' $\times$ 'Tanzania' population. (A) Filtering process considering *I. trifida* and *I. triloba* reference genomes. The first step consisted of removing SNPs with less than 20 reads on average. In the next step, we removed monomorphic SNPs and also those classified by SuperMASSA as diploids and tetraploids. We then removed SNPs with more than 25% of missing data. We considered a data point as missing if the highest posterior probability was smaller than 0.8. A classical chi-square test for Mendelian segregation was performed in the remaining SNPs ($P < 5 \times 10^{-4}$). We also removed SNPs with more than 1.5$\times$ interquartile range across the offspring. Finally, we filtered out SNPs with redundant information resulting in 33320 SNPs anchored to *I. trifida* reference and 33603 anchored to *I. triloba*. Both data sets were merged totaling 38701 high-quality SNPs. (B) Distribution of the marker doses in 'Beauregard' and 'Tanzania' genotypes. Among the 38701 high-quality scored SNPs, 55.5% were classified as simplex or double-simplex ($0 \times 1$, $1 \times 0$, $1 \times 1$, $0 \times 5$, $5 \times 5$, $1 \times 5$ and $5 \times 1$) and 44.5% were classified as multiplex.

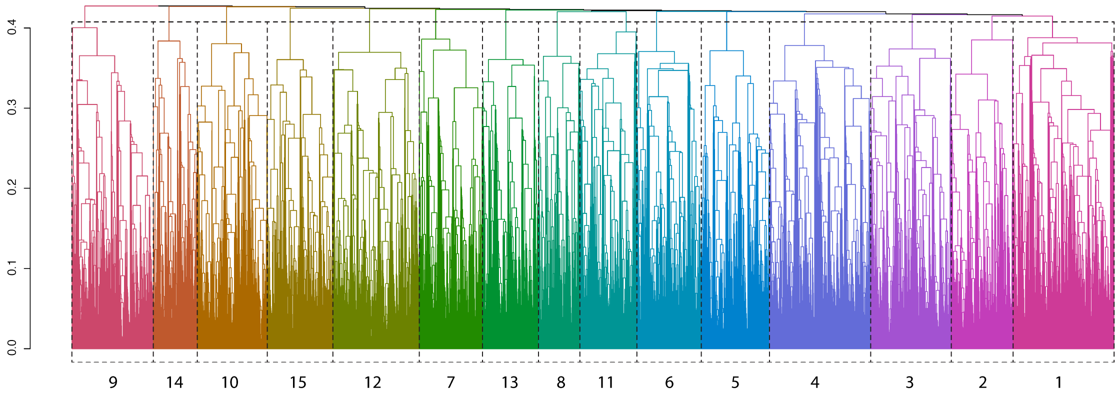

**Supplementary Figure 2.** Dendrogram comprising 15 LGs obtained using UPGMA clustering method. Numbers bellow the dendrogram indicate the correspondence between the groups found and the *I. trifida* and *I. triloba* chromosomes.

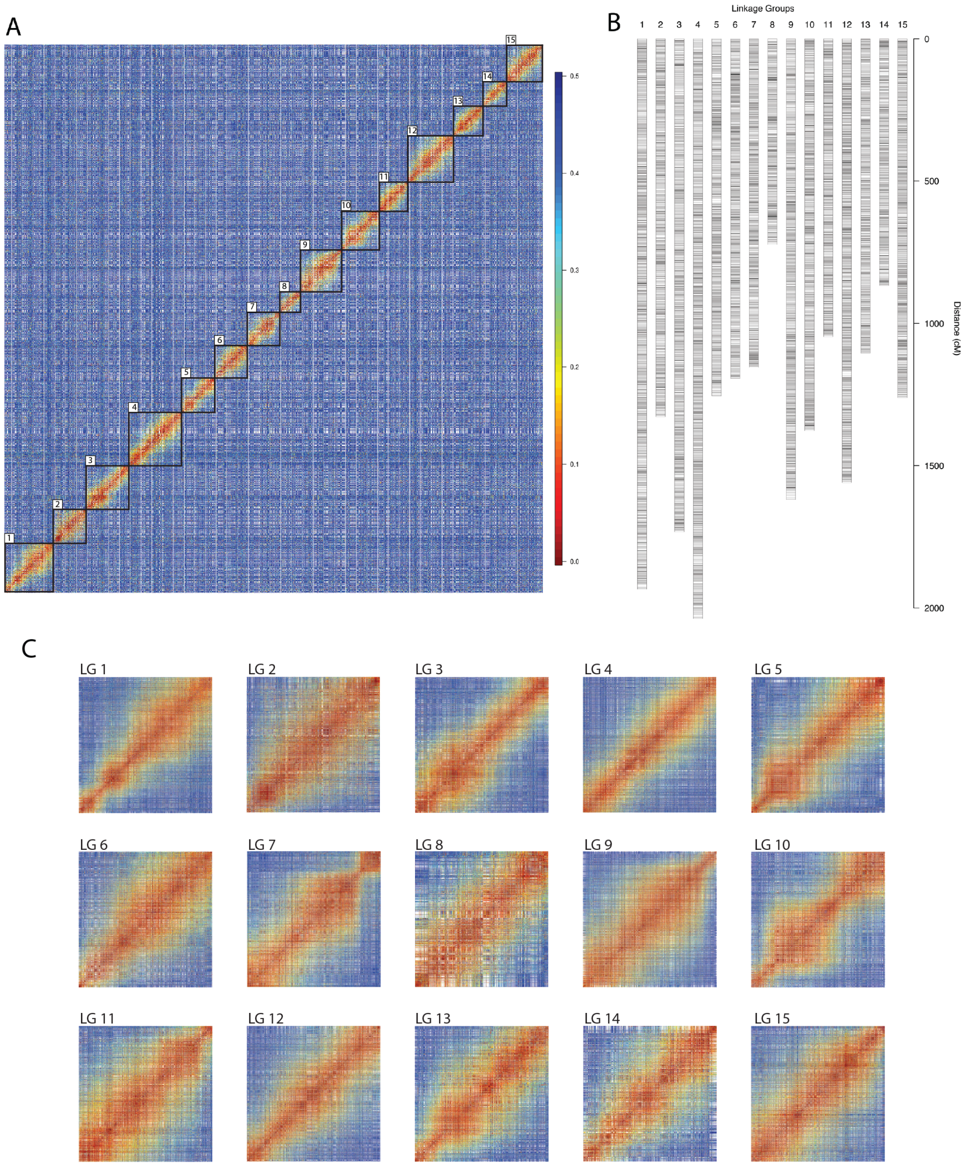

**Supplementary Figure 3.** Recombination fraction matrix and resulting multipoint *de novo* genetic map. (A) Recombination fraction between 32,200 SNPs positioned in the "de novo" map. The 15 monotonous sub-matrices on the diagonal of the block-diagonal matrix represent the ordered linkage groups (LGs) of *I. batatas*. (B) "De novo" genetic map with no with no considerable gaps between markers. (C) Isolated recombination fraction matrices for each LG with rather evident monotonous patterns.

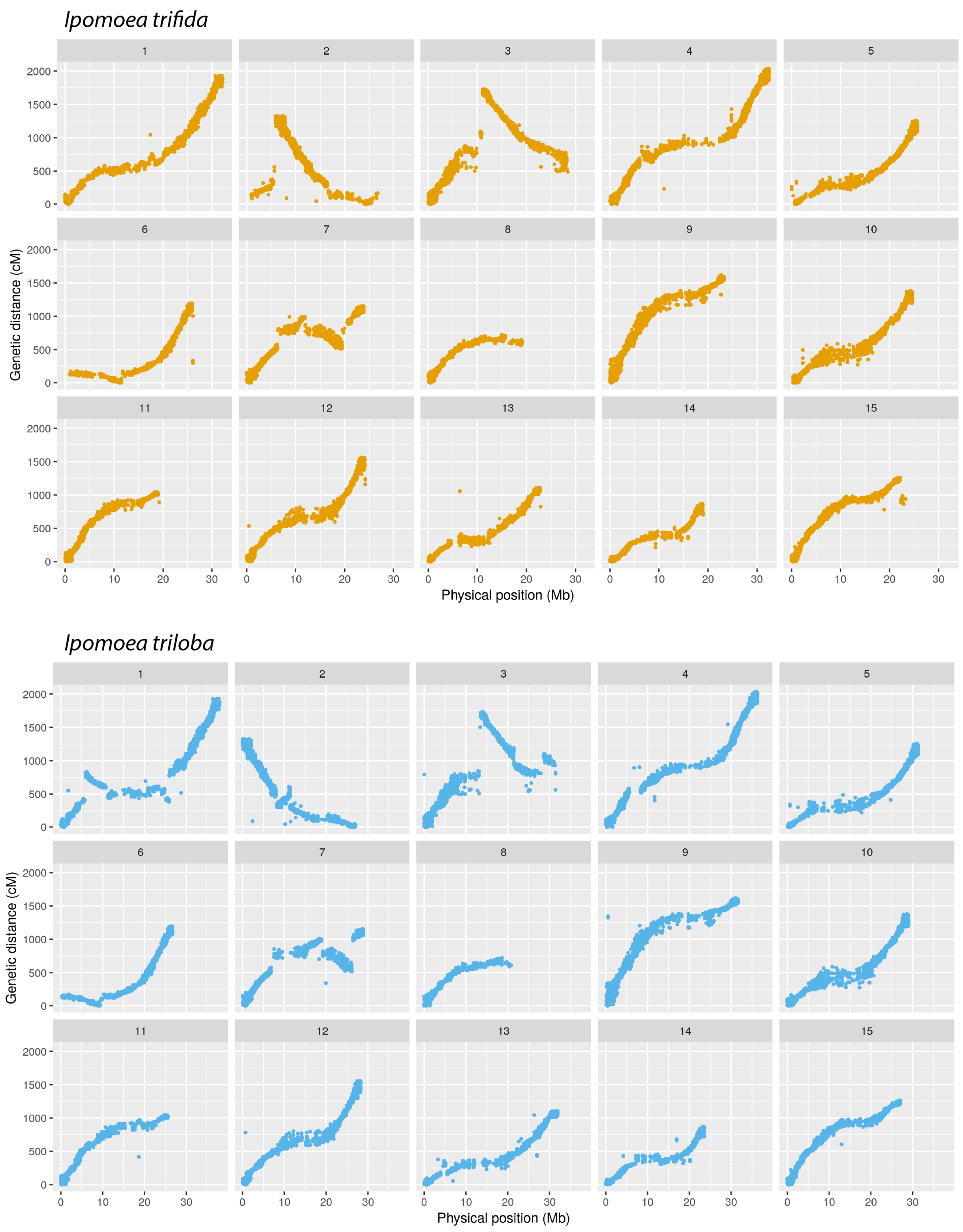

**Supplementary Figure 4.** Physical distance in *I. trifida* and *I. triloba* reference genomes versus genetic distance in the 15 linkage groups of the sweetpotato using *de novo* ordering.

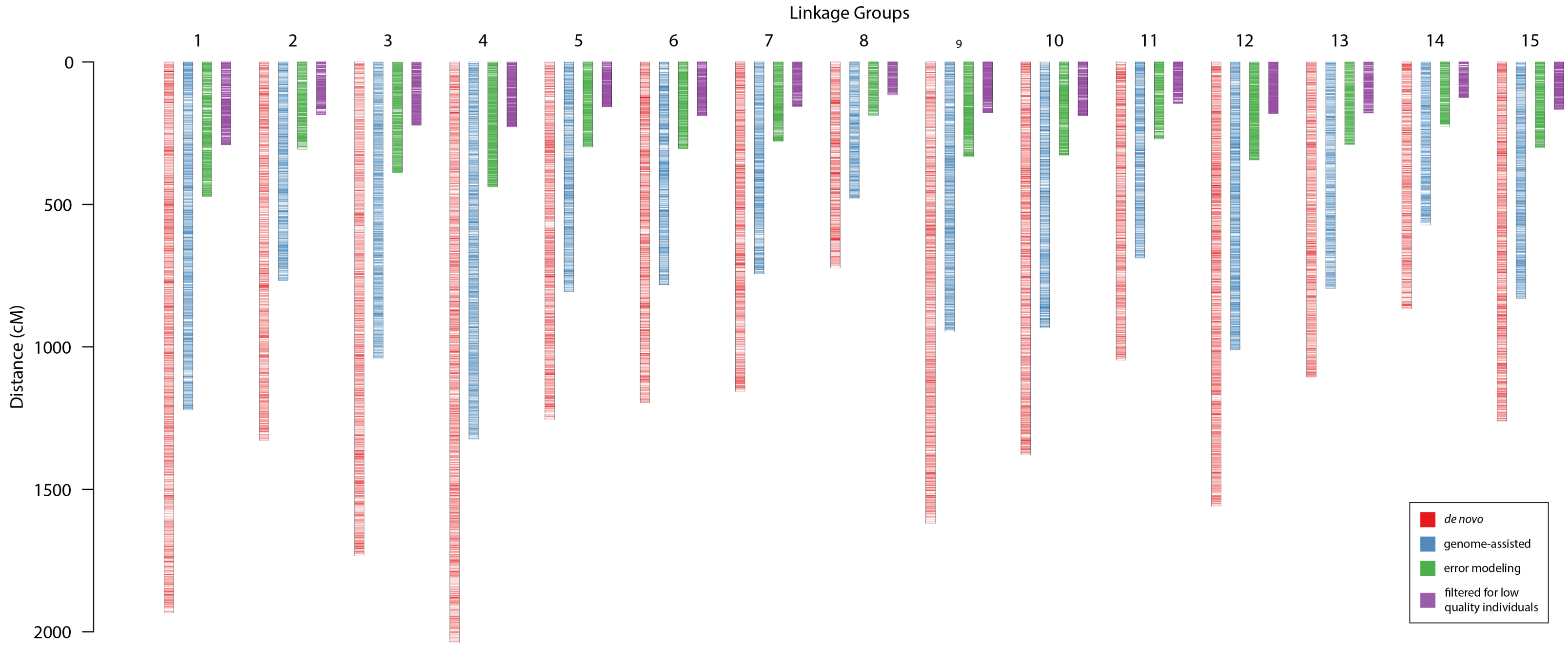

**Supplementary Figure 5.** Comparison between maps using three levels of improvement. The red map shows the *de novo* order using the MDS algorithm, with a total map length of 20,201.8 cM, 32,200 SNPs, and average inter-locus distance of $\approx 0.63$ cM. The blue map was obtained using the genome-assisted ordering procedure, resulting in a map with 30,723 SNPs spanning 12,937.3 cM with average inter-locus distance of $\approx 0.42$ cM. In the green map, the markers are in the same order and linkage phase as the blue map, however the recombination fractions were re-estimated using the probability distribution of the marker dosage genotypes provided by SuperMASSA as a prior information in the hidden Markov model. This procedure reduced the size of the map to 4,764.1 cM with average inter-locus distance of $\approx 0.16$ cM. Finally, the purple map was obtained removing problematic individuals with disrupted genotype profiles across all linkage groups, which could be caused by poor DNA amplification or sequencing errors. The final map contains 30,684 SNPs spanning 2,708.4 cM (average inter-locus distance of $\approx 0.09$ cM), with $\approx 60.7\%$ of the markers simplex and double-simplex, and 39.3% multiplex. In all cases, the linkage phase inference and the estimation of recombination fractions were obtained using the method proposed to Mollinari and Garcia[10]

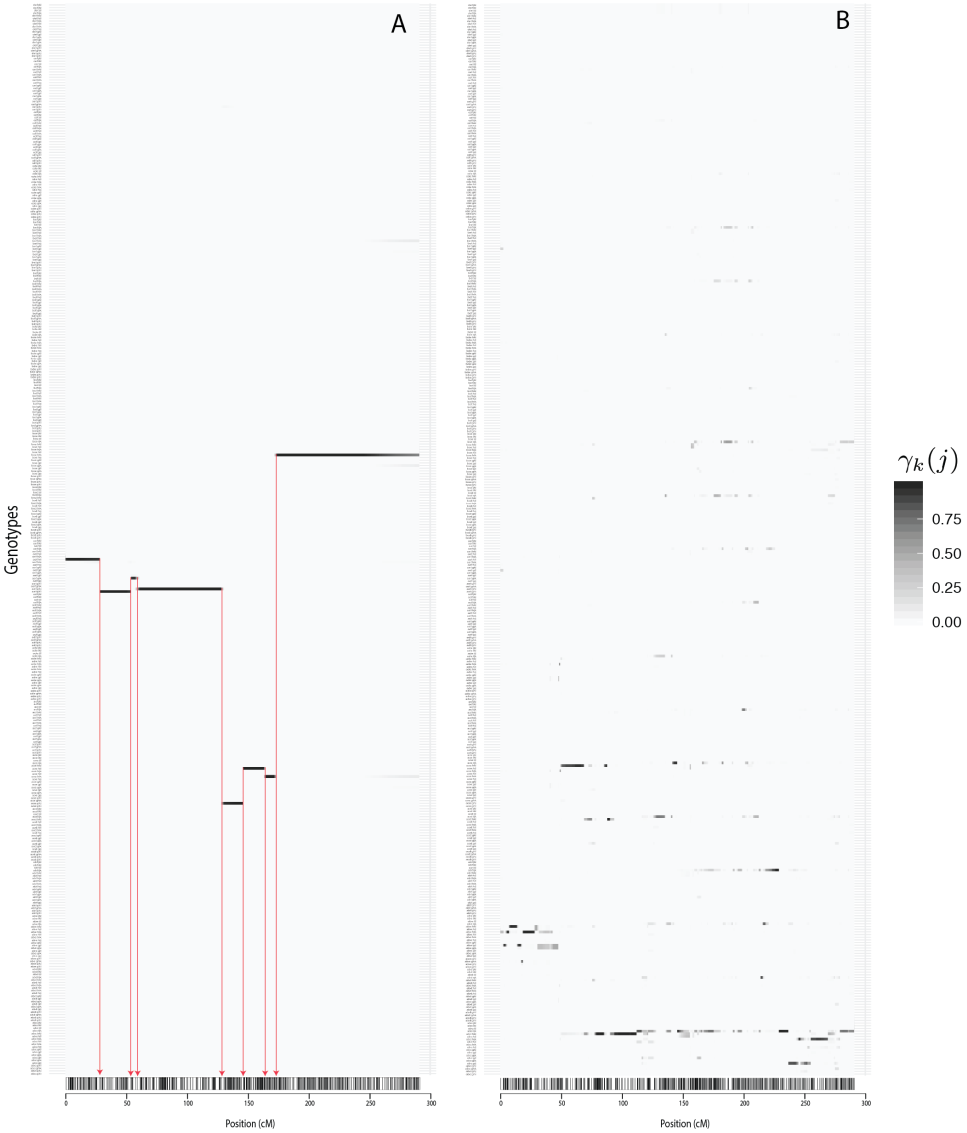

**Supplementary Figure 6.** Comparison between conditional probability profiles. A) Probability profile of individual *BT05.320*. This individual represents an ideal scenario with high probabilities associated with specific genotypes out of 400 possible along LG 1. The six abrupt breakages in the probability profile indicate crossing-over points (red arrows). B) Erratic probability profile of individual *BT13.359*, with probabilities are spread across several genotypic classes, and the recombination points cannot be trivially assessed.

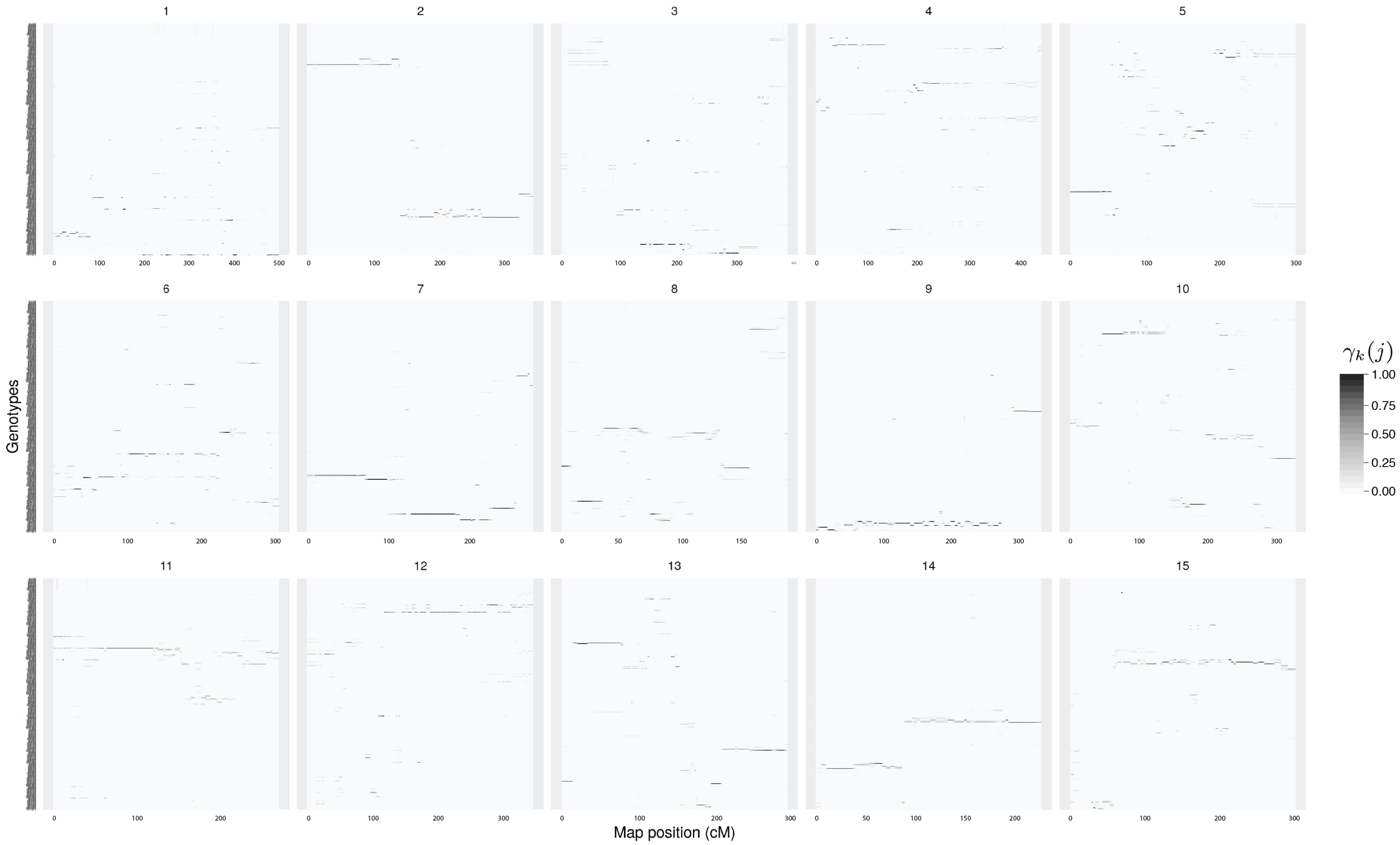

**Supplementary Figure 7.** Erratic probability profile of individual *BT13.359* across 15 LGs.

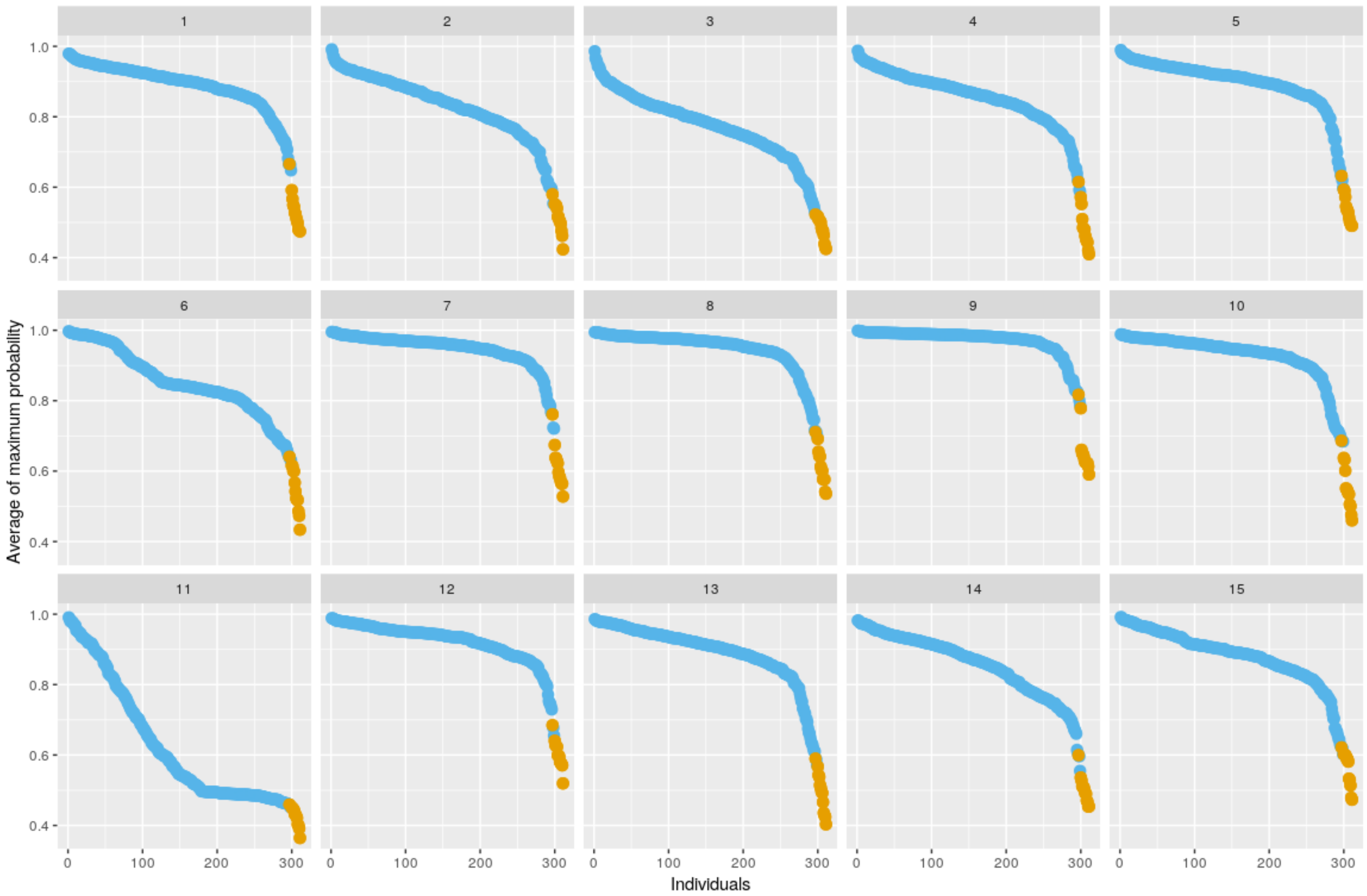

**Supplementary Figure 8.** Distribution of the average of the maximum genotypic probability across all linkage groups for all individuals. For each individual, we obtained $\frac{1}{N}\sum_{k}^{N} \max_j \gamma_k(j)$, where N is the number of positions in the considered LG. We selected individuals in the fifth percentile of the distribution for each LG and filtered out 13 individuals (orange dots) that consistently appeared in at least 10 LGs. Eliminated individuals: *BT05.358 BT13.197 BT13.237 BT05.323 BT05.324 BT13.049 BT13.061 BT13.063 BT13.135 BT13.196 BT13.153 BT13.172* and *BT13.359*

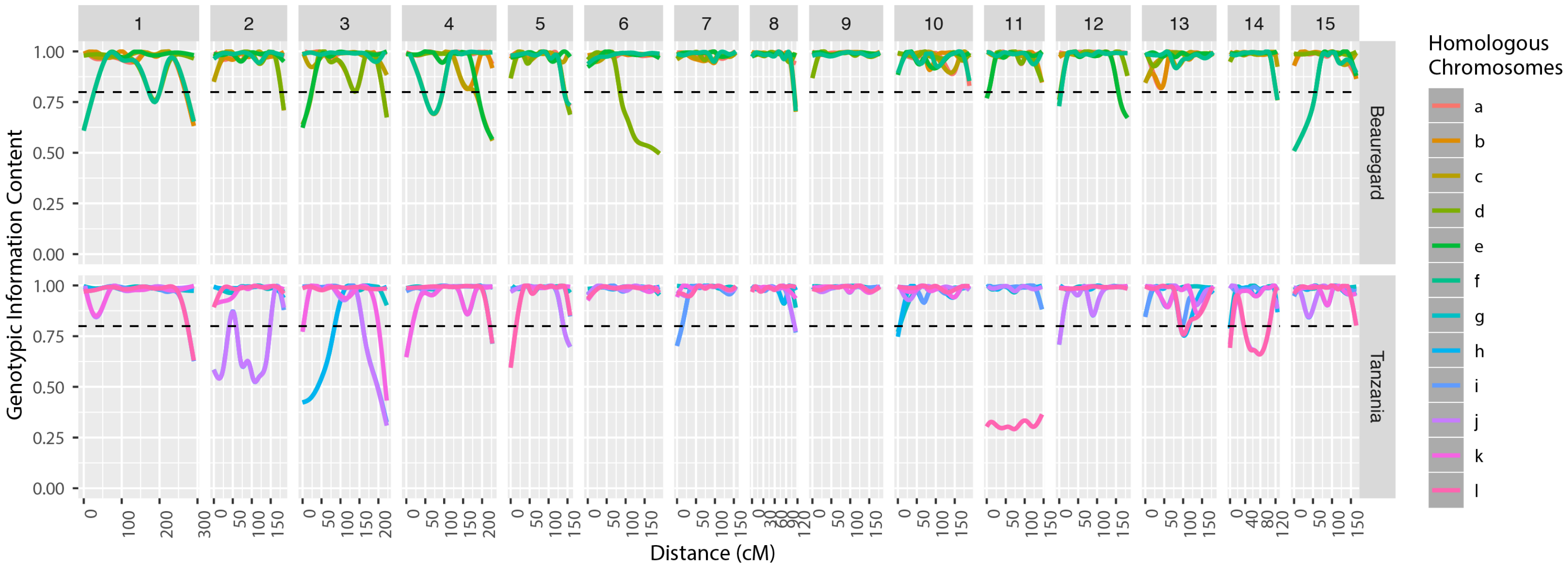

**Supplementary Figure 9.** Genotypic Information Content (GIC) for each homologs in parents 'Beauregard' and 'Tanzania' for all 15 linkage groups. The dashed line indicates $GIC = 0.8$.

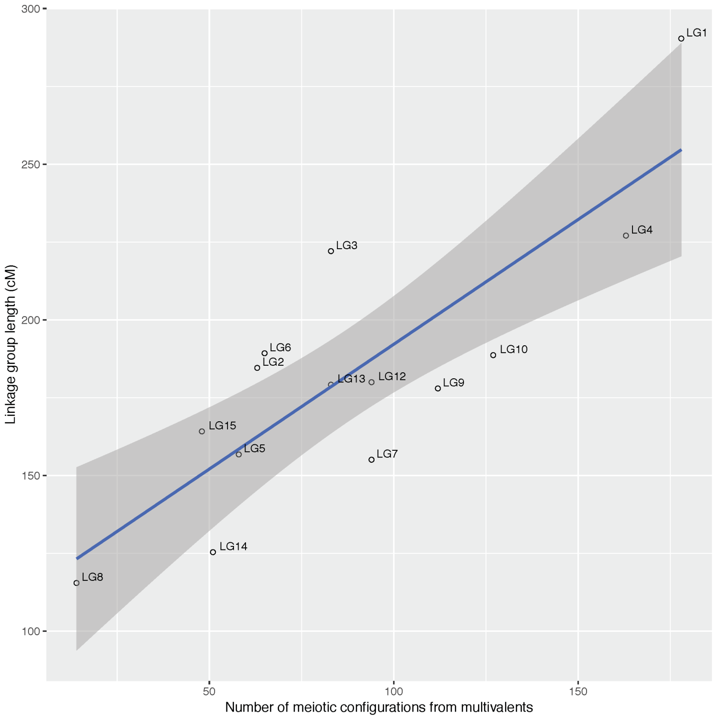

**Supplementary Figure 10.** Number of multivalent signatures versus linkage group size. It is possible to observe a linear relationship between multivalent signatures and LG size. The slope of the regression was 0.85 ($P < 10^{-3}$). LG 11 for parent 'Tanzania' was mostly inconclusive and was not considered.

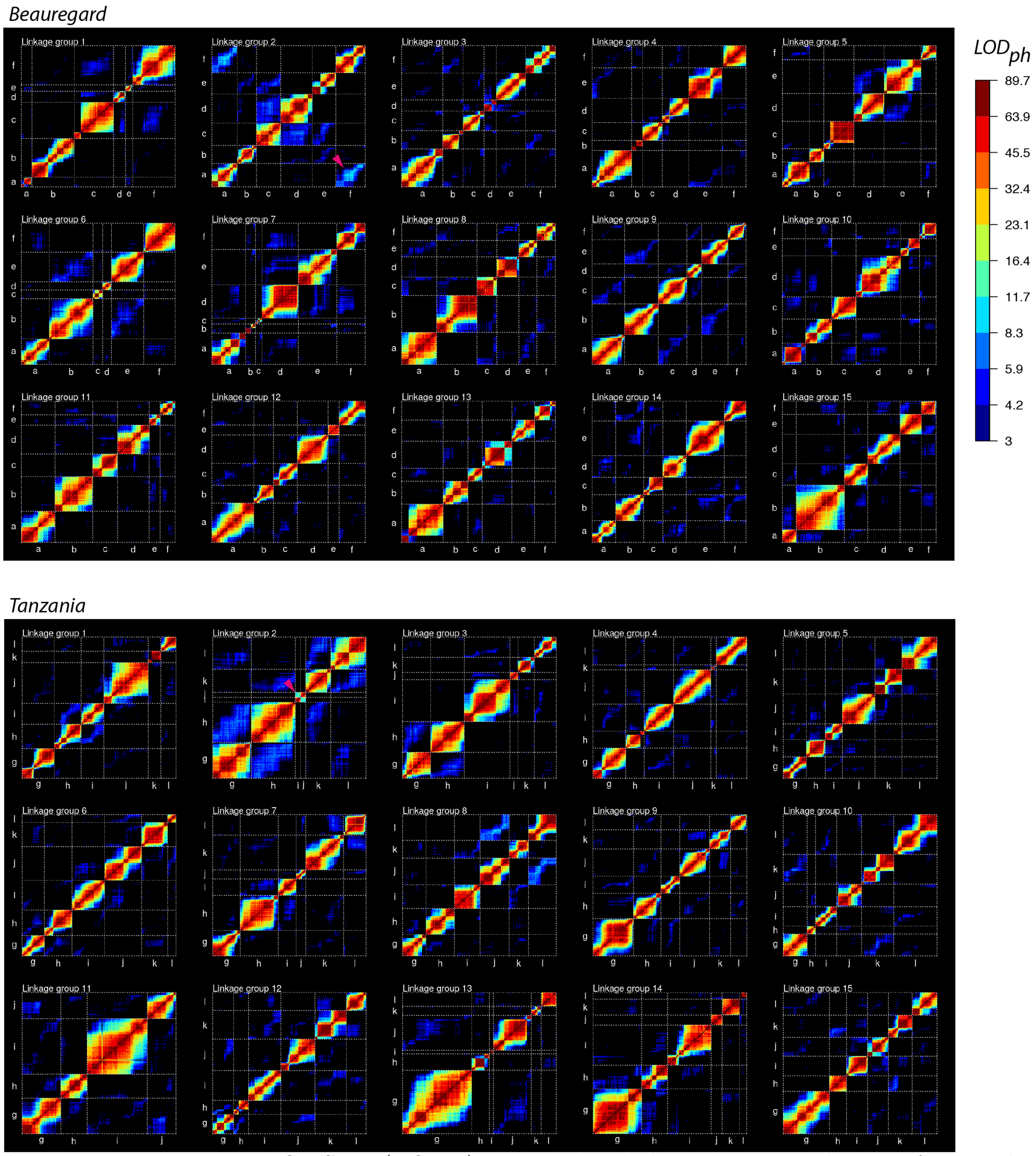

**Supplementary Figure 11.** LOD Score ($LOD_{ph}$) associated with the pairwise recombination fractions between simplex markers in parents 'Beauregard' and 'Tanzania'. We also addressed the question of preferential pairing using a more traditional approach proposed by [16]. For each LG, we isolated the simplex markers of each homolog, keeping their map order and arranged them from $a$ to $f$ in 'Beauregard' and from $g$ to $l$ in 'Tanzania'. Then, we obtained a heat map for the pairwise $LOD_{ph}$ between all simplex markers within each LG. Pairs with $LOD_{ph} < 3$ were removed from the heatmap. Within homologs we observe the monotonous pattern with $LOD_{ph}$ decreasing for markers positioned further away, as expected for simplex markers positioned in the same homolog. Between homologs, however, the levels of $LOD_{ph}$ are significantly smaller, indicating that simplex markers in repulsion carry less information than their coupling counterparts. This behavior is a characteristic of non-preferential pairing systems. In cases where preferential pairing exists, the of-diagonal blocks would contain higher values of $LOD_{ph}$, as showed by the pink arrowheads between homologs $a$ and $f$, and $i$ and $g$, in LG2, which confirm our findings using the whole chromosome estimation.

**Supplementary Table 1.** Number of SNPs identified using linkage analysis and physical location in *I. trifida* and *I. triloba* reference genomes.

| Homology group number | Chromosome number in *I. trifida* and *I. triloba* reference genomes | | | | | | | | | | | | | | |
|---|---|---|---|---|---|---|---|---|---|---|---|---|---|---|---|
| | 1 | 2 | 3 | 4 | 5 | 6 | 7 | 8 | 9 | 10 | 11 | 12 | 13 | 14 | 15 |
| 1 | 3182 | 63 | 59 | 32 | 55 | 33 | 34 | 10 | 38 | 30 | 34 | 35 | 20 | 14 | 31 |
| 2 | 20 | 2118 | 24 | 12 | 7 | 5 | 7 | 4 | 7 | 12 | 5 | 3 | 2 | 12 | 9 |
| 3 | 6 | 32 | 2687 | 26 | 25 | 21 | 21 | 22 | 22 | 11 | 10 | 17 | 7 | 22 | 21 |
| 4 | 8 | 18 | 39 | 3301 | 100 | 28 | 54 | 16 | 25 | 26 | 18 | 29 | 6 | 21 | 22 |
| 5 | 13 | 11 | 6 | 25 | 2219 | 27 | 29 | 23 | 26 | 20 | 19 | 22 | 12 | 6 | 13 |
| 6 | 24 | 7 | 7 | 21 | 28 | 2123 | 37 | 11 | 8 | 19 | 10 | 26 | 11 | 10 | 23 |
| 7 | 4 | 15 | 15 | 10 | 8 | 13 | 2101 | 23 | 42 | 9 | 16 | 5 | 28 | 2 | 7 |
| 8 | 9 | 10 | 11 | 9 | 8 | 7 | 8 | 1352 | 32 | 22 | 22 | 10 | 2 | 6 | 17 |
| 9 | 9 | 4 | 4 | 7 | 21 | 7 | 18 | 12 | 2827 | 16 | 14 | 18 | 3 | 16 | 19 |
| 10 | 7 | 15 | 8 | 2 | 13 | 9 | 7 | 12 | 11 | 2382 | 23 | 32 | 8 | 11 | 30 |
| 11 | 16 | 8 | 11 | 6 | 16 | 26 | 13 | 9 | 22 | 27 | 1781 | 50 | 31 | 17 | 29 |
| 12 | 3 | 5 | 7 | 13 | 14 | 17 | 13 | 5 | 14 | 10 | 22 | 2996 | 18 | 9 | 17 |
| 13 | 9 | 13 | 2 | 8 | 7 | 2 | 5 | 0 | 8 | 10 | 14 | 21 | 1862 | 27 | 26 |
| 14 | 0 | 2 | 3 | 9 | 5 | 5 | 0 | 9 | 3 | 8 | 3 | 8 | 11 | 1488 | 49 |
| 15 | 7 | 2 | 12 | 1 | 6 | 4 | 14 | 1 | 8 | 6 | 6 | 6 | 17 | 19 | 2315 |

**Supplementary Table 2.** Four hundred possible genotypes in a full-sib hexaploid population

| $j$ | $\mathcal{G}_{k,j}$ | $j$ | $\mathcal{G}_{k,j}$ | $j$ | $\mathcal{G}_{k,j}$ | $j$ | $\mathcal{G}_{k,j}$ | $j$ | $\mathcal{G}_{k,j}$ | $j$ | $\mathcal{G}_{k,j}$ | $j$ | $\mathcal{G}_{k,j}$ | $j$ | $\mathcal{G}_{k,j}$ | $j$ | $\mathcal{G}_{k,j}$ | $j$ | $\mathcal{G}_{k,j}$ |
|---|---|---|---|---|---|---|---|---|---|---|---|---|---|---|---|---|---|---|---|
| 1 | abc:ghi | 41 | abe:ghi | 81 | acd:ghi | 121 | acf:ghi | 161 | adf:ghi | 201 | bcd:ghi | 241 | bcf:ghi | 281 | bdf:ghi | 321 | cde:ghi | 361 | cef:ghi |
| 2 | abc:ghj | 42 | abe:ghj | 82 | acd:ghj | 122 | acf:ghj | 162 | adf:ghj | 202 | bcd:ghj | 242 | bcf:ghj | 282 | bdf:ghj | 322 | cde:ghj | 362 | cef:ghj |
| 3 | abc:ghk | 43 | abe:ghk | 83 | acd:ghk | 123 | acf:ghk | 163 | adf:ghk | 203 | bcd:ghk | 243 | bcf:ghk | 283 | bdf:ghk | 323 | cde:ghk | 363 | cef:ghk |
| 4 | abc:ghl | 44 | abe:ghl | 84 | acd:ghl | 124 | acf:ghl | 164 | adf:ghl | 204 | bcd:ghl | 244 | bcf:ghl | 284 | bdf:ghl | 324 | cde:ghl | 364 | cef:ghl |
| 5 | abc:gij | 45 | abe:gij | 85 | acd:gij | 125 | acf:gij | 165 | adf:gij | 205 | bcd:gij | 245 | bcf:gij | 285 | bdf:gij | 325 | cde:gij | 365 | cef:gij |
| 6 | abc:gik | 46 | abe:gik | 86 | acd:gik | 126 | acf:gik | 166 | adf:gik | 206 | bcd:gik | 246 | bcf:gik | 286 | bdf:gik | 326 | cde:gik | 366 | cef:gik |
| 7 | abc:gil | 47 | abe:gil | 87 | acd:gil | 127 | acf:gil | 167 | adf:gil | 207 | bcd:gil | 247 | bcf:gil | 287 | bdf:gil | 327 | cde:gil | 367 | cef:gil |
| 8 | abc:gjk | 48 | abe:gjk | 88 | acd:gjk | 128 | acf:gjk | 168 | adf:gjk | 208 | bcd:gjk | 248 | bcf:gjk | 288 | bdf:gjk | 328 | cde:gjk | 368 | cef:gjk |
| 9 | abc:gjl | 49 | abe:gjl | 89 | acd:gjl | 129 | acf:gjl | 169 | adf:gjl | 209 | bcd:gjl | 249 | bcf:gjl | 289 | bdf:gjl | 329 | cde:gjl | 369 | cef:gjl |
| 10 | abc:gkl | 50 | abe:gkl | 90 | acd:gkl | 130 | acf:gkl | 170 | adf:gkl | 210 | bcd:gkl | 250 | bcf:gkl | 290 | bdf:gkl | 330 | cde:gkl | 370 | cef:gkl |
| 11 | abc:hij | 51 | abe:hij | 91 | acd:hij | 131 | acf:hij | 171 | adf:hij | 211 | bcd:hij | 251 | bcf:hij | 291 | bdf:hij | 331 | cde:hij | 371 | cef:hij |
| 12 | abc:hik | 52 | abe:hik | 92 | acd:hik | 132 | acf:hik | 172 | adf:hik | 212 | bcd:hik | 252 | bcf:hik | 292 | bdf:hik | 332 | cde:hik | 372 | cef:hik |
| 13 | abc:hil | 53 | abe:hil | 93 | acd:hil | 133 | acf:hil | 173 | adf:hil | 213 | bcd:hil | 253 | bcf:hil | 293 | bdf:hil | 333 | cde:hil | 373 | cef:hil |
| 14 | abc:hjk | 54 | abe:hjk | 94 | acd:hjk | 134 | acf:hjk | 174 | adf:hjk | 214 | bcd:hjk | 254 | bcf:hjk | 294 | bdf:hjk | 334 | cde:hjk | 374 | cef:hjk |
| 15 | abc:hjl | 55 | abe:hjl | 95 | acd:hjl | 135 | acf:hjl | 175 | adf:hjl | 215 | bcd:hjl | 255 | bcf:hjl | 295 | bdf:hjl | 335 | cde:hjl | 375 | cef:hjl |
| 16 | abc:hkl | 56 | abe:hkl | 96 | acd:hkl | 136 | acf:hkl | 176 | adf:hkl | 216 | bcd:hkl | 256 | bcf:hkl | 296 | bdf:hkl | 336 | cde:hkl | 376 | cef:hkl |
| 17 | abc:ijk | 57 | abe:ijk | 97 | acd:ijk | 137 | acf:ijk | 177 | adf:ijk | 217 | bcd:ijk | 257 | bcf:ijk | 297 | bdf:ijk | 337 | cde:ijk | 377 | cef:ijk |
| 18 | abc:ijl | 58 | abe:ijl | 98 | acd:ijl | 138 | acf:ijl | 178 | adf:ijl | 218 | bcd:ijl | 258 | bcf:ijl | 298 | bdf:ijl | 338 | cde:ijl | 378 | cef:ijl |
| 19 | abc:ikl | 59 | abe:ikl | 99 | acd:ikl | 139 | acf:ikl | 179 | adf:ikl | 219 | bcd:ikl | 259 | bcf:ikl | 299 | bdf:ikl | 339 | cde:ikl | 379 | cef:ikl |
| 20 | abc:jkl | 60 | abe:jkl | 100 | acd:jkl | 140 | acf:jkl | 180 | adf:jkl | 220 | bcd:jkl | 260 | bcf:jkl | 300 | bdf:jkl | 340 | cde:jkl | 380 | cef:jkl |
| 21 | abd:ghi | 61 | abf:ghi | 101 | ace:ghi | 141 | ade:ghi | 181 | aef:ghi | 221 | bce:ghi | 261 | bde:ghi | 301 | bef:ghi | 341 | cdf:ghi | 381 | def:ghi |
| 22 | abd:ghj | 62 | abf:ghj | 102 | ace:ghj | 142 | ade:ghj | 182 | aef:ghj | 222 | bce:ghj | 262 | bde:ghj | 302 | bef:ghj | 342 | cdf:ghj | 382 | def:ghj |
| 23 | abd:ghk | 63 | abf:ghk | 103 | ace:ghk | 143 | ade:ghk | 183 | aef:ghk | 223 | bce:ghk | 263 | bde:ghk | 303 | bef:ghk | 343 | cdf:ghk | 383 | def:ghk |
| 24 | abd:ghl | 64 | abf:ghl | 104 | ace:ghl | 144 | ade:ghl | 184 | aef:ghl | 224 | bce:ghl | 264 | bde:ghl | 304 | bef:ghl | 344 | cdf:ghl | 384 | def:ghl |
| 25 | abd:gij | 65 | abf:gij | 105 | ace:gij | 145 | ade:gij | 185 | aef:gij | 225 | bce:gij | 265 | bde:gij | 305 | bef:gij | 345 | cdf:gij | 385 | def:gij |
| 26 | abd:gik | 66 | abf:gik | 106 | ace:gik | 146 | ade:gik | 186 | aef:gik | 226 | bce:gik | 266 | bde:gik | 306 | bef:gik | 346 | cdf:gik | 386 | def:gik |
| 27 | abd:gil | 67 | abf:gil | 107 | ace:gil | 147 | ade:gil | 187 | aef:gil | 227 | bce:gil | 267 | bde:gil | 307 | bef:gil | 347 | cdf:gil | 387 | def:gil |
| 28 | abd:gjk | 68 | abf:gjk | 108 | ace:gjk | 148 | ade:gjk | 188 | aef:gjk | 228 | bce:gjk | 268 | bde:gjk | 308 | bef:gjk | 348 | cdf:gjk | 388 | def:gjk |
| 29 | abd:gjl | 69 | abf:gjl | 109 | ace:gjl | 149 | ade:gjl | 189 | aef:gjl | 229 | bce:gjl | 269 | bde:gjl | 309 | bef:gjl | 349 | cdf:gjl | 389 | def:gjl |
| 30 | abd:gkl | 70 | abf:gkl | 110 | ace:gkl | 150 | ade:gkl | 190 | aef:gkl | 230 | bce:gkl | 270 | bde:gkl | 310 | bef:gkl | 350 | cdf:gkl | 390 | def:gkl |
| 31 | abd:hij | 71 | abf:hij | 111 | ace:hij | 151 | ade:hij | 191 | aef:hij | 231 | bce:hij | 271 | bde:hij | 311 | bef:hij | 351 | cdf:hij | 391 | def:hij |
| 32 | abd:hik | 72 | abf:hik | 112 | ace:hik | 152 | ade:hik | 192 | aef:hik | 232 | bce:hik | 272 | bde:hik | 312 | bef:hik | 352 | cdf:hik | 392 | def:hik |
| 33 | abd:hil | 73 | abf:hil | 113 | ace:hil | 153 | ade:hil | 193 | aef:hil | 233 | bce:hil | 273 | bde:hil | 313 | bef:hil | 353 | cdf:hil | 393 | def:hil |
| 34 | abd:hjk | 74 | abf:hjk | 114 | ace:hjk | 154 | ade:hjk | 194 | aef:hjk | 234 | bce:hjk | 274 | bde:hjk | 314 | bef:hjk | 354 | cdf:hjk | 394 | def:hjk |
| 35 | abd:hjl | 75 | abf:hjl | 115 | ace:hjl | 155 | ade:hjl | 195 | aef:hjl | 235 | bce:hjl | 275 | bde:hjl | 315 | bef:hjl | 355 | cdf:hjl | 395 | def:hjl |
| 36 | abd:hkl | 76 | abf:hkl | 116 | ace:hkl | 156 | ade:hkl | 196 | aef:hkl | 236 | bce:hkl | 276 | bde:hkl | 316 | bef:hkl | 356 | cdf:hkl | 396 | def:hkl |
| 37 | abd:ijk | 77 | abf:ijk | 117 | ace:ijk | 157 | ade:ijk | 197 | aef:ijk | 237 | bce:ijk | 277 | bde:ijk | 317 | bef:ijk | 357 | cdf:ijk | 397 | def:ijk |
| 38 | abd:ijl | 78 | abf:ijl | 118 | ace:ijl | 158 | ade:ijl | 198 | aef:ijl | 238 | bce:ijl | 278 | bde:ijl | 318 | bef:ijl | 358 | cdf:ijl | 398 | def:ijl |
| 39 | abd:ikl | 79 | abf:ikl | 119 | ace:ikl | 159 | ade:ikl | 199 | aef:ikl | 239 | bce:ikl | 279 | bde:ikl | 319 | bef:ikl | 359 | cdf:ikl | 399 | def:ikl |
| 40 | abd:jkl | 80 | abf:jkl | 120 | ace:jkl | 160 | ade:jkl | 200 | aef:jkl | 240 | bce:jkl | 280 | bde:jkl | 320 | bef:jkl | 360 | cdf:jkl | 400 | def:jkl |

- We are denoting the 12 possible alleles present in a hexaploid full-sib population with the letters $a$ to $f$ from 'Beauregard' parent and $g$ to $l$ from 'Tanzania'.
- The genotypes are denoted by their multi-allelic composition, were the first three letters were donated by 'Beauregard' and the second three letters were donated by 'Tanzania'.

**Supplementary Table 3.** Collinearity blocks aligned to *I. trifida* genome

| LG | Collinearity blocks | Segment length (cM) | (Mb) | Number of SNPs | Flanking Markers | | Orientation[1] |
|---|---|---|---|---|---|---|---|
| 1 | 1 | 79.00 | 7.52 | 531 | Tf_Tl_S1_29275_S1_1326265 | Tf_Tl_S1_7483434_S1_12668753 | forward |
| | 2 | 5.60 | 1.16 | 42 | Tf_Tl_S1_11926609_S1_21330091 | Tf_Tl_S1_13083712_S1_21220729 | forward |
| | 3 | 4.80 | 0.51 | 23 | Tf_Tl_S1_16223287_S1_24579737 | Tf_Tl_S1_16734334_S1_10632436 | forward |
| | 4 | 16.40 | 1.56 | 75 | Tf_Tl_S1_18331375_S1_10236812 | Tf_Tl_S1_19887833_S1_8179071 | forward |
| | 5 | 6.20 | 0.83 | 56 | Tf_Tl_S1_17233341_S1_7933361 | Tf_Tl_S1_18002738_S1_7503293 | forward |
| | 6 | 164.80 | 12.16 | 1233 | Tf_Tl_S1_20049620_S1_6890479 | Tf_S1_32150791 | forward |
| 2 | 1 | 132.20 | 9.46 | 1009 | Tf_Tl_S2_6022375_S2_166315 | Tf_Tl_S2_15486399_S2_8839719 | forward |
| | 2 | 6.40 | 1.31 | 46 | Tf_Tl_S2_3909445_S2_13303480 | Tf_S2_2685149 | backward |
| | 3 | 10.40 | 6.26 | 86 | Tf_Tl_S2_16666147_S2_14554551 | Tf_S2_22926064 | forward |
| 3 | 1 | 70.70 | 6.03 | 644 | Tf_S3_32030 | Tf_Tl_S3_6054979_S3_6854332 | forward |
| | 2 | 3.20 | 2.04 | 36 | Tf_Tl_S3_9849090_S3_12908964 | Tf_Tl_S3_7808516_S3_25024630 | backward |
| | 3 | 36.60 | 6.35 | 295 | Tf_Tl_S3_24593896_S3_26579108 | Tf_Tl_S3_18246508_S3_29056878 | backward |
| | 4 | 96.10 | 7.15 | 730 | Tf_Tl_S3_18200242_S3_29004840 | Tf_S3_11120432 | backward |
| 4 | 1 | 89.20 | 6.03 | 731 | Tf_S4_42497 | Tf_Tl_S4_6058108_S4_9418238 | forward |
| | 2 | 10.60 | 2.00 | 126 | Tf_Tl_S4_8789534_S4_10057931 | Tf_S4_6870014 | backward |
| | 3 | 4.80 | 1.17 | 45 | Tf_Tl_S4_9242703_S4_12336166 | Tf_Tl_S4_10352960_S4_13490533 | forward |
| | 4 | 120.50 | 21.85 | 1477 | Tf_Tl_S4_10615883_S4_13753157 | Tf_Tl_S4_32455528_S4_36257541 | forward |
| 5 | 1 | 157.10 | 25.06 | 1410 | Tf_Tl_S5_635908_S5_145116 | Tf_Tl_S5_25663026_S5_31247161 | forward |
| 6 | 1 | 30.40 | 10.55 | 185 | Tf_Tl_S6_11507438_S6_9365714 | Tf_Tl_S6_1169387_S6_11469757 | backward |
| | 2 | 158.90 | 13.39 | 1247 | Tf_S6_12595629 | Tf_S6_25945586 | forward |
| 7 | 1 | 79.20 | 6.18 | 655 | Tf_Tl_S7_104717_S7_153160 | Tf_Tl_S7_6229347_S7_6855558 | forward |
| | 2 | 21.80 | 7.44 | 364 | Tf_Tl_S7_19414899_S7_26111000 | Tf_Tl_S7_11978706_S7_13878608 | backward |
| | 3 | 9.30 | 4.36 | 116 | Tf_S7_6368703 | Tf_Tl_S7_10717988_S7_15769888 | forward |
| | 4 | 13.60 | 1.05 | 49 | Tf_Tl_S7_10805060_S7_17225271 | Tf_Tl_S7_11846988_S7_18660965 | forward |
| | 5 | 22.50 | 2.32 | 216 | Tf_Tl_S7_21529244_S7_18834983 | Tf_Tl_S7_23814851_S7_28838000 | forward |
| 8 | 1 | 95.50 | 9.73 | 768 | Tf_Tl_S8_19730_S8_67559 | Tf_Tl_S8_9744774_S8_12826247 | forward |
| | 2 | 4.10 | 1.02 | 27 | Tf_Tl_S8_10672128_S8_16731883 | Tf_Tl_S8_11687200_S8_17548648 | forward |
| | 3 | 5.20 | 0.77 | 24 | Tf_Tl_S8_14940643_S8_18606697 | Tf_Tl_S8_15600424_S8_18084540 | forward |
| 9 | 1 | 178.10 | 23.23 | 1939 | Tf_Tl_S9_32662_S9_165802 | Tf_S9_23192872 | forward |
| 10 | 1 | 188.70 | 24.38 | 1687 | Tf_Tl_S10_323131_S10_19201 | Tf_Tl_S10_24703543_S10_29035414 | forward |
| 11 | 1 | 145.60 | 18.90 | 1228 | Tf_Tl_S11_78090_S11_27388 | Tf_S11_18869116 | forward |
| 12 | 1 | 181.00 | 24.01 | 2036 | Tf_Tl_S12_84747_S12_104335 | Tf_Tl_S12_24045595_S12_28240613 | forward |
| 13 | 1 | 51.70 | 4.47 | 315 | Tf_Tl_S13_92583_S13_164808 | Tf_S13_4557388 | forward |
| | 2 | 8.70 | 5.48 | 121 | Tf_Tl_S13_12123894_S13_10529102 | Tf_S13_6645044 | backward |
| | 3 | 32.30 | 4.15 | 267 | Tf_S13_12233043 | Tf_Tl_S13_16380554_S13_24738853 | forward |
| | 4 | 10.40 | 0.77 | 71 | Tf_Tl_S13_17125139_S13_24888039 | Tf_Tl_S13_17900102_S13_25837095 | forward |
| | 5 | 71.30 | 4.82 | 481 | Tf_Tl_S13_18009588_S13_25969701 | Tf_Tl_S13_22831871_S13_31942957 | forward |
| 14 | 1 | 125.30 | 18.89 | 994 | Tf_Tl_S14_54061_S14_65490 | Tf_Tl_S14_18935024_S14_23563377 | forward |
| 15 | 1 | 166.60 | 22.10 | 1564 | Tf_Tl_S15_21822_S15_56000 | Tf_Tl_S15_22089841_S15_27054346 | forward |

1 Orientation of the segment when aligned to *I. trifida* genome.

**Supplementary Table 4.** Collinearity blocks aligned to *I. triloba* genome

| LG | Collinearity blocks | Segment length (cM) | (Mb) | Number of SNPs | Flanking Markers | | Orientation[1] |
|---|---|---|---|---|---|---|---|
| 1 | 1 | 7.10 | 1.30 | 84 | Tf_Tl_S1_57849_S1_22992 | Tf_Tl_S1_1131985_S1_1310872 | forward |
| | 2 | 54.10 | 4.23 | 325 | Tf_Tl_S1_1158286_S1_1338982 | Tf_Tl_S1_4826898_S1_5570702 | forward |
| | 3 | 7.50 | 1.56 | 61 | Tf_Tl_S1_6057894_S1_11267122 | Tf_Tl_S1_7483434_S1_12668753 | forward |
| | 4 | 6.10 | 1.03 | 46 | Tf_Tl_S1_21098794_S1_25934798 | Tf_Tl_S1_22334649_S1_26960935 | forward |
| | 5 | 6.50 | 0.77 | 54 | Tf_Tl_S1_22415841_S1_27162650 | Tf_Tl_S1_23098337_S1_27933185 | forward |
| | 6 | 31.60 | 3.73 | 323 | Tl_S1_27933241 | Tf_Tl_S1_26334481_S1_31660481 | forward |
| | 7 | 15.70 | 0.51 | 66 | Tf_Tl_S1_26431233_S1_31749608 | Tf_Tl_S1_26884800_S1_32262918 | forward |
| | 8 | 7.90 | 0.51 | 60 | Tl_S1_32297632 | Tf_Tl_S1_27341351_S1_32804566 | forward |
| | 9 | 42.80 | 3.04 | 304 | Tf_Tl_S1_27466912_S1_32965748 | Tf_Tl_S1_30316621_S1_35999501 | forward |
| | 10 | 13.40 | 0.35 | 59 | Tf_Tl_S1_30321098_S1_36003953 | Tl_S1_36331230 | forward |
| | 11 | 24.20 | 1.68 | 180 | Tf_Tl_S1_30665086_S1_36376446 | Tl_S1_38021290 | forward |
| 2 | 1 | 11.80 | 0.51 | 84 | Tf_Tl_S2_6022375_S2_166315 | Tf_Tl_S2_6471852_S2_625339 | forward |
| | 2 | 13.30 | 0.64 | 59 | Tl_S2_685685 | Tf_Tl_S2_7146647_S2_1324883 | forward |
| | 3 | 43.70 | 2.58 | 285 | Tf_Tl_S2_7149431_S2_1327959 | Tl_S2_3906489 | forward |
| | 4 | 34.40 | 2.98 | 283 | Tf_Tl_S2_9374419_S2_3909310 | Tf_Tl_S2_12060002_S2_6813139 | forward |
| | 5 | 10.70 | 1.08 | 87 | Tf_Tl_S2_12192826_S2_6922517 | Tf_Tl_S2_13020358_S2_7994159 | forward |
| | 6 | 3.70 | 0.72 | 34 | Tf_Tl_S2_14608512_S2_9753005 | Tf_Tl_S2_15267484_S2_9074844 | backward |
| | 7 | 4.10 | 1.08 | 30 | Tf_Tl_S2_3909445_S2_13303480 | Tl_S2_14364680 | forward |
| 3 | 1 | 72.20 | 7.09 | 636 | Tf_Tl_S3_32354_S3_22131 | Tl_S3_7103396 | forward |
| | 2 | 2.30 | 0.40 | 25 | Tl_S3_7265793 | Tf_Tl_S3_27590515_S3_7665371 | forward |
| | 3 | 2.80 | 1.26 | 36 | Tl_S3_7733994 | Tf_Tl_S3_26315975_S3_8993355 | forward |
| | 4 | 1.70 | 0.80 | 24 | Tf_Tl_S3_22384802_S3_23612789 | Tf_Tl_S3_21579582_S3_22808509 | backward |
| | 5 | 4.30 | 0.67 | 44 | Tf_Tl_S3_18841018_S3_29813927 | Tl_S3_29144198 | backward |
| | 6 | 3.80 | 0.39 | 21 | Tl_S3_21350185 | Tl_S3_20963574 | backward |
| | 7 | 1.90 | 0.17 | 31 | Tl_S3_20686766 | Tf_Tl_S3_17359096_S3_20516004 | backward |
| | 8 | 1.80 | 0.15 | 25 | Tf_Tl_S3_14821800_S3_17645995 | Tf_Tl_S3_14688858_S3_17497096 | backward |
| | 9 | 5.80 | 0.56 | 55 | Tf_Tl_S3_14340050_S3_17105234 | Tf_Tl_S3_13833939_S3_16547817 | backward |
| | 10 | 3.60 | 0.21 | 21 | Tl_S3_15150115 | Tf_Tl_S3_12407618_S3_14938849 | backward |
| | 11 | 4.10 | 0.47 | 28 | Tl_S3_14536763 | Tf_Tl_S3_11590962_S3_14068048 | backward |
| | 12 | 4.40 | 0.27 | 38 | Tf_Tl_S3_11588703_S3_14065783 | Tf_Tl_S3_11367494_S3_13828552 | backward |
| 4 | 1 | 77.70 | 5.62 | 551 | Tf_Tl_S4_104183_S4_27376 | Tf_Tl_S4_4865255_S4_5645598 | forward |
| | 2 | 18.40 | 2.26 | 194 | Tf_Tl_S4_4996736_S4_8349128 | Tf_Tl_S4_8282899_S4_10607605 | forward |
| | 3 | 13.70 | 4.45 | 160 | Tf_Tl_S4_8084173_S4_10775296 | Tl_S4_15226024 | forward |
| | 4 | 4.10 | 1.96 | 53 | Tf_Tl_S4_11823581_S4_15277834 | Tf_Tl_S4_13271494_S4_17236636 | forward |
| | 5 | 8.80 | 2.04 | 96 | Tl_S4_25122962 | Tl_S4_27164284 | forward |
| | 6 | 13.60 | 1.36 | 167 | Tf_Tl_S4_24912577_S4_27727566 | Tf_Tl_S4_26153038_S4_29086938 | forward |
| | 7 | 17.00 | 1.68 | 264 | Tf_Tl_S4_26325738_S4_29431191 | Tf_Tl_S4_27802680_S4_31113965 | forward |
| | 8 | 9.10 | 0.64 | 102 | Tl_S4_31126630 | Tf_Tl_S4_28385921_S4_31770263 | forward |
| | 9 | 50.80 | 4.46 | 564 | Tl_S4_31802612 | Tf_Tl_S4_32455528_S4_36257541 | forward |
| 5 | 1 | 41.80 | 6.97 | 292 | Tf_Tl_S5_635908_S5_145116 | Tf_Tl_S5_7038596_S5_7021460 | forward |
| | 2 | 2.50 | 1.80 | 50 | Tf_Tl_S5_12203156_S5_15662930 | Tf_Tl_S5_13198341_S5_17388037 | forward |
| | 3 | 2.30 | 1.19 | 33 | Tf_Tl_S5_13725685_S5_18012762 | Tf_Tl_S5_14893736_S5_19202238 | forward |
| | 4 | 28.20 | 4.87 | 244 | Tl_S5_19906010 | Tf_Tl_S5_19824179_S5_24771028 | forward |
| | 5 | 14.40 | 1.64 | 154 | Tf_Tl_S5_20273199_S5_25262493 | Tf_Tl_S5_21773576_S5_26904448 | forward |
| | 6 | 43.30 | 3.38 | 377 | Tf_Tl_S5_21831005_S5_26966096 | Tf_Tl_S5_24838095_S5_30339693 | forward |
| | 7 | 12.30 | 0.91 | 127 | Tf_Tl_S5_24880523_S5_30378175 | Tf_Tl_S5_25663026_S5_31247161 | forward |
| 6 | 1 | 4.30 | 2.19 | 39 | Tf_Tl_S6_1411732_S6_11289037 | Tf_Tl_S6_14081350_S6_13453545 | forward |
| | 2 | 19.10 | 3.56 | 148 | Tf_Tl_S6_14313357_S6_13695787 | Tl_S6_17258142 | forward |
| | 3 | 64.90 | 5.29 | 525 | Tf_Tl_S6_17893289_S6_17490185 | Tf_Tl_S6_22581137_S6_22764969 | forward |
| | 4 | 24.80 | 1.33 | 208 | Tl_S6_22882911 | Tf_Tl_S6_23759979_S6_24151356 | forward |
| | 5 | 44.30 | 2.48 | 284 | Tf_Tl_S6_23852908_S6_24253366 | Tl_S6_26692439 | forward |
| 7 | 1 | 21.60 | 1.38 | 156 | Tf_Tl_S7_104717_S7_153160 | Tl_S7_1529523 | forward |
| | 2 | 8.80 | 0.88 | 95 | Tl_S7_1881945 | Tl_S7_2757562 | forward |
| | 3 | 17.00 | 1.12 | 125 | Tf_Tl_S7_2606845_S7_2767196 | Tf_Tl_S7_3683723_S7_3887501 | forward |
| | 4 | 27.10 | 2.87 | 214 | Tf_Tl_S7_3696434_S7_3899939 | Tl_S7_6774703 | forward |
| | 5 | 1.80 | 0.52 | 55 | Tf_Tl_S7_18548499_S7_25085801 | Tf_Tl_S7_18168273_S7_24599871 | backward |
| | 6 | 1.80 | 0.49 | 26 | Tf_Tl_S7_17692567_S7_24127425 | Tf_Tl_S7_17263661_S7_23633522 | backward |
| | 7 | 3.80 | 1.35 | 59 | Tf_Tl_S7_9455056_S7_14148674 | Tf_Tl_S7_10430722_S7_15492801 | forward |
| | 8 | 2.50 | 0.28 | 21 | Tf_Tl_S7_10438987_S7_15499672 | Tf_Tl_S7_10717988_S7_15769888 | forward |
| | 9 | 17.00 | 1.64 | 58 | Tf_Tl_S7_10804966_S7_17225177 | Tf_Tl_S7_21557539_S7_18862031 | forward |
| | 10 | 5.80 | 0.70 | 40 | Tf_Tl_S7_21729104_S7_26422262 | Tf_Tl_S7_22378155_S7_27119652 | forward |
| | 11 | 11.30 | 1.42 | 139 | Tf_Tl_S7_22617578_S7_27455930 | Tl_S7_28837994 | forward |
| 8 | 1 | 8.50 | 0.86 | 90 | Tf_Tl_S8_19730_S8_67559 | Tf_Tl_S8_740571_S8_880890 | forward |
| | 2 | 80.50 | 8.61 | 598 | Tf_Tl_S8_836713_S8_979128 | Tf_Tl_S8_7612856_S8_9588058 | forward |
| 9 | 1 | 6.80 | 0.49 | 91 | Tf_Tl_S9_32662_S9_165802 | Tf_Tl_S9_486602_S9_659522 | forward |
| | 2 | 18.10 | 1.35 | 181 | Tf_Tl_S9_530290_S9_872845 | Tf_Tl_S9_1748627_S9_2219923 | forward |
| | 3 | 16.00 | 0.60 | 119 | Tf_Tl_S9_1770275_S9_2242641 | Tf_Tl_S9_2278873_S9_2797499 | forward |
| | 4 | 72.70 | 8.19 | 893 | Tl_S9_2903732 | Tf_Tl_S9_9485147_S9_11068446 | forward |
| | 5 | 9.10 | 4.41 | 142 | Tf_Tl_S9_9918112_S9_11566782 | Tf_Tl_S9_13868549_S9_15420189 | forward |
| | 6 | 47.40 | 7.79 | 305 | Tf_Tl_S9_18264262_S9_23550142 | Tf_Tl_S9_23003510_S9_31339449 | forward |
| 10 | 1 | 2.80 | 0.28 | 26 | Tf_Tl_S10_323131_S10_19201 | Tf_Tl_S10_574798_S10_303708 | forward |
| | 2 | 64.20 | 9.52 | 465 | Tf_Tl_S10_589768_S10_326554 | Tf_Tl_S10_7822069_S10_9628784 | forward |
| | 3 | 48.60 | 7.69 | 462 | Tf_Tl_S10_12609267_S10_16669649 | Tf_Tl_S10_20065184_S10_24357774 | forward |
| | 4 | 4.40 | 0.38 | 38 | Tf_Tl_S10_20214674_S10_24545917 | Tf_Tl_S10_20582216_S10_24923575 | forward |
| | 5 | 60.20 | 3.87 | 531 | Tf_Tl_S10_21162980_S10_25162682 | Tf_Tl_S10_24703543_S10_29035414 | forward |
| 11 | 1 | 2.50 | 0.24 | 61 | Tf_Tl_S11_78090_S11_27388 | Tl_S11_265837 | forward |
| | 2 | 1.10 | 0.08 | 23 | Tl_S11_310588 | Tl_S11_365252 | forward |
| | 3 | 6.70 | 0.71 | 61 | Tf_Tl_S11_539878_S11_513123 | Tl_S11_1221341 | forward |
| | 4 | 21.40 | 1.34 | 152 | Tl_S11_1241649 | Tf_Tl_S11_2446677_S11_2582697 | forward |
| | 5 | 27.30 | 1.78 | 251 | Tf_Tl_S11_2487926_S11_2909146 | Tl_S11_4694110 | forward |
| | 6 | 14.30 | 1.40 | 113 | Tf_Tl_S11_4001503_S11_4796937 | Tf_Tl_S11_5032922_S11_6180380 | forward |
| | 7 | 26.50 | 4.92 | 218 | Tf_Tl_S11_5176716_S11_6554942 | Tl_S11_11476185 | forward |
| | 8 | 1.30 | 0.49 | 22 | Tf_Tl_S11_9770531_S11_12596143 | Tf_Tl_S11_10422188_S11_13090284 | forward |
| | 9 | 3.80 | 0.85 | 32 | Tf_Tl_S11_15630311_S11_21784824 | Tf_Tl_S11_16289221_S11_22637610 | forward |
| | 10 | 15.60 | 2.41 | 108 | Tf_Tl_S11_17181171_S11_23184162 | Tl_S11_25434031 | forward |
| 12 | 1 | 11.10 | 0.99 | 118 | Tf_Tl_S12_84747_S12_104335 | Tl_S12_1091578 | forward |
| | 2 | 9.00 | 0.47 | 74 | Tf_Tl_S12_991659_S12_1183028 | Tf_Tl_S12_1413505_S12_1649434 | forward |
| | 3 | 41.50 | 4.29 | 321 | Tf_Tl_S12_1438422_S12_1676770 | Tl_S12_5970050 | forward |
| | 4 | 23.50 | 5.28 | 269 | Tf_Tl_S12_4763751_S12_6087682 | Tf_Tl_S12_9241128_S12_11093043 | forward |
| | 5 | 2.70 | 1.10 | 36 | Tf_Tl_S12_9462557_S12_11452192 | Tf_Tl_S12_10358937_S12_12549455 | forward |
| | 6 | 18.50 | 3.73 | 254 | Tf_Tl_S12_15852681_S12_19326860 | Tf_Tl_S12_19532922_S12_23057769 | forward |
| | 7 | 63.60 | 4.78 | 677 | Tf_Tl_S12_19540865_S12_23065650 | Tf_Tl_S12_23659810_S12_27808405 | forward |
| | 8 | 6.00 | 0.39 | 80 | Tf_Tl_S12_23698267_S12_27850253 | Tf_Tl_S12_24045595_S12_28240613 | forward |
| 13 | 1 | 34.30 | 3.82 | 176 | Tf_Tl_S13_92583_S13_164808 | Tf_Tl_S13_2304078_S13_3988683 | forward |
| | 2 | 17.40 | 2.83 | 101 | Tf_Tl_S13_2335261_S13_5497598 | Tf_Tl_S13_4557896_S13_8323307 | forward |
| | 3 | 1.00 | 0.95 | 22 | Tf_Tl_S13_9919288_S13_8856116 | Tf_Tl_S13_9157972_S13_9801335 | forward |
| | 4 | 43.40 | 6.53 | 327 | Tl_S13_19352891 | Tl_S13_25881791 | forward |
| | 5 | 6.50 | 0.74 | 79 | Tf_Tl_S13_18216678_S13_26254636 | Tf_Tl_S13_18887009_S13_26958210 | forward |
| | 6 | 57.70 | 4.73 | 357 | Tf_Tl_S13_19105035_S13_27211345 | Tf_Tl_S13_22831871_S13_31942957 | forward |

Continued on next page

**Supplementary Table 4 – continued from previous page**

| LG | Collinearity blocks | Segment length (cM) | (Mb) | Number of SNPs | Flanking Markers | | Orientation[1] |
|---|---|---|---|---|---|---|---|
| 14 | 1 | 55.10 | 6.20 | 316 | Tf_Tl_S14_54061_S14_65490 | Tf_Tl_S14_5384499_S14_6266762 | forward |
| | 2 | 5.00 | 3.40 | 67 | Tf_Tl_S14_6170838_S14_13061670 | Tl_S14_16459800 | forward |
| | 3 | 16.50 | 1.70 | 105 | Tl_S14_18776018 | Tf_Tl_S14_16181551_S14_20474284 | forward |
| | 4 | 34.20 | 2.89 | 352 | Tf_Tl_S14_16380210_S14_20688413 | Tf_Tl_S14_18935024_S14_23563377 | forward |
| 15 | 1 | 51.60 | 3.41 | 446 | Tf_Tl_S15_21822_S15_56000 | Tf_Tl_S15_3053327_S15_3466955 | forward |
| | 2 | 17.20 | 1.64 | 174 | Tf_Tl_S15_3062973_S15_3477689 | Tf_Tl_S15_4566719_S15_5120949 | forward |
| | 3 | 45.00 | 10.09 | 515 | Tl_S15_5150780 | Tl_S15_15238297 | forward |
| | 4 | 50.00 | 7.14 | 372 | Tl_S15_19946745 | Tf_Tl_S15_22089841_S15_27054346 | forward |

1 Orientation of the segment when aligned to *I. triloba* genome.

**Supplementary Table 5.** Number of multivalent signatures in parents 'Beauregard' and 'Tanzania'

| Linkage group | Multivalent signatures* in 'Beauregard' | Multivalent signatures* in 'Tanzania' | Total | Map length |
|---|---|---|---|---|
| 1 | 77 | 101 | 178 | 290.4 |
| 2 | 25 | 38 | 63 | 184.6 |
| 3 | 63 | 20 | 83 | 222.1 |
| 4 | 74 | 89 | 163 | 227.1 |
| 5 | 35 | 23 | 58 | 156.8 |
| 6 | 22 | 43 | 65 | 189.3 |
| 7 | 51 | 43 | 94 | 155.1 |
| 8 | 11 | 3 | 14 | 115.5 |
| 9 | 38 | 74 | 112 | 178.0 |
| 10 | 69 | 58 | 127 | 188.7 |
| 11 | 18 | -** | -** | 145.6 |
| 12 | 39 | 55 | 94 | 180.0 |
| 13 | 41 | 42 | 83 | 179.2 |
| 14 | 25 | 26 | 51 | 125.4 |
| 15 | 22 | 26 | 48 | 164.2 |

* The number of multivalent signatures was compared between both parents using a paired t-test and there was no evidence of difference between 'Beauregard' and 'Tanzania' (P = 0.52).
** LG 11 for parent 'Tanzania' was mostly inconclusive and was not considered.
