## Supplementary File 1 for "Unraveling the hexaploid sweetpotato inheritance using ultra-dense multilocus mapping"

**Description:** This file contains a schematic overview of the map order improvement using *I. trifida* reference genome in all LGs (except 7, which is already in the Supplementary Note).

1. In linkage groups 1, 2, 3, 4, 6, 8, and 13, we detected collinearity blocks by visually inspecting abrupt breakages in the scatter plots continuity. The description of the four panels for each figure is the following:
A) Scatter plot of the physical distance in *I. trifida* reference genome (horizontal axis) versus the genetic distance in the *de novo* genetic map (colored dots) and the genome-assisted genetic map (dark blue dots). The *de novo* and the genome-assisted maps and their recombination fraction matrices are presented to the left and to the right-hand side of the scatter plot, respectively. Using visual inspection, we detected distinct segments (separated by vertical dashed lines) in the *de novo* scatter plot. We reconstructed sub-maps for each segment using both *de novo* and reference genome orders. The multilocus log-likelihood and the final length of each segment is presented in panel B. We then tested a set of complete map models concatenating all sub-maps segments (panel C). For all models, based on visual inspection, we proposed the segment order. In panel C, (ga) indicates genomic-assisted order within segments and (dn) indicates *de novo* order. The orientation of the segments, presented in panel C by the black arrows, were also obtained by visual inspection. Right arrows (🡪) indicate collinearity blocks with the same orientation of the *I. trifida* genome (forward); left arrows (🡨) indicate collinearity blocks with the opposite orientation of the I. trifida genome (backward); The model with highest log-likelihood is indicated in bold. Panel D shows a final comparison of the map obtained using the *de novo* and the genome-assisted map indicating, in all cases, a clear superiority of the latter. The final order of the sub-maps is also indicated in panel A by the read arrows connecting segments.
2. In linkage groups 5, 9, 10, 11, 12, 14, and 15, we did not detect abrupt breakages in the scatter plots continuity, we only compared the *de novo* order and the genome-assisted order. The description of the two panels for each figure is the following:

A) Scatter plot of the physical distance in *I. trifida* reference genome (horizontal axis) versus the genetic distance in the *de novo* genetic map (coloured dots) and the genome-assisted genetic map (dark blue dots). The *de novo* and the genome-assisted maps and their recombination fraction matrices are presented to the left and to the right-hand side of the scatter plot, respectively. Panel B shows a final comparison of the map obtained using the *de novo* and the genome-assisted map indicating, in all cases, a clear superiority of the latter.

**Linkage Group 1**

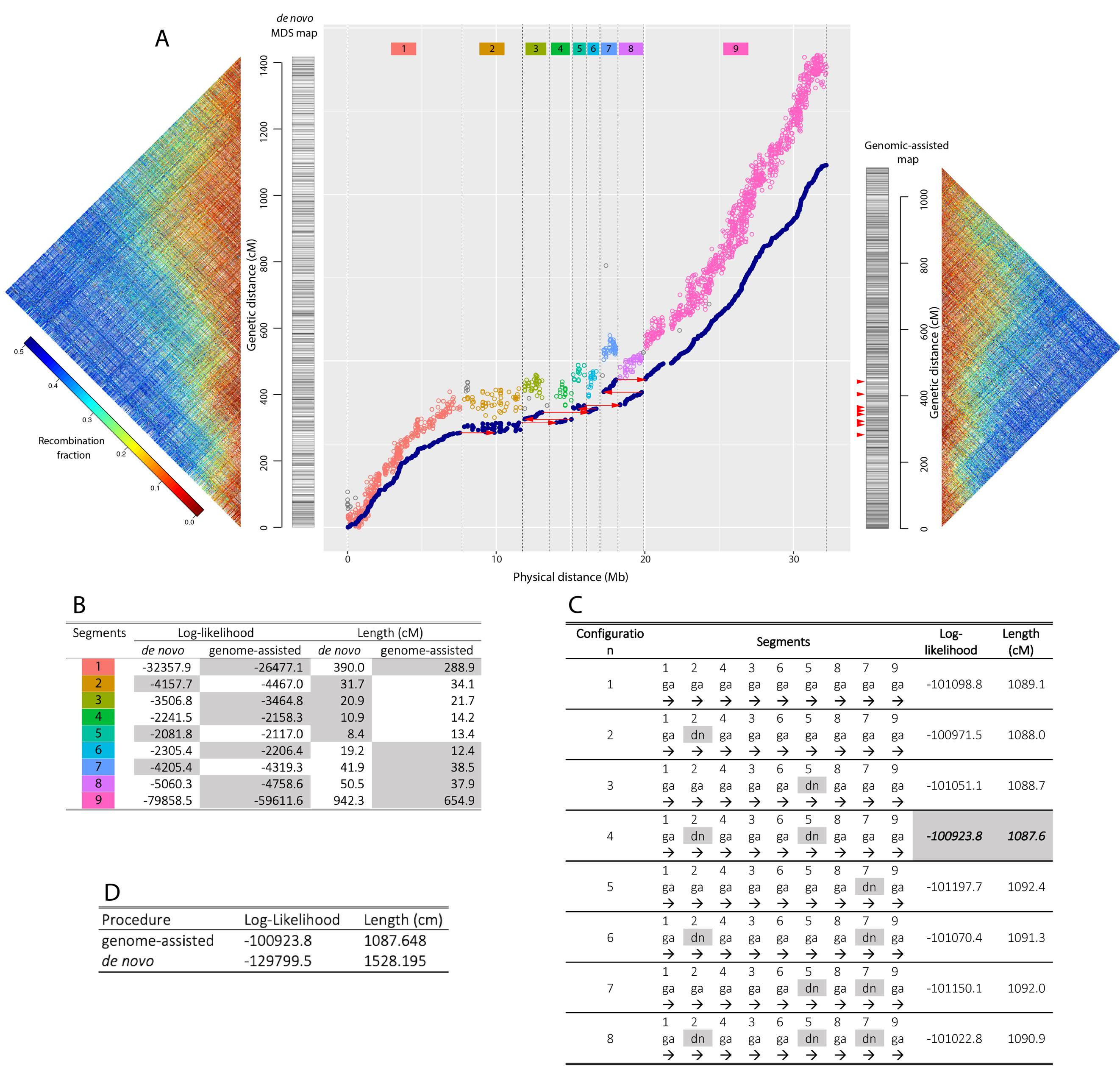

**Linkage Group 2^*^**

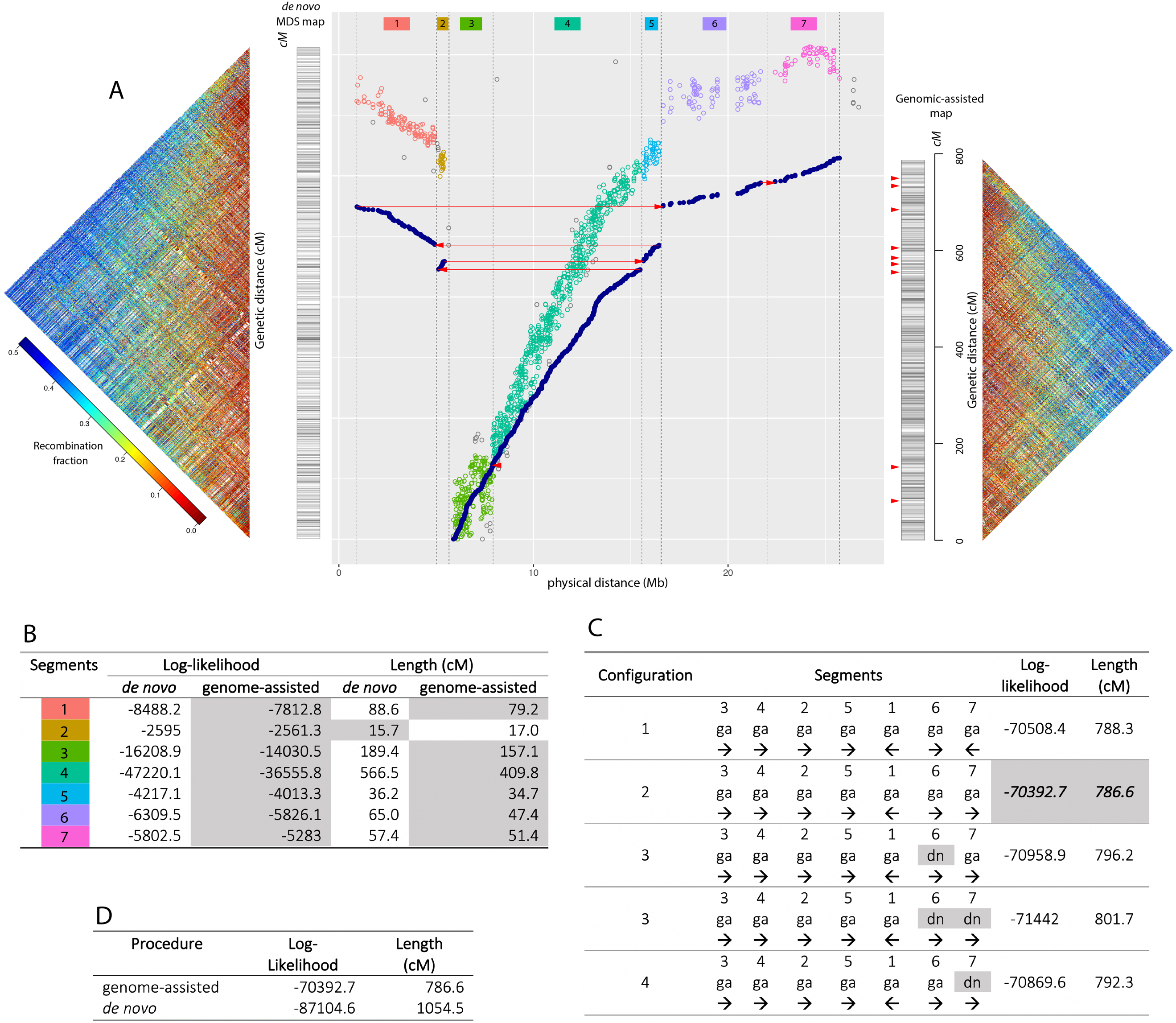

* In the final map, the marker order selected here produced substantial gaps between segments 4 and 5, and 6 and 7. Thus, we used the *de novo* order of segments 5 and 7 in the final map.

**Linkage Group 3**

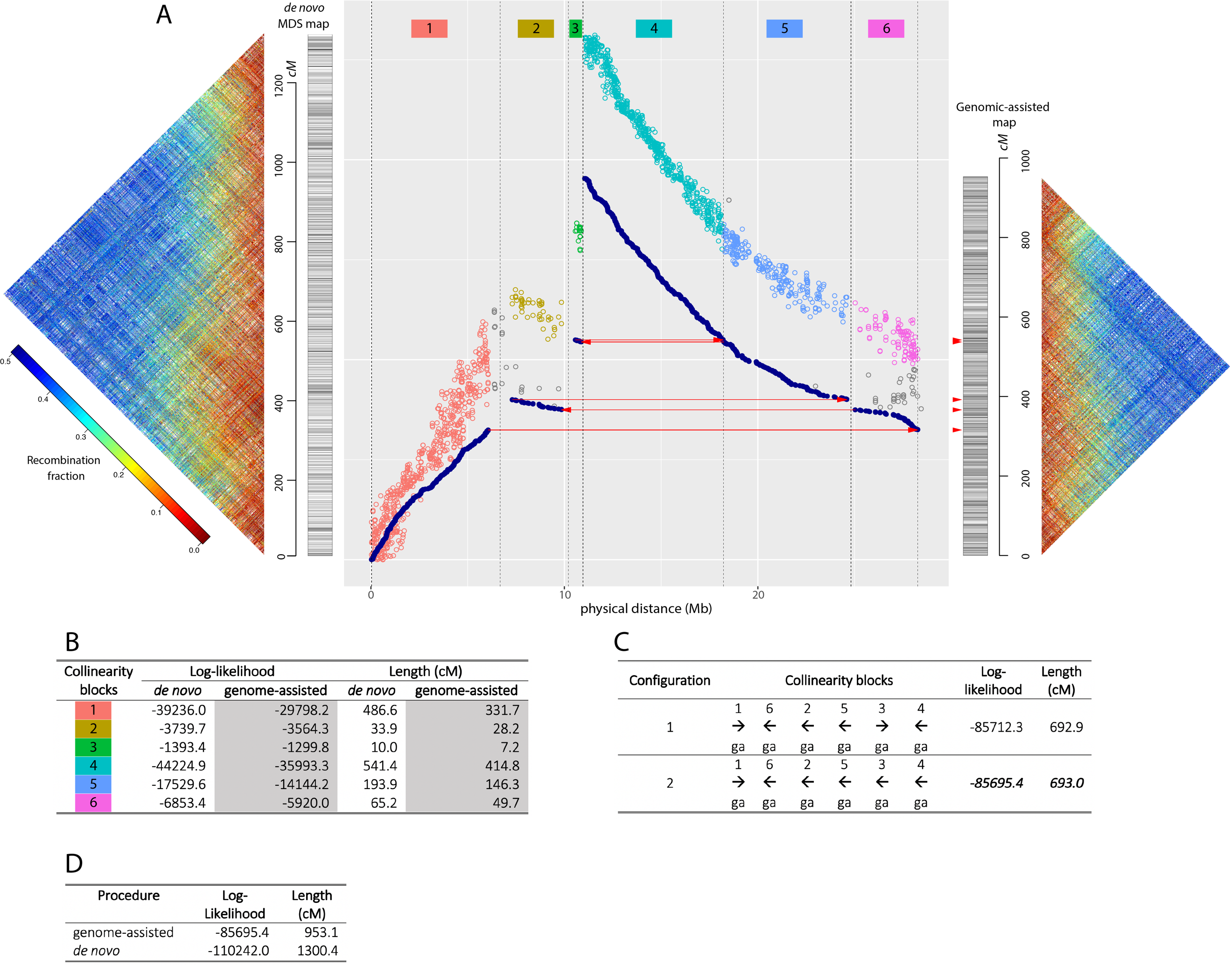

**Linkage Group 4**

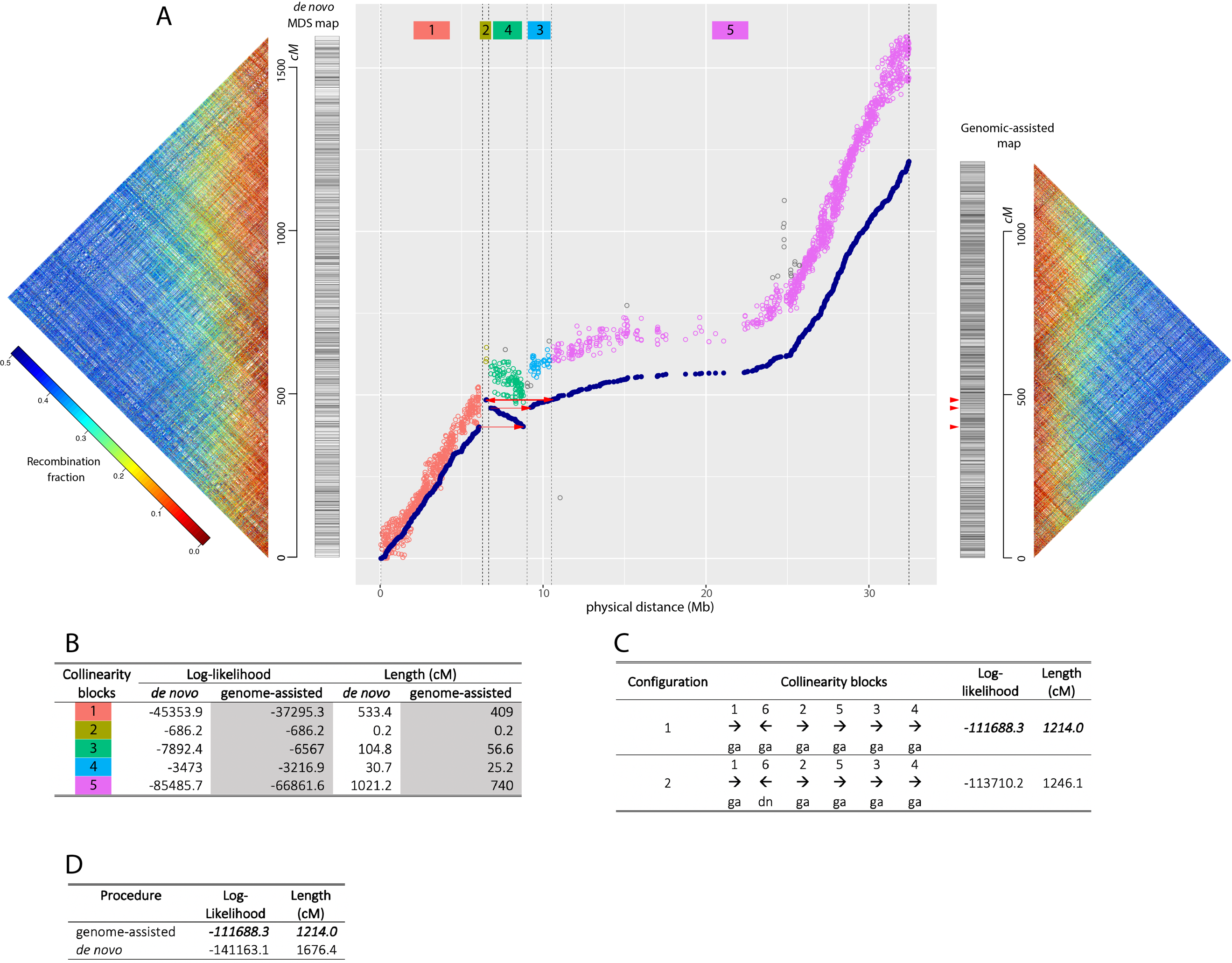

**Linkage Group 5**

A

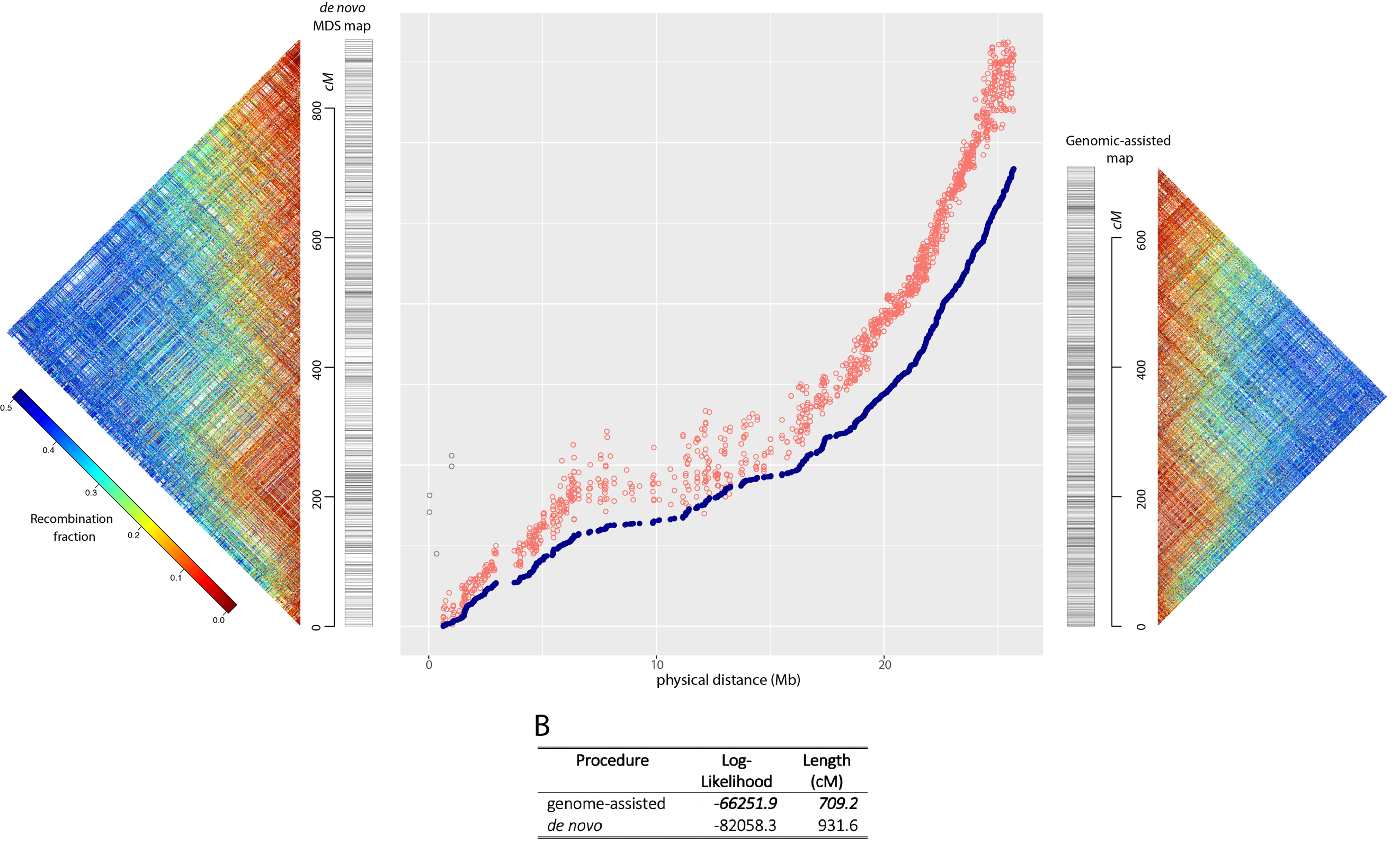

**Linkage Group 6**

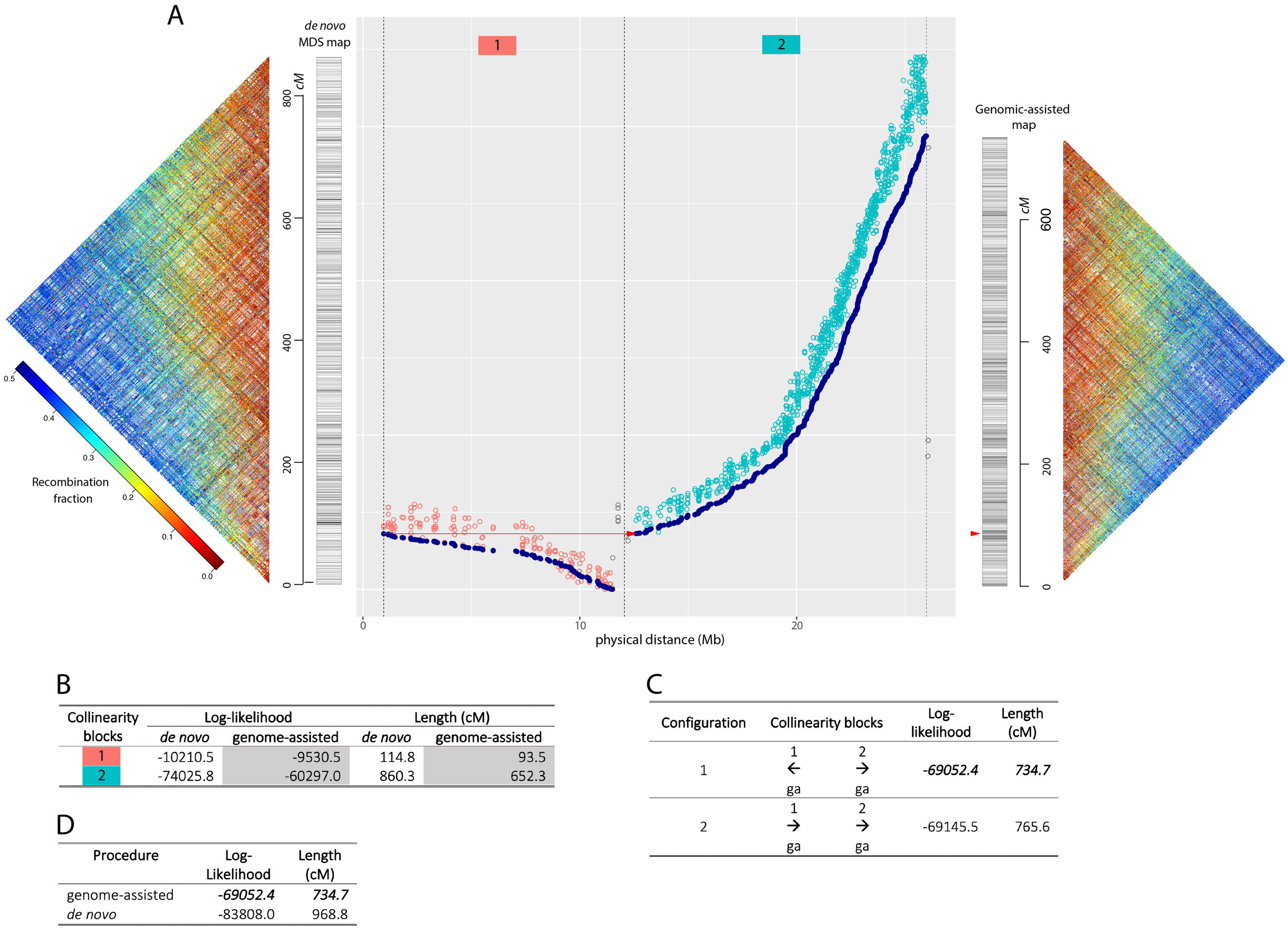

**Linkage Group 8**
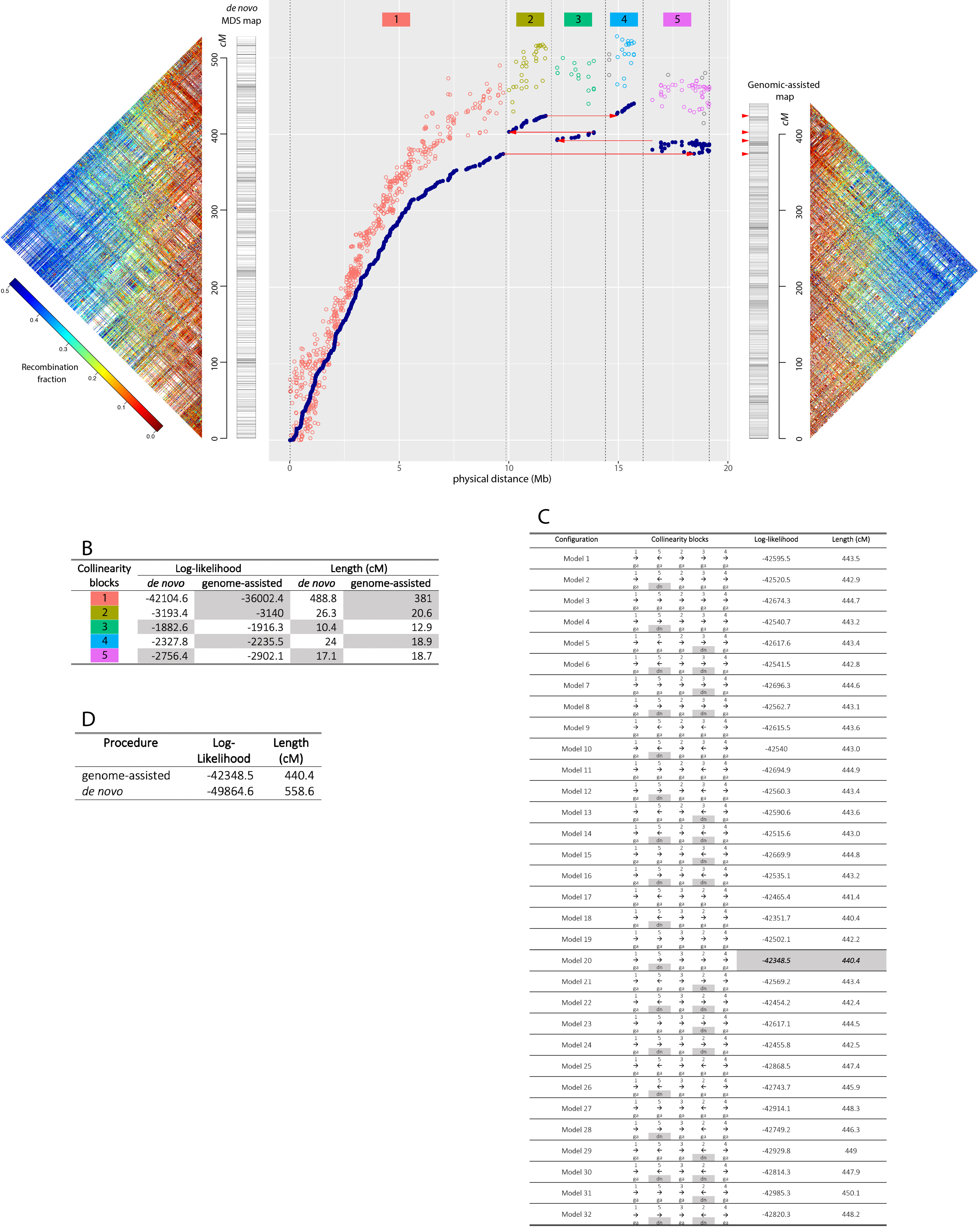

A

**Linkage Group 9**

A

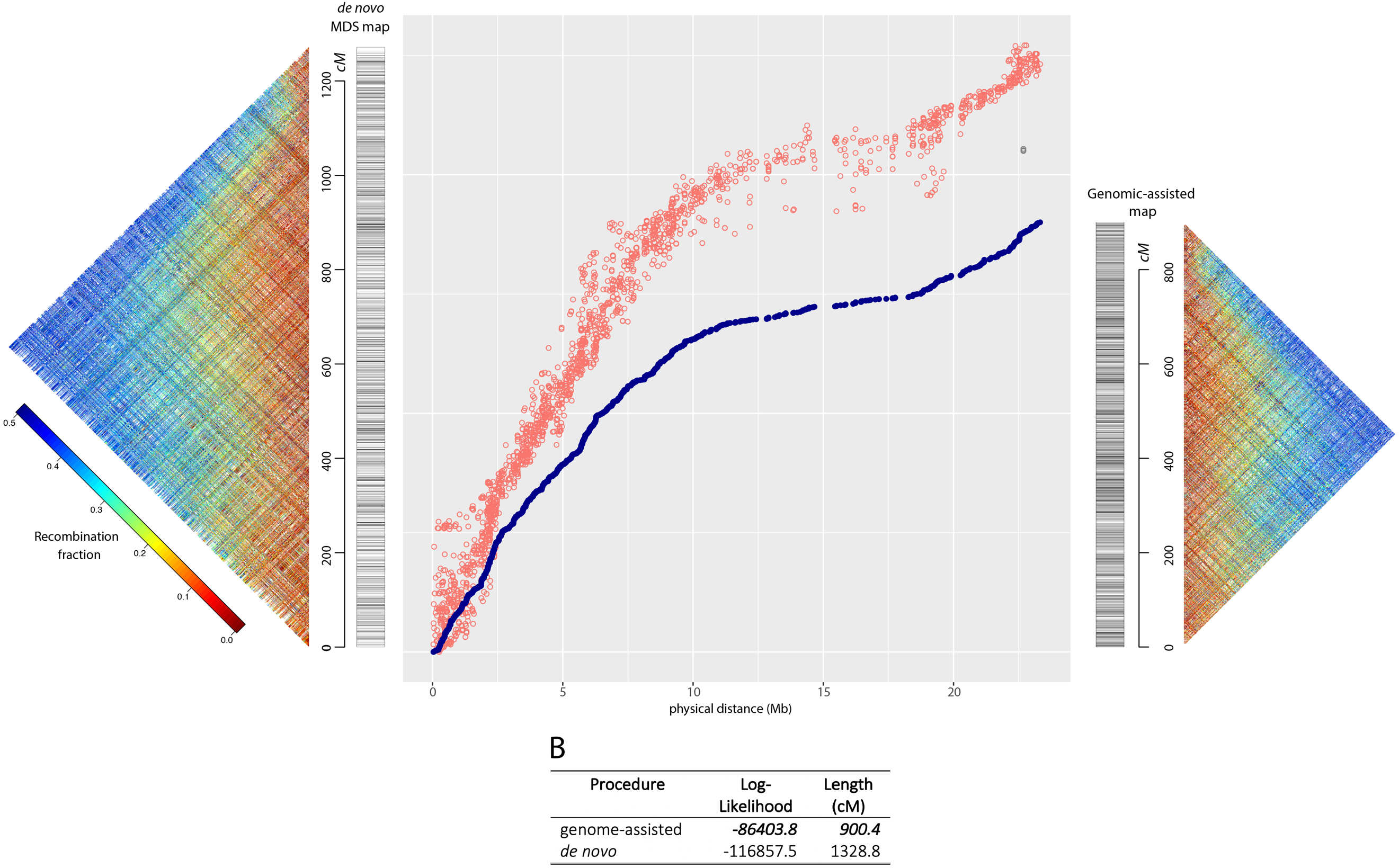

**Linkage Group 10**
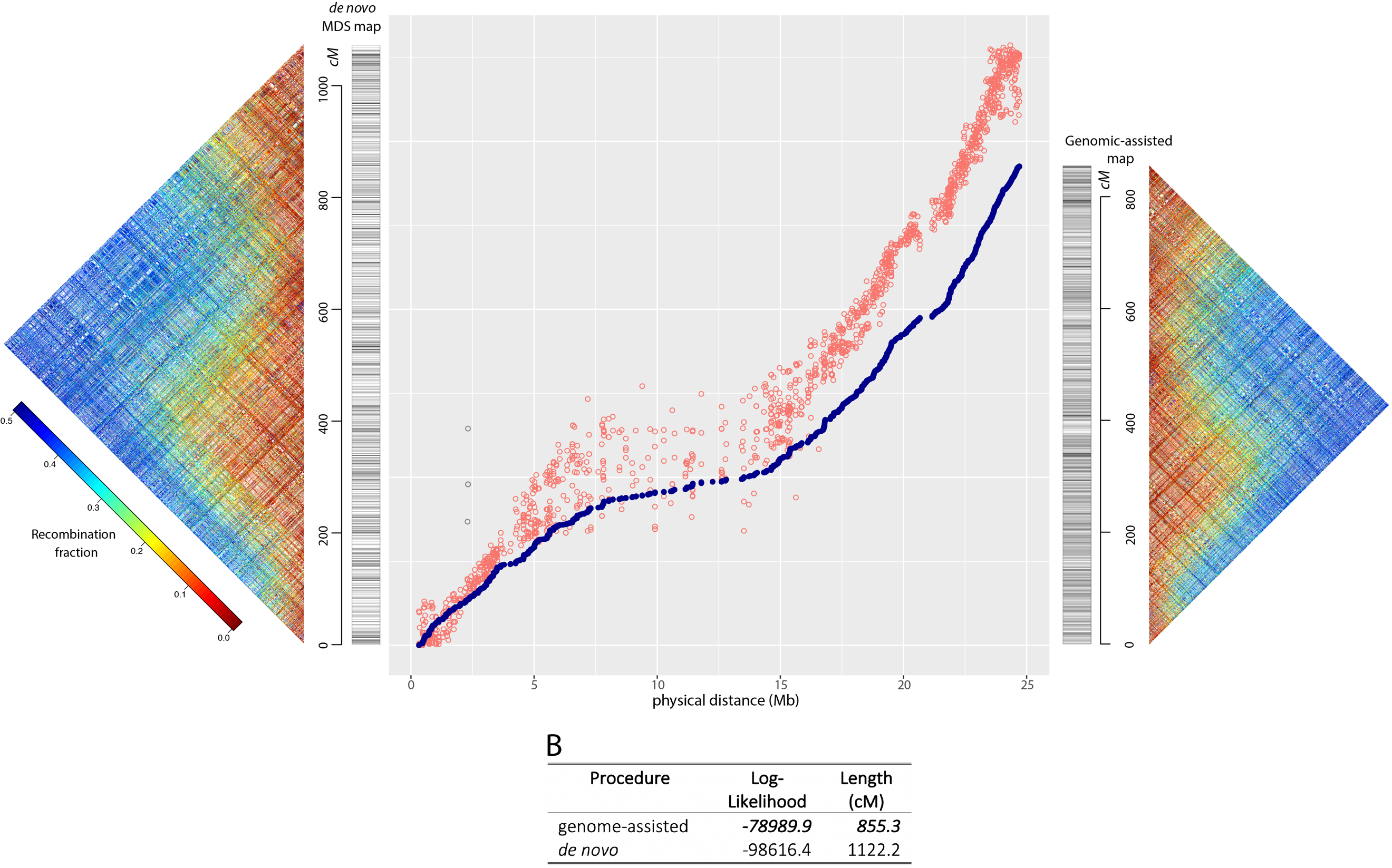

A

**Linkage Group 11**

A

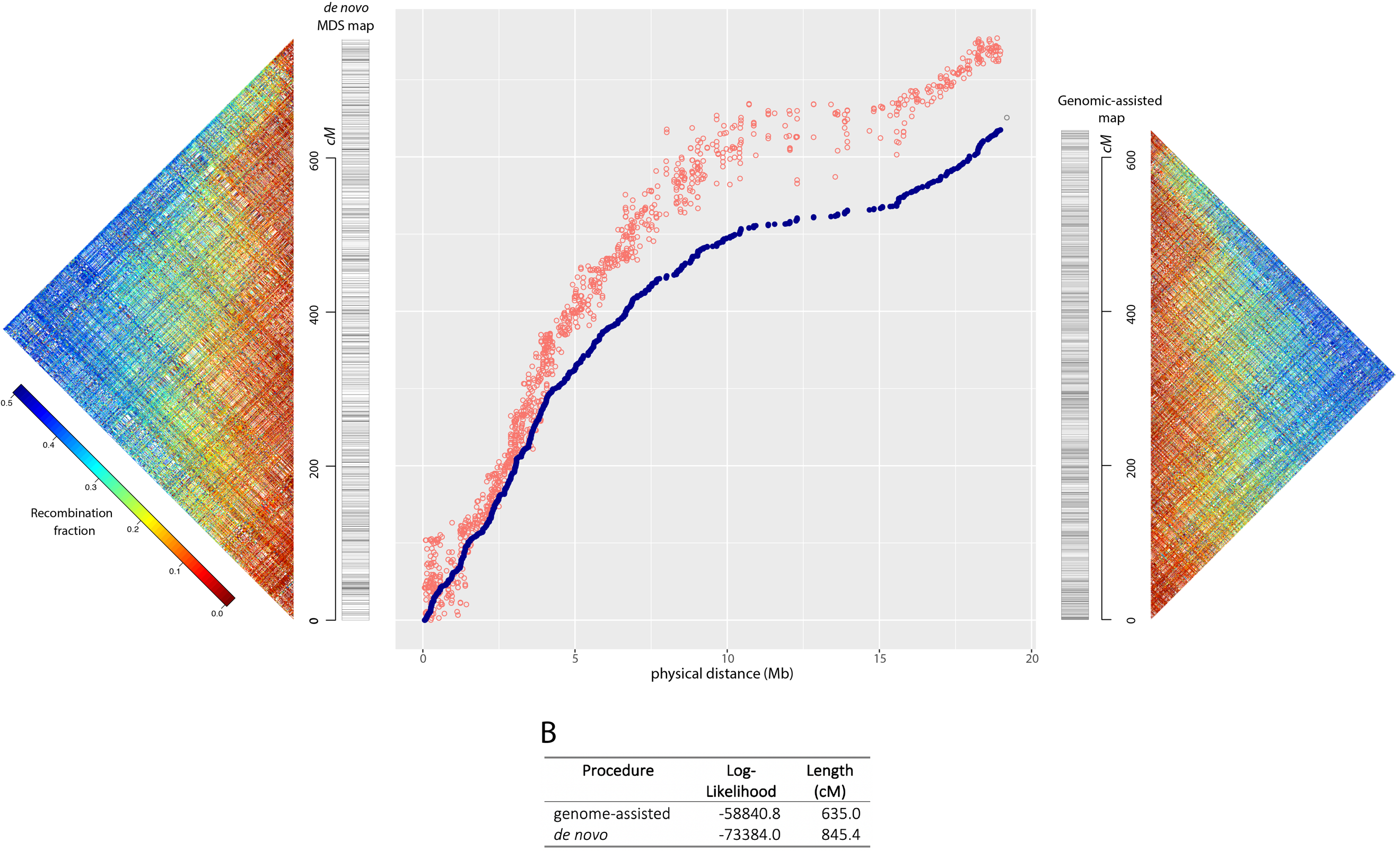

**Linkage Group 12**
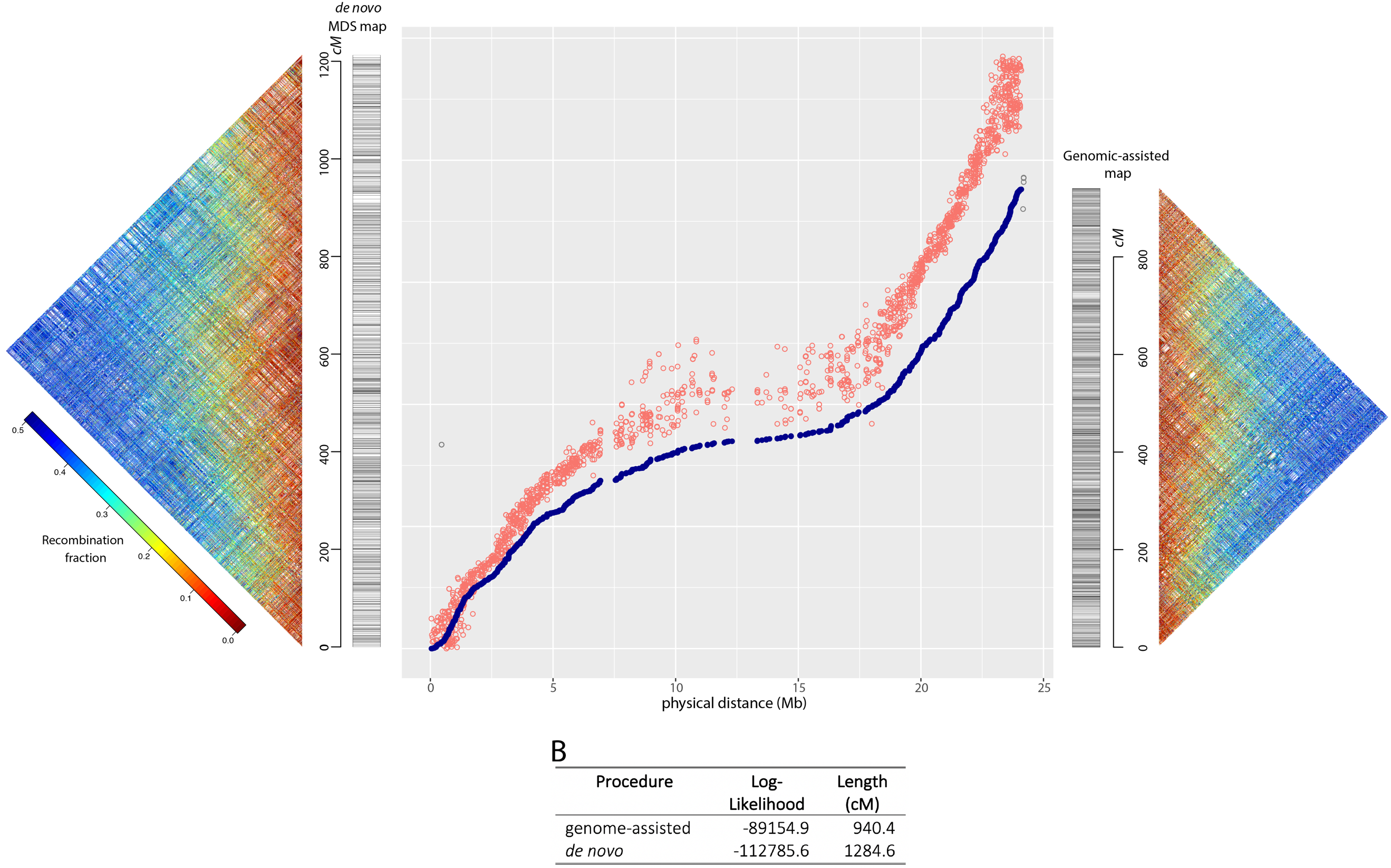

A

**Linkage Group 13**

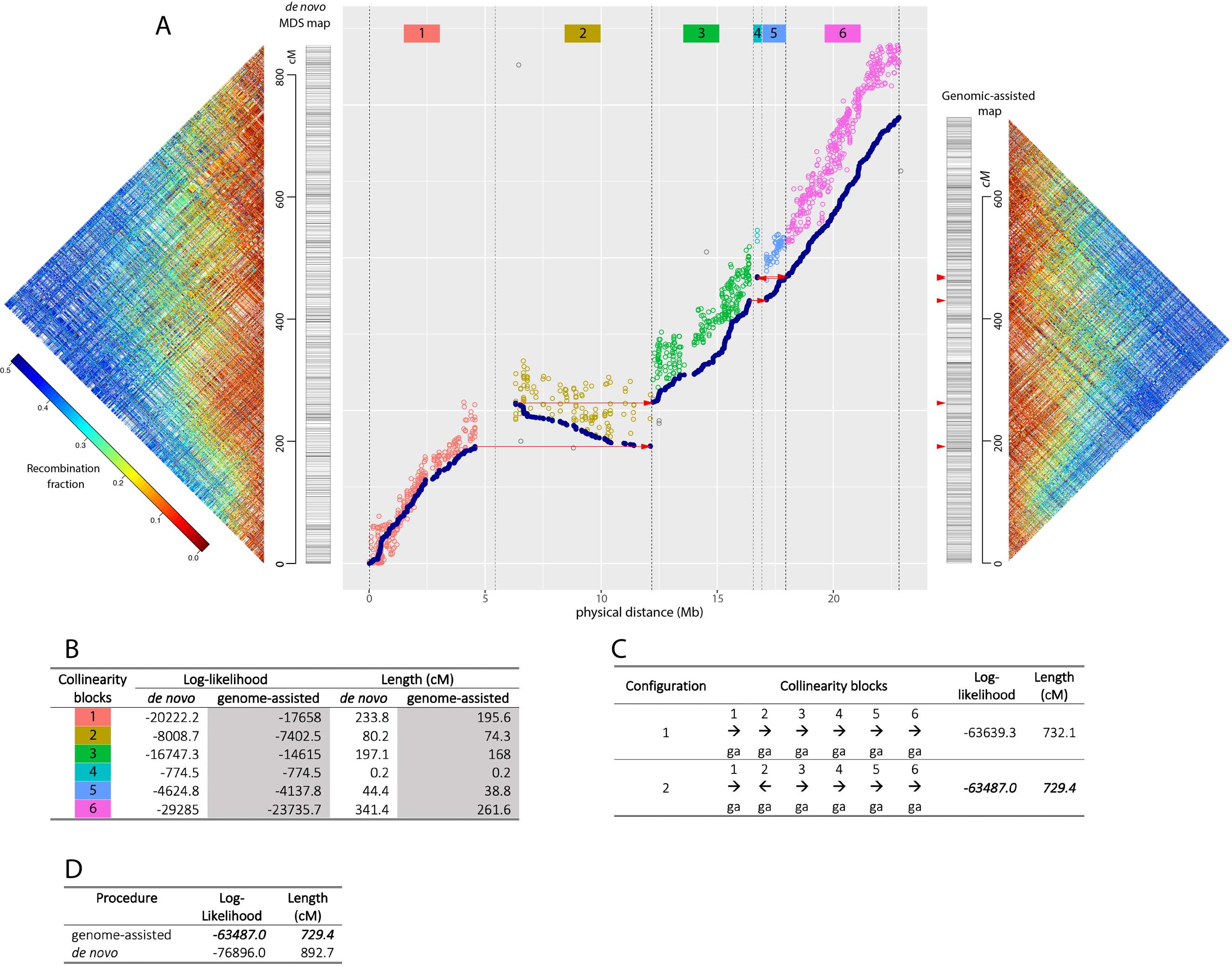

**Linkage Group 14**

A

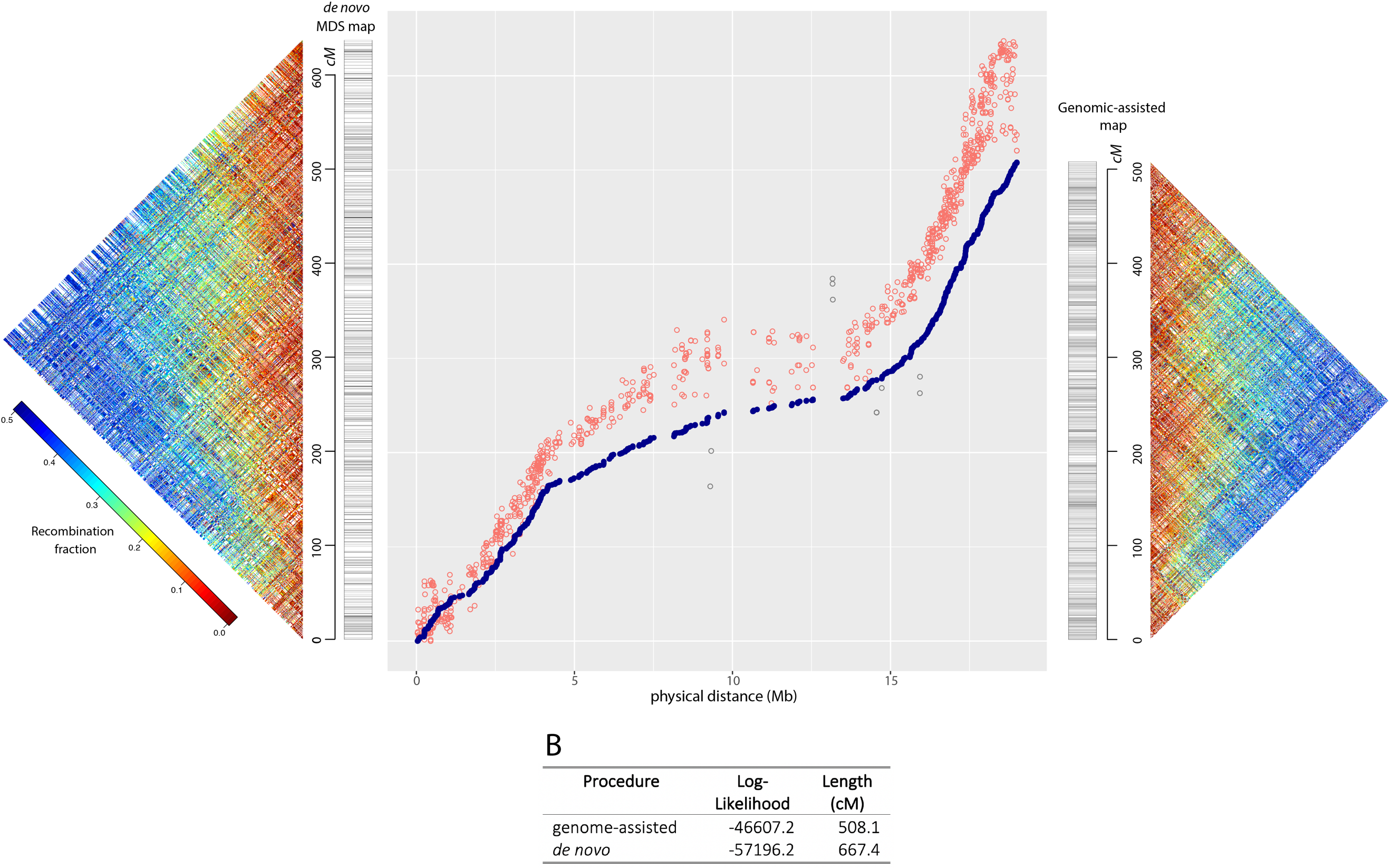

**Linkage Group 15**
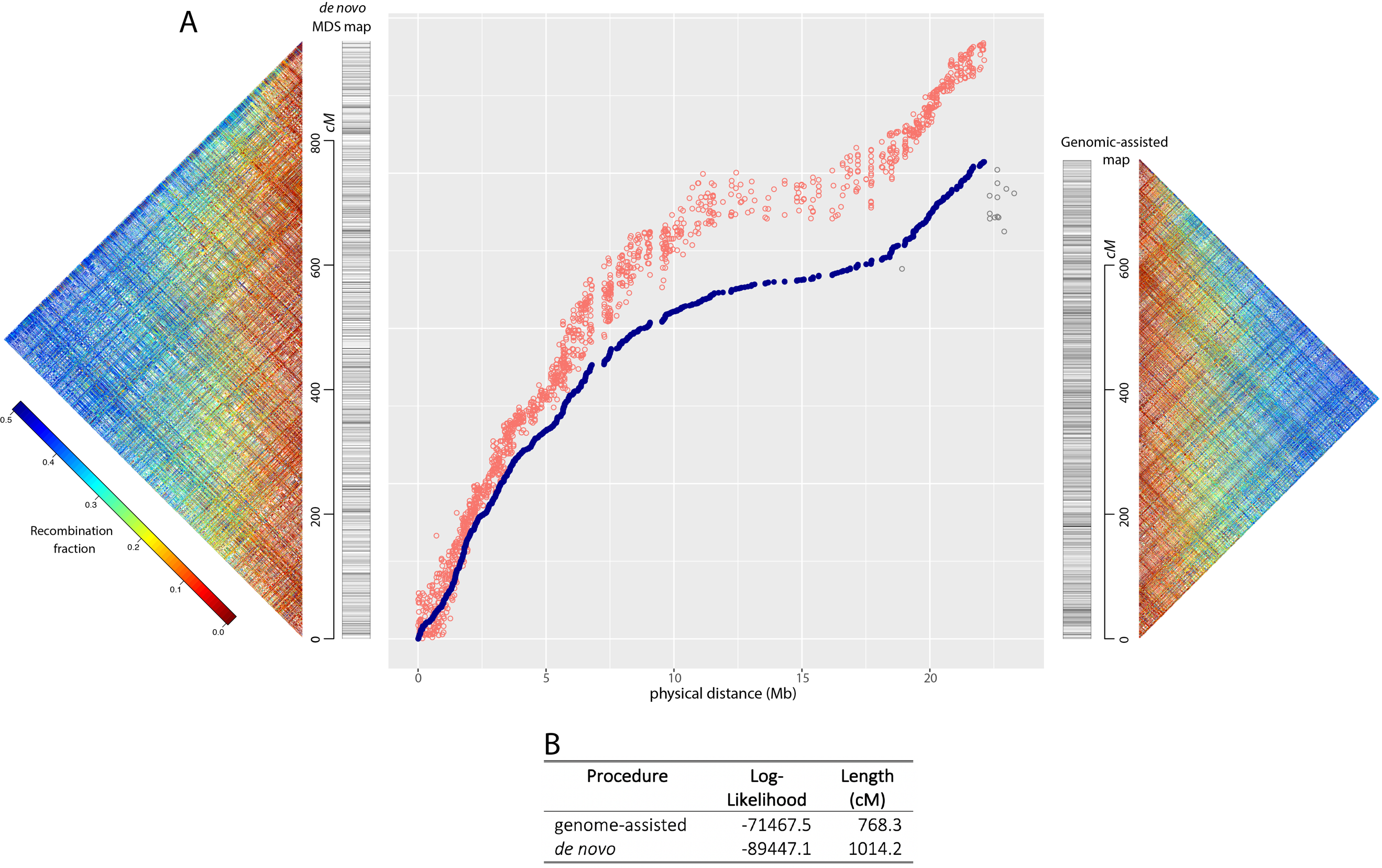
